# Neuronal RNA granules are ribosome complexes stalled at the pre-translocation state

**DOI:** 10.1101/2022.01.31.478155

**Authors:** Kalle Kipper, Abbas Mansour, Arto Pulk

## Abstract

The polarized cell morphology of neurons dictates many neuronal processes, including the axodendridic transport of specific mRNAs and subsequent translation. mRNAs together with ribosomes and RNA-binding proteins form RNA granules that are targeted to axodendrites for localized translation in neurons. It has been established that localized protein synthesis in neurons is essential for long-term memory formation, synaptic plasticity, and neurodegeneration. We have used proteomics and electron microscopy to characterize neuronal RNA granules (nRNAg) isolated from rat brain tissues or human neuroblastoma. We show that ribosome containing RNA granules are morula-like structures when visualized by electron microscopy. Crosslinking-coupled mass-spectrometry identified potential G3BP2 binding site on the ribosome near the eIF3d-binding site on the 40S ribosomal subunit. We used cryo-EM to resolve the structure of the ribosome-component of nRNAg. The cryo-EM reveals that ribosomes in the nRNAg are stalled at the elongation state where tRNA’s are in the hybrid A/P and P/E site, and resemble the pre-translocation state ribosomes. We also describe a new kind of principal motion of the ribosome, which we call the rocking motion.

## Introduction

Mammalian neurons are the largest cells in the body and are highly compartmentalized. In a highly elongated neuron, these compartments may be tens to hundreds of micrometers apart. Due to the kinetic constraints of long-distance intracellular transport, even when active mechanisms are used, these subcellular compartments must be able to function largely independently, necessitating separate pre-allocated pools of macromolecules. One important effect of such a functional separation is neuronal protein synthesis. While the bulk of neuronal protein synthesis occurs in the soma, ribosomes in the axo-dendritic compartments translate specific mRNAs in a precise spatiotemporal manner to ensure a rapid response to signals from neighboring cells in a neural circuit. This localized protein synthesis in axons and dendrites has attracted considerable attention in recent years, due to its link to synaptic plasticity and thereby long-term memory formation. Translational remodeling of gene expression has been found to underly behavioral plasticity (Sutton and Schuman, 2006), a local translational program directed by specialized ribosomes, required for synaptic plasticity and memory formation.

The ribosome follows the genetic instructions to produce proteins in all organisms. Actively translating ribosomes form polyribosomes where multiple ribosomes occupy a single mRNA molecule. Polyribosome complexes are abundant in neuronal soma but have also been detected in dendrites (Steward & Falk, 1986; Steward & Levy, 1982) and in axons (Hafner *et al*, 2019; Koenig *et al*, 2000; Li *et al*, 2005; Ostroff *et al*, 2019; Shigeoka *et al*, 2016; Sotelo *et al*, 1999; Spencer *et al*, 2000; Steward & Ribak, 1986). A prerequisite for such compartment-specific mRNA translation is the transport and storage of ribosomes to distant locations in a neuron. Active protein synthesis in the pre- and postsynaptic compartments of rodent brain has been monitored recently (Glock *et al*, 2021; Hafner *et al*., 2019). Synaptic receptors DCC and Nrp1 in *Xenopus laevis* embryonic brains interact with the ribosomes (Koppers *et al*, 2019). They also identified ∼ 48 RBPs that mediate the ribosome-mRNA association to the receptors (Koppers *et al*., 2019). Numerous studies indicate that neuronal mRNAs are transported to and stored in dendrites as stably paused (stalled) polyribosomes (Buxbaum *et al*, 2014; Dynes & Steward, 2012; El Fatimy *et al*, 2016; Graber *et al*, 2017; Graber *et al*, 2013; Krichevsky & Kosik, 2001; Langille *et al*, 2019; Lebeau *et al*, 2011; Shiina *et al*, 2005). Since kind of mechanism can bypass the translation initiation process that is the rate limiting step for protein synthesis (Graber *et al*., 2013). Upon synaptic stimulation, translation of the stored mRNA resumes (Buxbaum *et al*., 2014; Graber *et al*., 2017; Graber *et al*., 2013; Krichevsky & Kosik, 2001; Shiina *et al*., 2005). Transport of mRNA to distant locations within a neuron followed by local translation is more cost-efficient than transportation and storage of proteins (Spaulding & Burgess, 2017). In growing neurons, mRNA’s are targeted to neuronal processes as granules which also contain ribosomal subunits and translation factors (Knowles *et al*, 1996; Olink-Coux & Hollenbeck, 1996; Spaulding & Burgess, 2017). mRNAs together with ribosomes and RNA-binding proteins (RBP) form RNA granules that are targeted to axodendrites for localized translation in neurons (Buxbaum *et al*., 2014; Chu *et al*, 2019; El Fatimy *et al*., 2016; Graber *et al*., 2013; Liao *et al*, 2019; Ohashi & Shiina, 2020; Ostroff *et al*., 2019; Sahoo *et al*, 2018). Recently it was proposed that the eIF4G microexons function as a translational brake by causing ribosome stalling in cooperation with RBPs to propagate the cytoplasmic granule formation, and that misregulation of eIF4G microexons causes autism spectrum disorder (Gonatopoulos-Pournatzis *et al*, 2020). The RNA granule components are associated with various neurodegenerative disease, including autism spectrum disorder, multiple sclerosis, amyotrophic lateral sclerosis (ALS), and frontotemporal dementia (Alami *et al*, 2014; Bassell & Warren, 2008; Bentmann *et al*, 2013; Bosco *et al*, 2010; Bramham & Wells, 2007;

Broadbelt *et al*, 2006; Burguete *et al*, 2015; Conicella *et al*, 2016; Darnell *et al*, 2011; DeJesus-Hernandez *et al*, 2011; Dictenberg *et al*, 2008; Dobra *et al*, 2018; Liu-Yesucevitz *et al*, 2011; Nakayama *et al*, 2017; Russo *et al*, 2017; Salapa *et al*, 2020). Also, upregulation of axonal ribosomes occurs early in the pathogenesis of ALS (Verheijen et al., 2014).

Localized protein synthesis in neurons is essential for long-term memory formation, and synaptic plasticity (Alves *et al*, 2019; Chen *et al*, 2017; Costa-Mattioli *et al*, 2009; Graber *et al*., 2017; Lebeau *et al*., 2011; Miyakawa *et al*, 2001; Nakayama *et al*., 2017; Ohashi *et al*, 2016; Shepherd *et al*, 2006; Sudhakaran *et al*, 2014; Weng *et al*, 2018). Neuronal RNA granules serve as the functional units for transport and translational control (Alami *et al*., 2014; Gopal *et al*, 2017; Liao *et al*., 2019; Sahoo *et al*., 2018). The translational activity within the RNA granules is suppressed during their transport from soma and storage in axodendrites by specific RBPs. For example, it has been shown that G3BP1 binds to Nrn1 and Impβ1 mRNAs and attenuates their translation in axons (Sahoo *et al*., 2018). It is thought that upon synaptic stimulation, RBP-mediated repression of the granules is relieved by the dissociation of the RBP’s and translation follows of the synapse-specific mRNA’s. Targeting translationally silenced ribosomes in the form of RNA granules into the synaptic compartments could be nature’s way of regulating neuronal functions.

Although synapse-specific mRNAs are ubiquitous components of RNA granules, the number and identities of mRNAs within the granules are unclear. A transcriptome analysis of the rat hippocampus revealed 2550 transcripts localized within dendrites and/or axons (Cajigas *et al*, 2012) or 800 dominant mRNA’s in neurophiles (Glock *et al*., 2021). While several different mRNAs can be carried by the same RNA granule (Kiebler *et al*, 1999), recent quantitative imaging approaches in neurons indicate that some mRNA species can transit in granules harboring just a single mRNA species (Batish *et al*, 2012; Buxbaum *et al*., 2014; Mateu-Regue *et al*, 2019; Mikl *et al*, 2011; Park *et al*, 2014; Sahoo *et al*., 2018). This indicates that there are multiple species of RNA granules, each containing distinct populations of mRNAs and RBP’s (Elvira *et al*, 2006; Kanai *et al*, 2004). Ribosome profiling of RNA granules extracted from rat brains revealed that the most abundant mRNAs within the granules code for cytoskeletal proteins that are expressed developmentally in neuronal projection and synaptic compartments (Anadolu *et al*, 2021). Also, significant over-representation was seen of mRNA’s encoding RNA binding proteins or proteins involved in RNA metabolism (Anadolu *et al*., 2021).

Local translation requires post-translational modifications (PTM) of the bound RBP, including (de)phosphorylation and methylation that are mediated by locally-hosted enzymes in response to intracellular signaling (Coffee *et al*, 2012; Jobert *et al*, 2009; Narayanan *et al*, 2007; Urbanska *et al*, 2017). Therefore, after depolarization, localized mRNAs, including those involved in plasticity, rapidly shift from the RNA granule state to polysomes (Krichevsky & Kosik, 2001), from an inactive granule state to a translationally active polysome state.

Here we use proteomics and electron microscopy approaches to characterize molecular mechanisms of neuronal RNA granules isolated from Wistar rat brain tissues or a human neuroblastoma cell line. Proteomics detected the presence of important RBP’s known to be part of nRNAg (Caprin-1, FMRP, Staufen 1 or 2, G3BP1 or 2, hnRNP’s, ELAV-like proteins, Serbp1, and others) (Aschrafi *et al*, 2005; Chu *et al*., 2019; El Fatimy *et al*., 2016; Elvira *et al*., 2006; Krichevsky & Kosik, 2001; Nakayama *et al*., 2017). Chemical cross-linking experiments conducted with purified nRNAg samples revealed a potential G3BP2 binding site on the ribosome. Post-translational modification analysis of nRNAg shows the presence of phosphorylation and methyl-arginine sites on G3BP1 or G3BP2. The comparison of PTMs of G3BP1 or G3BP2 in nRNAg sample vs. nRNAg unbound sample, revealed differences in the modification patterns that indicate functional significance for identified PTMs for nRNAg assembly or disassembly.

The integrity and gross morphology of the purified nRNAgs was evaluated by negative staining electron microscopy (ns-EM). Differently from polysomes that tend to exhibit circular and zigzag-like morphologies (Myasnikov et al., 2014), RNA granules are known to possess a characteristic morula-like structure where individual ribosomes are tighly packed against each other (Elvira et al., 2006; Krichevsky and Kosik, 2001; El Fatimy et al., 2016). We observed a similar morula-like morphology in the cortex-hippocampus-derived nRNAg’s. and the presence of microtubules/filaments or vesicles in the proximity of nRNAg’s, indicating their involvement in the active transport of RNA granules.

The structure of the ribosomes within the RNA granule was resolved by cryo-EM, using either intact nRNAgs or individual ribosomes obtained upon an RNase T1 treatment of the granules. Based on previous studies, ribosomes form the major structural unit of the nRNAg’s (El Fatimy *et al*. 2016). The RNase T1 treated nRNAg sample was resolved to a 2.9 Å resolution whereas the intact map was resolved to 5.5 Å. We find that ribosomes are stalled in polysomic state in the nRNAg particles. This finding is consistent with Graber et al, 2014 where the authors used chemical ribopuromycylation assay and concluded that synaptic mRNAs are transported as stably paused polyribosomes. Recent ribosome profiling results of nRNAg support the polysome stalling as well (Anadolu *et al*., 2021). We show the structural proof that ribosomes in nRNAg’s are mainly stalled at polysomic state. The most suprising finding of our cryo-EM data is the discovery of new kind of principal motion inside the ribosome that we call “rocking” motion, where ribosomal small subunit moves around the horizontal axis.

## Results

### Neuronal RNA granules are morula-like particles

Wistar rat cerebral cortex, cerebellum, and hippocampus were used as sources of RNA granule material, as those tissues are involved in various memory-related tasks (Alves *et al*., 2019; Chen *et al*., 2017; Costa-Mattioli *et al*., 2009; Graber *et al*., 2017; Lebeau *et al*., 2011; Miyakawa *et al*., 2001; Nakayama *et al*., 2017; Ohashi *et al*., 2016; Shepherd *et al*., 2006). To reduce contamination from the surrounding glial cells, the tissues were extracted from the brains of postnatal day 0 to 6 rats of both sexes, as gliogenesis starts from postnatal week 2 (Bandeira *et al*, 2009). RNA granules from tissue homogenates were purified using a combination of ultracentrifugation and size-exclusion chromatography. Briefly, the cytoplasmic extracts of the cortex-hippocampus tissue were first centrifuged through a 60 % sucrose cushion at 246 000 × g, followed by the solubilization of the resulting opalescent pellet (hereafter referred to as “Pellet 1”). According to a negative staining EM analysis, this material was heterogenous, containing filamentous polysome-like structures, as well as tightly packed, round-shaped particles of 100 – 200 nm diameter (Supplementary Figure S1A). The overall size and shape of the spherical-shaped particles resembles density-gradient purified RNA granules (Anadolu *et al*., 2021; El Fatimy *et al*., 2016; Krichevsky & Kosik, 2001).

To obtain a more homogenous preparation of the RNA granules, the redissolved sucrose gradient pellet fraction was passed through a Sephacryl S500 size-exclusion resin. The gel filtration technique has been used in the past to isolate Staufen-containing ribonucleoprotein particles from rat brain (Mallardo *et al*, 2003). Upon size-exclusion chromatography, 20 % - 50 % of the original pelleted material was recovered in the flow-through (referred to as “nRNAg”). EM analysis of the Sephacryl S500-purified material revealed a near-homogenous population of the spherical-shaped particles seen in the original sucrose gradient pellet (Figure 1 or Supplementary Figure S1B). When subjected to treatment with different RNases (RNase T1, MNase) at a physiological potassium concentration (150 mM), a majority of the morula-like particles in an nRNAg preparation falls apart into what appear as monosomes or smaller polysomes (di- or trisomes) (Figure 2; Supplementary Figure S2). However, depending on the type of nuclease, a variable fraction of the large and compact particles persist even after the nuclease treatment, particularly with MNase (Supplementary Figure S2A).

**Figure 1.**
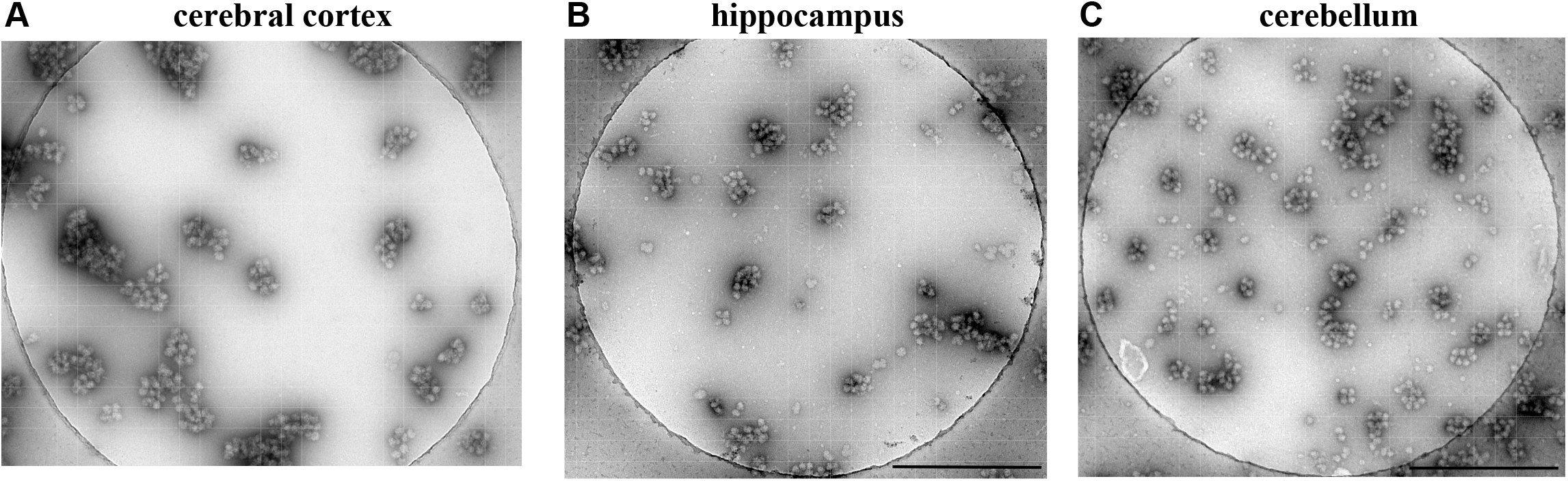
Neuronal RNA granules resemble morula like compact particles. Neuronal RNA granules purified from different rat brain tissues are visualized with negative stain TEM. RNA granules purified from rat P0 brain tissues extracted from the: (A) cerebral cortex, (B) hippocampus, or (C) cerebellum. The scale bar on the bottom corner represents 500 nm distance.

**Figure 2.**
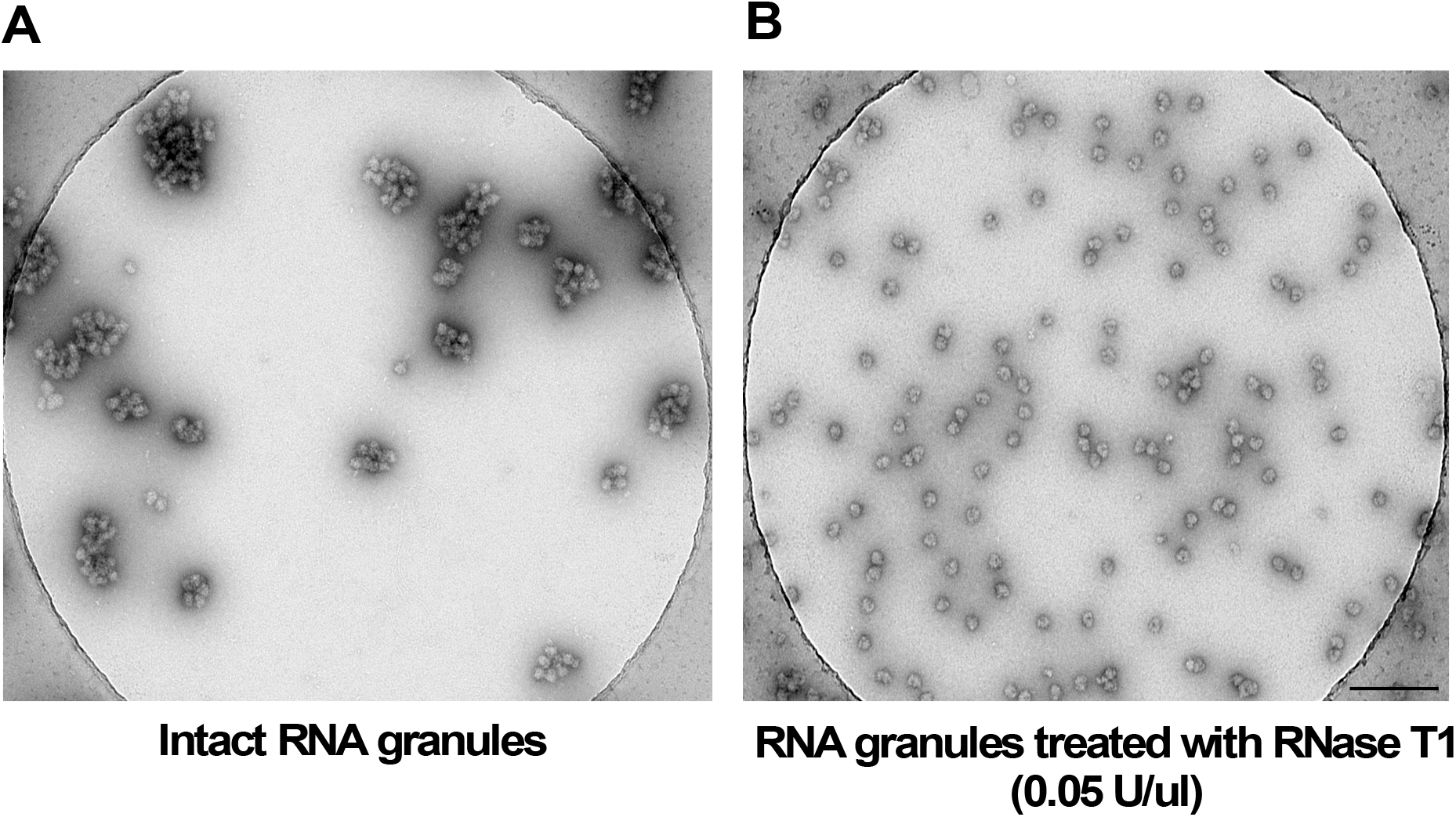
Negative stain TEM images of RNase T1 treatment of neuronal RNA granules. (A) RNA granules purified from P3 rat cerebral cortices. (B) The same nRNAg sample after treatment with 1U RNase T1 for 30 min at room temperature. After RNase T1 treatment, tightly packed granules fall-apart into mono/disomes. The scale bar on the bottom corner represents 200 nm distance.

The same purification protocol was used to obtain nRNAg’s from hippocampus (P0 to P5 rats) and cerebellum (P6 to P7 rats). The overall morphology of nRNAgs purified from those tissues was similar to the cortex-derived granules (Figure 1B and 1C). Consistent with the smaller number of neurons in hippocampus and cerebellum (Bandeira *et al*., 2009), the yield of nRNAgs was reduced 10-fold for the cerebellum and 40-fold for the hippocampus relative to the cortex. Since the small amount of material obtained from hippocampus/cerebellum was insufficient for the downstream biochemical analyses, the experiments described below were exclusively performed with the the cortex-hippocampus (CH)-derived nRNAgs.

To make sure that the tightly packed particles in the nRNAg preparation are not an artifact of polysome aggregation due to the pelleting of the particles under high g-forces, we omitted the pelleting of the cortex-hippocampus postmitochondrial lysate and analyzed the sample for the presence of the morula-like particles. In the first experiment, the clarified lysate was directly passed through the Sephacryl-S500 resin and the particles were analyzed by ns-EM. Though larger contaminating complexes (vesicles, membranes) remain in the sample, a few tightly packed particles similar in size and shape to the nRNAg’s were still observed (Supplementary Figure S3A – S3C). In the second experiment, the clarified lysate was centrifuged over a double sucrose cushion (90% bottom/ 60% above), so that the nRNAg material remained in the interface of two sucrose cushions (hence no pelleting of the particles) and was subsequently concentrated in a stirred cell unit where particles remain in solution during the concentration. Again, particles similar to the nRNAg’s fraction were observed in an ns-EM analysis (Supplementary Figure S3D). Collectively, these observations indicate that the tightly packed complexes in the nRNAg fraction are not pelleting artifacts, but rather represent *bona fide* RNA granules.

Stress-granules (SG) are another type of RNA-protein complexes, whose assembly is similar to neuronal RNA granules in that it requires RBPs containing low complexity (LC) and intrinsically disordered (ID) regions (Kedersha *et al*, 2013). Unlike nRNAg, stress granules are phase-separated liquid-like droplets exhibiting the macroscopic behavior of hydrogels (Gopal *et al*., 2017; Langdon & Gladfelter, 2018; Shiina, 2019). We observe such liquid-like droplets, when purified recombinant G3BP1 is visualized by ns-EM without additional factors (Supplementary Figure S4A and B). No droplets were observed with an MBP-tagged G3BP1 (Supplementary Figure S4C). MBP-tag is known to keep proteins in their solublized state (Costa *et al*, 2014). These liquid-like droplets of G3BP1 are markedly different from the solid morula-like particles that we obtain from the brain tissue. Additionally, the morula-like particles persist when the extraction and purification of the RNA granules is performed in the presence of cycloheximide, a known inhibitor of stress granule assembly (Bounedjah *et al*, 2014; Buchan *et al*, 2011; Kedersha *et al*, 2000). Below we present mass-spectrometry data for the presence of a full set of r-proteins (including the proteins of the 60S subunit) in the RNA granules as a further argument against the morula-like particles being stress granules, as 60S subunits are excluded form SG (Kimball *et al*, 2003; Wolozin & Ivanov, 2019). Coupled with the fact that the processing of the cryo-EM data (2D and 3D classification) revealed the 80S ribosome as the prevalent component of the granules, these observations collectively indicate that the morula-like particles in the nRNAg fraction are unlikely to be stress granules and rather represent *bona fide* neuronal RNA granules, as observed previously (Anadolu *et al*., 2021; El Fatimy *et al*., 2016; Krichevsky & Kosik, 2001).

### Proteomic analysis of nRNAg composition

An LC-MS/MS analysis of the cortex-derived nRNAg preparation revealed the presence of a full set of ribosomal proteins, as well as some r-protein analogs. The r-proteins were the most abundant identified proteins based on their iBAQ (intensity based absolute quantification) value (Supplementary Table S1). We also identified some uncharacterized r-protein analogs in the nRNAg preparation (Supplementary Table S2). We estimated the relative abundance of the identified r-proteins by normalizing the iBAQ value of each individual r-protein to the mean iBAQ value over all identified r-proteins. We chose the iBAQ value as the metric for r-protein abundance since it is commonly used to estimate the relative abundance of proteins from label-free proteomics data (Nagaraj *et al*, 2011; Schwanhausser *et al*, 2011) and has been shown to represent the best correlation between biological replicates, a normal distribution among all protein abundances, and the lowest variation among ribosomal protein abundances (Arike *et al*, 2012). For 66 out of the 80 identified r-proteins in the corticohippocampal nRNAg sample, the fold difference in the relative abundance with respect to the mean did not exceed 2 (Supplementary Table S3 and Supplementary Figure S5). A reanalysis of previously published data on r-protein abundance from mouse embryonic stem cells (Supplementary Figure S5) (Slavov *et al*, 2015) and HEK293-6E cells (Supplementary Figure S6) (van de Waterbeemd *et al*, 2018) revealed a similar “within 2-fold” variation in the relative abundances of the r-proteins. Out of the remaining 14 r-proteins in the nRNAg, the relative abundances of 12 r-proteins were 2 to 11-fold below the mean while the abundance of ubiquitin-60S ribosomal protein L40 (Uba52) was 97-fold below the mean (Supplementary Table 3). With the exception of Rpl37 and Uba52, the low-abundance r-proteins tended to be localized on the ribosomal surface and may therefore have been partially lost during nRNAg isolation. We also note that the number of identified unique + razor peptides was lower for the low-abundance proteins, particularly for Rps27l, Rpl37 (7 peptides) and Rpl39 and Uba52 (3 peptides) (Supplementary Table 3).

The RBPs (Caprin-1, G3BP1 or 2, Staufen2, Serbp1 etc.) were the second class of proteins that were identified in the RNAg preparations by the MS (Supplementary Table S1) or Western blot analysis (Supplementary Figure S7). To estimate the stoichiometry of RBPs relative to the ribosome in the RNAg we used the iBAQ intensity values determined by/obtained from the proteomic analysis of RNA granules. We first calculated the ribosome iBAQ value as an average of the individual iBAQ values of the r-proteins in two biological replicates, see Material and Methods (Supplementary Table S1).This average iBAQ intensity value of r-proteins was taken as 1 equimole of ribosome and the iBAQ values of the RBPs were normalized to it. Based on these estimations, the relative abundance of key RBPs varies from 8 to 123 fold compared to ribosomes (Supplementary Table S4) with Caprin-1 and G3BP2 being the most abundant ones at 9.5 and 8.4 ribosomes per RBP, respectively.

### Neuronal RNA granules in a human neuroblastoma cell line

Human neuroblastoma cell line SH-SY5Y is a widely used in vitro model for various neurodegenerative disorders, as SH-SY5Y can be induced to differentiate into neuron-like cells by treatment with differentiation-promoting agents, including all-trans-retinoic acid (ATRA) (Kovalevich & Langford, 2013). We cultivated SH-SY5Y cells in the presence of 10 μM ATRA for 5 days, followed by cell lysis and purification of the post-mitochondrial supernatant identically to the cortical material. An ns-EM analysis of the Sephacryl-S500 purified material from SH-SY5Y revealed the presence of occasional morula-like particles similar to the particles obtained from the rodent cerebral cortex (Figure 3). A full set of r-proteins was detected in the nRNAg preparation in an MS analysis (Supplementary Table S1). Overall, the relative abundances of r-proteins in the RNAgs from rat and SH-SY5Y were significantly correlated (Pearson corrleation coefficient 0.59) (Supplementary Figure S8A). We also noted a significant positive correlation between the r-protein abundances in the SH-SY5Y RNA granules and 80S ribosomes from HEK293-6E (correlation coefficient 0.55) (Supplementary Figure S8C). Moreover, out of the 18 low-abundance r-proteins in the SH-SY5Y RNA granules, 7 proteins (Rps27a, Rpl10a, Rpl37, Rps26, Rps15, Uba52, and Rps27l) were shared with the low-abundance r-protein set of the rat RNA granules (Supplementary Table S3). Interestingly, two of the low-abundance r-proteins in rat nRNAg (Rplp2, Rpl35) were in contrast present in the high-abundance r-protein set (r-proteins with a relative abundance more than 2-fold higher relative to the mean) in the SH-SY5Y RNA granules (Supplementary Table S3).

**Figure 3.**
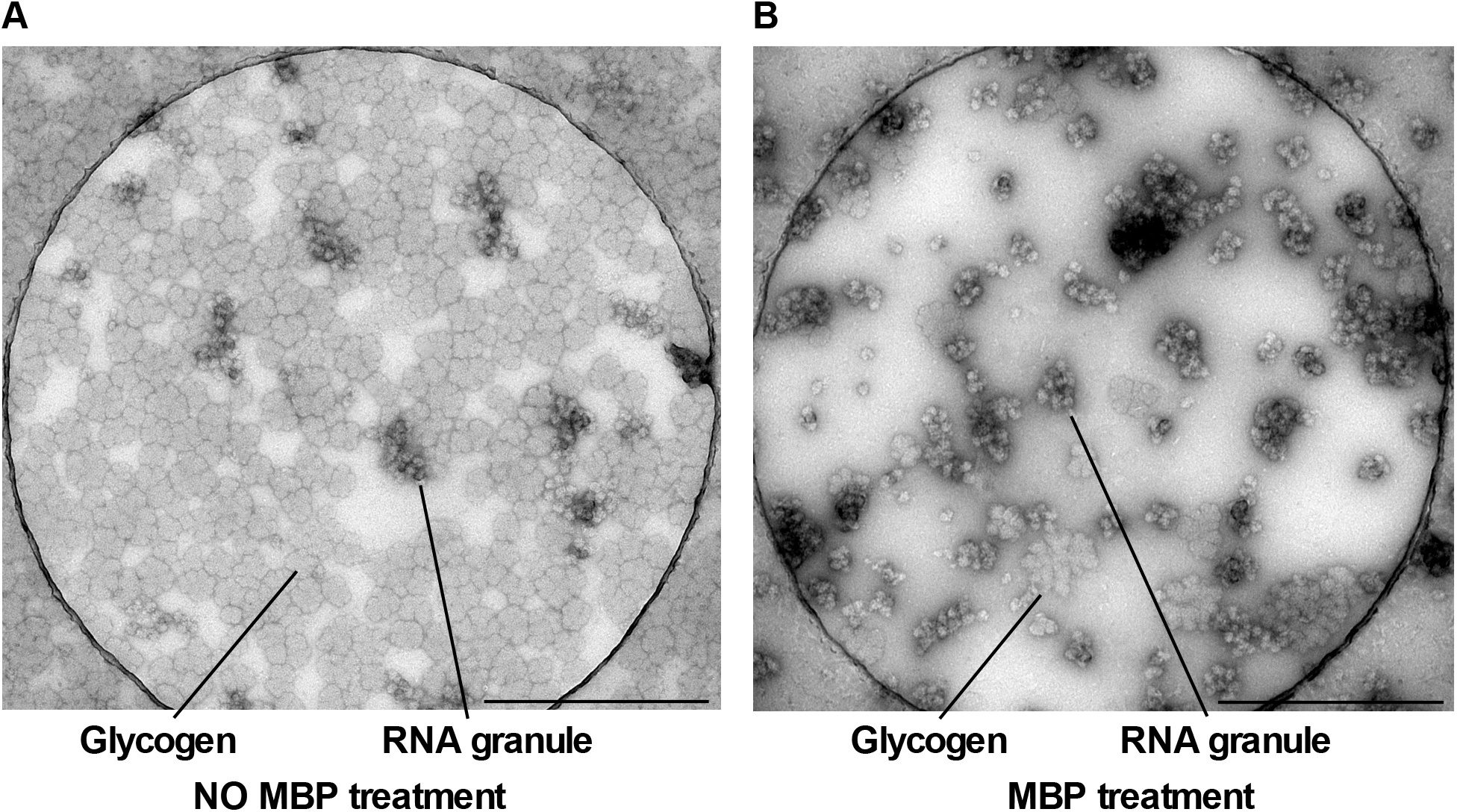
Negative stain TEM images of RNA granules purified from ATRA-differentiated human neuroblastoma cell line SH-SY5Y. (A) RNA granule preparation after centrifugation through sucrose cushion followed by size-exclusion chromatography. Co-purifying glucogen granules are the major species in the sample at this stage, with only occasional nRNAg visible. (B) the contaminating glycogen granules have been removed by affinity chromatography on an MBP-coated Ni-Sepharose resin while nRNAg’s remain in the sample. The scale bar on the bottom corner represents 500 nm distance.

Similarly to the rat corticohippocampal nRNAgs, we estimated the relative abundance of RBPs in the SH-SY5Y derived nRNAg. Though the variation of the relative abundance of the RBPs with respect to ribosomis higher between the two replicates of the SH-SY5Y RNA granules the overall trend is that there is lesser amount of RBPs per ribosome in the SH-SY5Y nRNAg compared to the corticohippocampal RNA granules. Caprin-1 together with ELAVL4 and Serbp1 were the most abundant RBPs in the SH-SY5Y RNA granules, with a stoichiometry of one molecule of the respective RBP per 18.6, 7.8 and 8.1 ribosomes (Supplementary Table S4).

However, as shown by the ns-EM analysis, the major component of the nRNAg preparation consisted of lighter-staining particles, which bear close resemblance to glycogen storage granules (Figure 3A) (Besford *et al*, 2012; Sullivan *et al*, 2010). The levels of glycogen are up-regulated in various cancers: breast, kidney, uterus, bladder, ovary, skin and brain cancer cell lines (Zois *et al*, 2014).

We tried to remove the contaminating glycogen granules from the nRNAg preparation by subtractive affinity chromatography on a Ni-Sepharose resin pre-coated with a His-tagged maltose-binding protein (His-MBP) as MBP is known to bind various oligosaccharides (Quiocho *et al*, 1997; Walker *et al*, 2010). The unbound fraction (i.e. material not bound to MBP-Ni-Sepharose) obtained after pelleting the Ni-Sepharose resin consisted mostly of the morula-like stained particles similar to the ones observed in the cortical nRNAg sample (Figure 3B), indicating the ability of MBP to remove the glycogen contamination. However, a potential drawback of the Ni-Sepharose-coupled purification scheme is the ability of the Ni-resin to nonspecifically bind poly(A) containing RNA sequences (Nastasijevic *et al*, 2008). Since the majority of eukaryotic mRNAs contain poly(A) tails, we anticipated that at least a fraction of the mRNA-containing RNA granules may be selectively retained on the resin, thus biasing the downstream biochemical and cryo-EM analysis of the RNA granules. Indeed, when the His-MBP coated Ni-Sepharose resin was incubated with a cortical RNA granule preparation, followed by a wash-out of the resin with buffer and a subsequent elution of the nonspecifically bound material in 8 M urea, a mass-spectrometry analysis of the urea eluate detected 127 proteins, most of them r-proteins, as well as a number of RBP components of RNA granules (Caprin-1, Hnrnpu, Hnrnph1, Hnrnpd, Hnrnpr, Hnrnpf, Ybx1, Npm1, Serbp1, G3BP1, G3BP2, PABPC1, Ncl, Celf2, Cirbp, TDP-43, Elavl2, Stau2) (Supplementary Table S5). The non-specifically bound material was also tested against the anti-eL8, anti-G3BP1 and anti-Caprin-1 antibodies by Western blot analysis (Supplementary Figure S9) that shows the presence of key RBPs. Based on this analysis we conclude that extraction of RNA granules from the neuroblastoma cell line requires further optimization with respect to removing the extensive glycogen contamination, while simultaneously preserving the compositional integrity of the RNA granules. Hence, in the following we restrict our analysis to the brain-derived RNA granules.

### nRNAg active transport by microtubules or filaments

In a ns-EM analysis of the cortical RNA granules we occasionally observe very large nRNAg that overstain, but have clearly visible and abundant microtubule/actin filaments associated with them (Figure 4). To our knowledge this is the first time that such a proximity between the nRNAg’s and components of the cytoskeleton has been captured by EM. This result supports the notion that nRNAgs are actively transported on the microtubule/actin filaments (Balasanyan & Arnold, 2014; Heisler *et al*, 2011; Hirokawa, 2006). The association of nRNAgs with the components of the cytoskeleton is supported by our proteomic analysis of nRNAg, where we observe a number of microtubule related proteins in the cortical nRNAg preparation, including Tubb4b, Map1b, Tubb2b, Tubb3, Map1a, Mapt, Tubb2a, Tuba1b/Tuba4a, Kif5a, Kif1c, DynII1, DynII2, Vbp1, and Map2 (Supplementary Table S1). Components of the actin filaments were also detected in the nRNAg preparation, including Actb, Actr8, Acta1, Capzb, Dctn2, Fmnl2, Fnbp1 and Capza1 (Supplementary Table S1).

**Figure 4.**
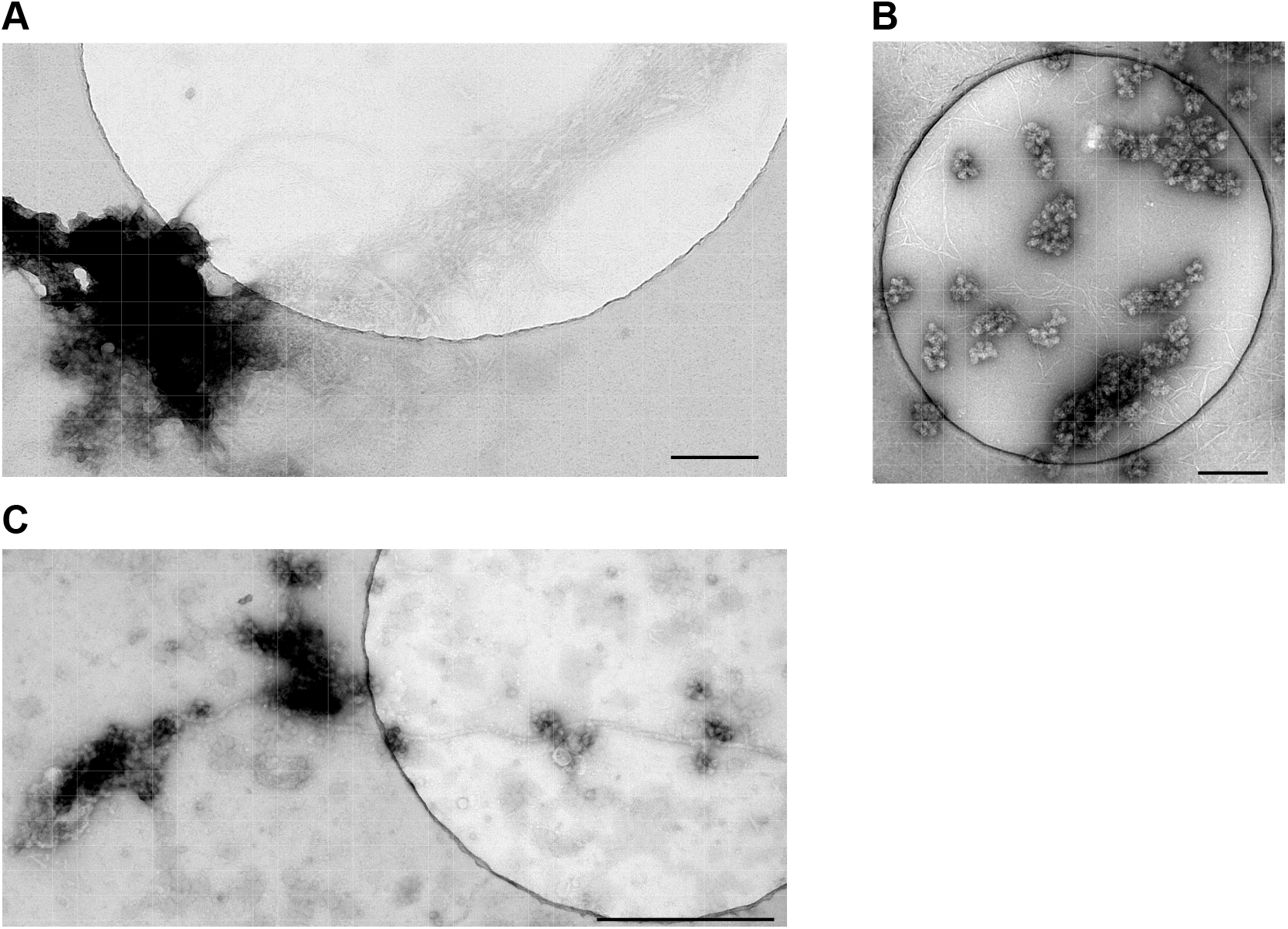
Microtubules/actin filaments are in close proximity to nRNAg’s. (A-B) nRNAg purified from rat cortices. The scale bar on the bottom corner represents 200 nm distance. (C) nRNAg purified from rat hippocampus tissue. White filaments resembling the microtubule or actin filaments are entangled with the nRNAg particles. The scale bar on the bottom corner represents 500 nm distance.

The close association of the cytoskeletal components with nRNAg’s is further supported by crosslinking-coupled mass-spectrometry (CL-MS) experiments aimed at identifying interaction networks in RNA granules. Using cross-linkers of varying spacer arm lengths (DSS, DSG, BS3, BS(PEG)5, and Leiker), we were able to identify crosslinks between microtubule components and r-proteins, translation initiation factors, as well as uncharacterized proteins (Table 1). For example, using BS(PEG)5 we observed a cross-link between residue K402 of Dctn2 and K129 of the ribosomal protein eS24. Dctn2 is involved in dynein binding to an organelle and plays a role in synapse formation during brain development (LaMonte *et al*, 2002; Urnavicius *et al*, 2015). Likewise, using DSG we observed a cross-link between residue K143 of eS19 and K2 of kinesin-like protein 1 A (Kif1A), a neuron-specific protein involved in anterograde axonal transport of synaptic vesicle precursors (Okada *et al*, 1995). Residue K724 of another kinesin isoform, 5C (Kif5C) cross-linked to residue K11 of eS19 (Table 1). However, the interactions of cytoskeletal proteins in the RNA granules are not restricted to r-proteins as shown by the BS(PEG)5-specific cross-links of microtubule associated proteins Map1b and Map2 to translation initiation factors eIF5b and eIF3a (Table 1). Additionally, the microtubule associated protein Map1a gave a cross-link to Lrrc49, an uncharacterized protein predicted to be part of neuronal tubulin polyglytamylase complex (Huttlin *et al*, 2020). Hence, our observation of a Map1b-Lrrc49 cross-link provides further evidence that Lrrc49 is indeed involved in the microtubule trafficking complex.

**Table 1.**
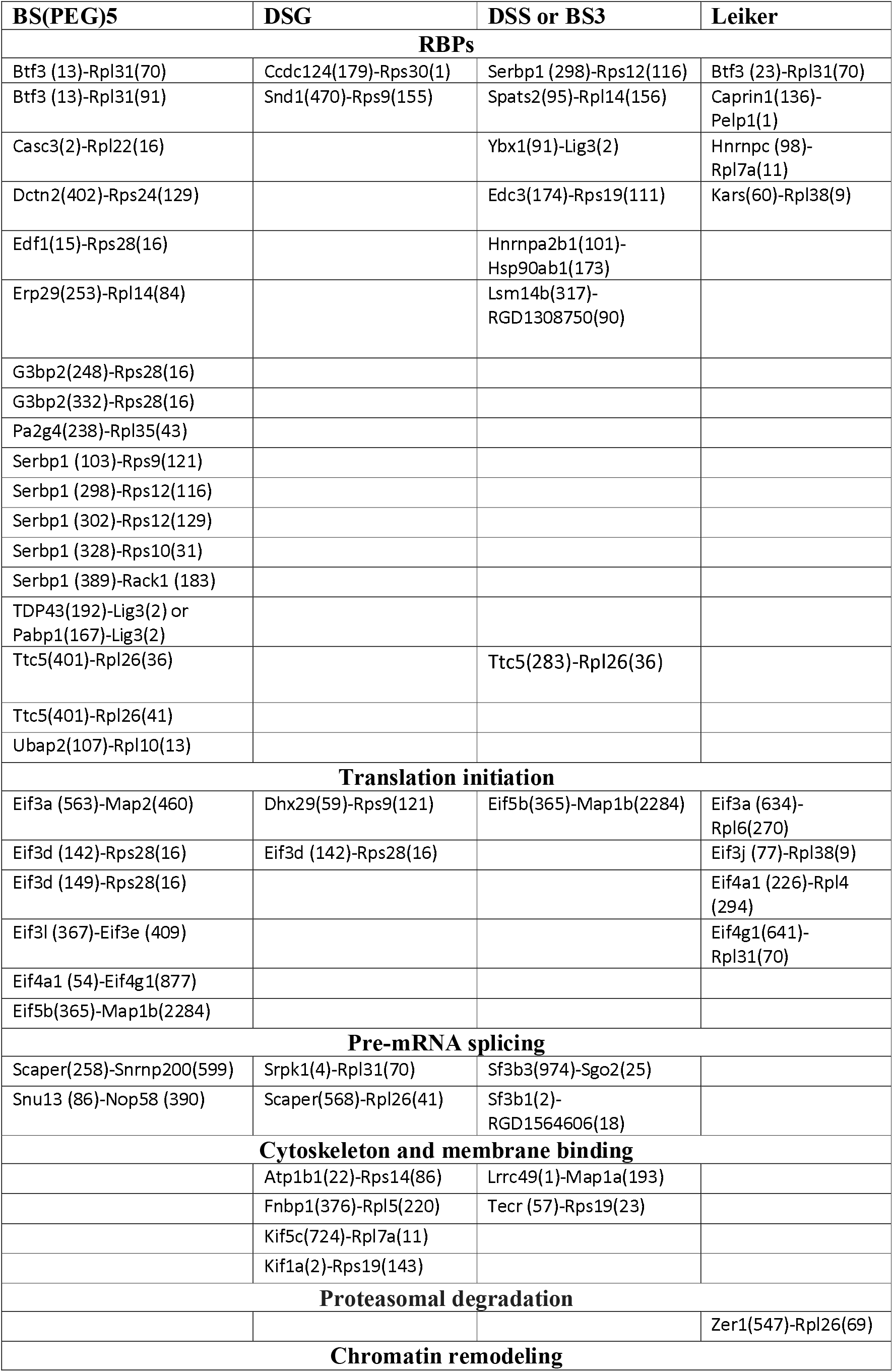
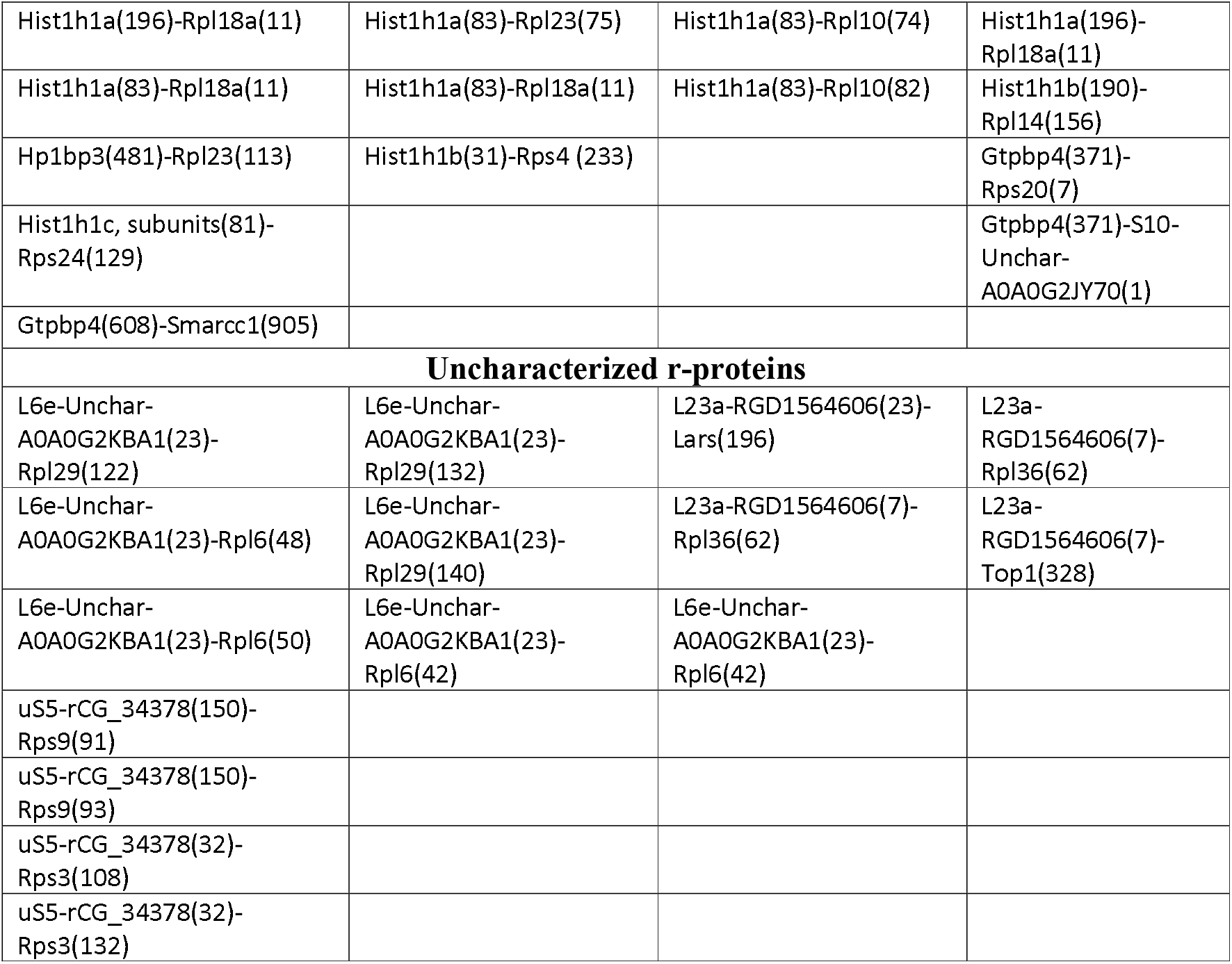
Cross-linking coupled mass-spectrometry results of nRNAg. nRNAg were purified from rat cortices and cross-linkers having various spacer arm lengths were used. Cross-linkers DSG (spacer arm length 7.7 Å), Leiker (9.3 Å), DSS or BS3 (11.4 Å), and BS(PEG)_5_ (21.7 Å) are indicated on top row and CL proteins are sorted by functional groups. Only intermolecular CL are listed in the table. All the rest of the identified peptides are listed in Supplementary Table S5. pLink software was used to process the mass-spectrometry CL data (Yang *et al*, 2012).

In addition to the microtubule/filament components, the RNA-binding protein Rtraf (hCLE) was detected in nRNAg’s in our proteomic analysis. In mature brain, Rtraf is found in mRNA-containing kinesin-associated granules in dendrites (Kanai *et al*., 2004) whereas in developing brain Rtraf is a component of cytosolic, ribosone-containing RNA granules that transport specific mRNAs to dendrites for local translation (Elvira *et al*., 2006). Recent evidence indicates that Rtraf in complex with Ddx1, Fam98B and/or Hspc117 (Rtcb) forms an alternative cap-binding complex which may modulate the translation of specific mRNAs in response to particular synaptic signals (Pazo *et al*, 2019). We note in this connection that Ddx1, Rtcb and Fam98A were also detected in nRNAgs in our proteomic analysis. Together, the identification of cytoskeletal components and regulatory proteins like Rtraf in our nRNAg preparation as well as the physical association of the granules with microtubules/actin filaments seen in ns-EM supports the notion that nRNAg move along the cytoskeleton components.

### Lysosome trafficking of nRNAg

When the purified cortical nRNAg’s are analyzed by ns-EM, we occasionally observe RNA granules closely apposed to what appears as broken vesicles of a ca. 200 nm diameter (Figure 5). Those vesicles could be derived from e.g. endosomes or lysosomes. Given that our purification scheme includes an initial hypotonic lysis step and that lysosomes are known to be destroyed under hypotonic conditions, in contrast to osmotically insensitive endosomes (Schroter *et al*, 1999), the vesicle remnants are more likely to represent lysosomes. Since endocytic organelles do not penetrate dense sucrose solution during ultracentrifugation (de Araujo *et al*, 2008), the presence of the vesicle remnants in the nRNAg preparations indicates their physical association with the RNA granules prior to and during the purification. Importantly, in a recent study combining fluorescence microscopy and proximity labeling proteomics, lysosomes were shown to serve as vehicles for RNA granule transport with Anxa11 functioning as a tether between RNA granules and lysosomes (Liao *et al*., 2019). Anxa11 presence was not detected in purified nRNAg’s in this study. Instead, Anxa6 was present in our nRNAg sample and may play a similar role in granule trafficking as Anxa11. This observation fits in with a growing body of evidence indicating a central role for membrane-bound organelles in RNA granule trafficking in different cell types (Pushpalatha & Besse, 2019), including neurons, where the organelles may additionally serve as platforms for localized mRNA translation (Cioni *et al*, 2019). Importantly in this respect, our proteomic analysis identified Ras-related proteins in brain (Rab) GTPases that are molecular switches cycling between a cytosolic GDP-bound inactive form and a membrane-associated GTP-bound active form to regulate membrane trafficking (Stenmark, 2009; Tang *et al*, 2014). Different Rab’s localize to distinct membrane-bound cellular compartments in mammalian cells, including the endoplasmic reticulum (ER)-Golgi region (Rab1a, Rab2, Rab6, Rab30, and Rab33b), early and recycling endosomes (Rab4, Rab5, Rab11, Rab15, Rab17, Rab18, Rab22, and Rab25), late endosomes/lysosomes (Rab7 and Rab9), and specialized organelles such as synaptic vesicles (Rab3a and 3c), secretory granules (Rab3D, Rab37), and melanosomes (Rab27) (Junutula *et al*, 2004; Zerial & McBride, 2001). Rab18, Rab11a, Rab11b, Rab7a, Rab1a, Rab6b and Rab2a were detected in our nRNAg preprations (Supplementary Table S1). In addition, Gdi1 was detected that regulates the Rab binding to membranes through GDP/GTP exchange (Oesterlin *et al*, 2012). Our nRNAg proteomics analysis also identified uncharacterized protein (Gene ID: A0A0G2KAW3) that is believed to be part of lysosome trafficking (predicted by Uniprot). The protein sequence is 45% identical to Rab7a.

**Figure 5.**
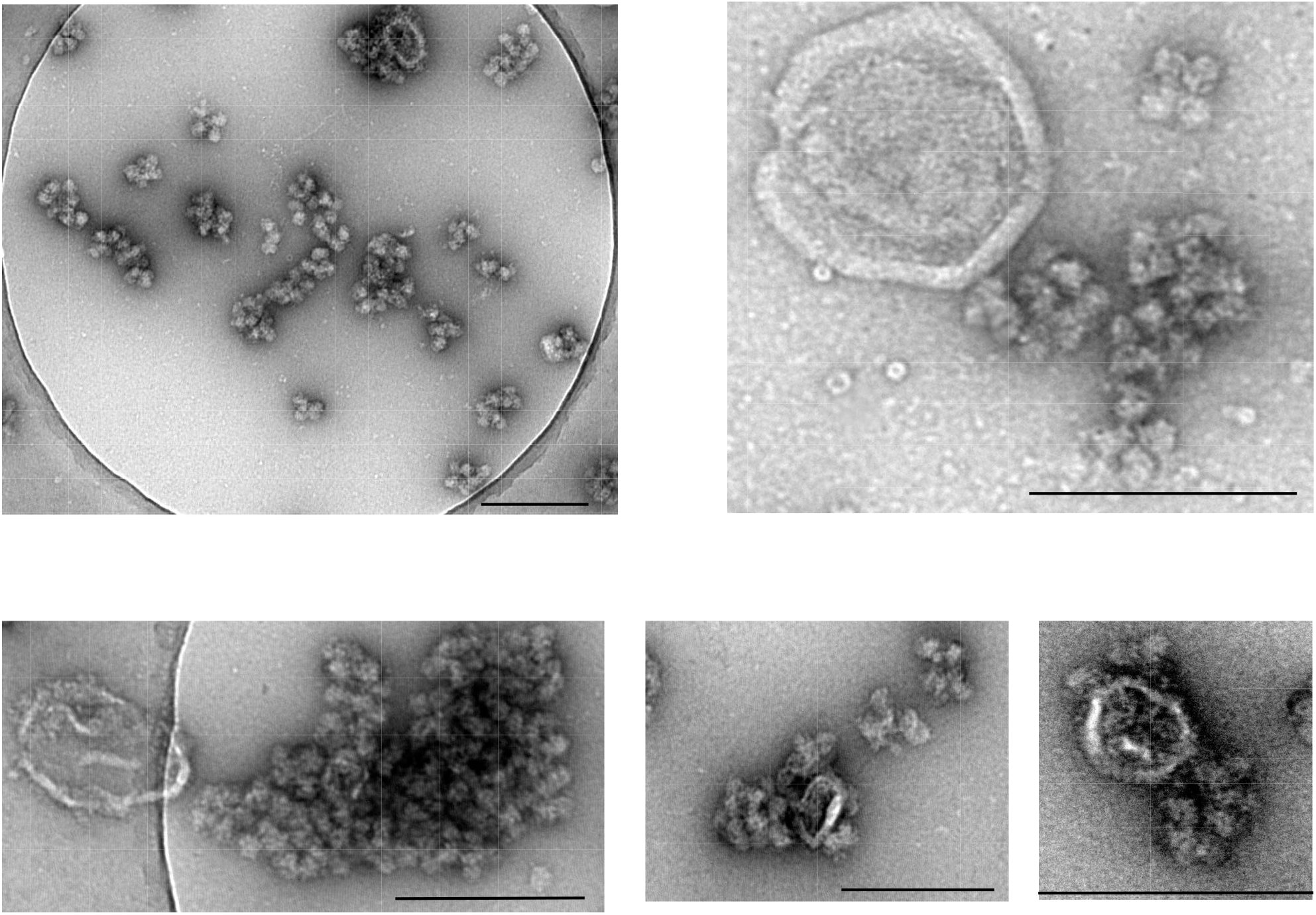
Lysosomes seen in close proximity to nRNA’s and their involvement in nRNAg trafficking. ns-EM images of nRNAg purified from rat cortices showing lysosomes/vesicles (white broken round organelles) in proximity to densely packed nRNAg’s. The scale bar on the bottom corner represents 200 nm distance.

The crosslinking-coupled proteomics study was unable to detect cross-links between lysosome components and other proteins, probably to do the low abundance of complexes that co-purify with nRNAg.

Hence, the observed nRNAg-adjacent broken vesicles may be remnants of the RNA granule trafficking machinery which co-purifies with the granules.

### RBPs interactions with the ribosome

In vivo fluorescence co-localization, together with co-sedimentation and pull-down assays, are commonly used to probe the interaction pattern of RBPs within RNA granules (Kedersha *et al*, 2016; Russo *et al*., 2017). However, given the central role of RBPs in RNA granule assembly/disassembly, those methods are not sufficient to provide detailed information about the molecular contacts involved in the modulation of the translational activity of the RNA granules by RBPs. This information can potentially be provided by cross-linking or proximity-based labelling techniques (Leitner *et al*, 2016; O’Reilly & Rappsilber, 2018). To characterize the interactions of RBPs with components of the RNA granule at a molecular level, we subjected the purified RNA granules to cross-linking with various lysine-specific cross-linkers, followed by a chromatographic enrichment of the cross-linked peptides and a mass-spectrometric identification, essentially as described by Leitner and coworkers (Leitner *et al*, 2012). As expected, the majority of the identified cross-links were between r-proteins as the ribosomes are the main component of the RNA granules (Supplementary Table S6). CL-MS study identified over 50 different intermolecular and over 200 intramolecular CL between r-proteins (rProt). The rProt -rProt crosslinks are within the distance spanned by the cross-linker BS(PEG)_5_ when mapped on the ribosome structure (5AJ0), validating our cross-linking protocol (Figure 6). However, we managed to identify a number of cross-links between known RBPs and r-proteins or other components of the translational apparatus (Table 1). The most interesting CL to us is a cross-link between G3BP2 (residues 248 or 332) and eS28 (residue 16) (Table 1). This cross-link indicates that in the cortex-derived RNA granules G3BP2 is located close to the ribosomal E-site, hinting at a direct interaction of G3BP2 with the ribosome. The Ras-GAP SH3 domain binding proteins (G3BPs) are well-known RNA-binding proteins whose medical relevance is underscored by the observation that their aberrant expression is common in various types of cancer, misregulated stress granule formation, defects in the translation of axodendritic mRNAs as well as by the fact that they are frequently targeted by viruses (Alam & Kennedy, 2019; Bidet *et al*, 2014; Reineke & Lloyd, 2013; Tourriere *et al*, 2003; White *et al*, 2007). As mammals contain two versions of G3BP (G3BP1, G3BP2), it is of note that the cross-link to eS28 was only detected with G3BP2, despite G3BP1 appearing in the list of proteins identified in our nRNAg preparations. Moreover, the same residue (K16) of eS28 that cross-linked to G3BP2 also gave a cross-link to eukaryotic initiation factor eIF3d (residues 142 and 149) (Table 1). Unfortunately, this cross-link could not be directly mapped to the structure of eIF3d in the 48S complex since the region of eIF3d containing the cross-linked residues is missing from the existing cryo-EM map (PDB id: 6fec or 6zmw) (Brito Querido *et al*, 2020; Eliseev *et al*, 2018). However, in the 48S structure of Eliseev and coworkers containing residues 172 – 536 of eIF3d the overall placement of the protein indicates that the missing region containing the cross-linking sites could indeed approach residue 16 of eS28 (Figure 7). eS28 together with eS26, and uS11 appear to interact with the eukaryotic mRNA and thus line out an alternative path of the mRNA exit (Budkevich *et al*, 2014). G3BP2 binding near the E-site or in the same pocket as eIF3d may indicate that G3BP2 interacts with the 5’-side of the mRNA that exits the ribosome. A binding location near the E-site presents clues about the mechanism of how the ribosomes could be stalled by G3BP2 since many translation inhibitors bind to 40S or 60S subunit E-site: cycloheximide, lactimidomycin, and phyllanthoside in 60S E-site or emetine, pactamycin, cryptopleurine and edeine in the 40S E-site.

**Figure 6.**
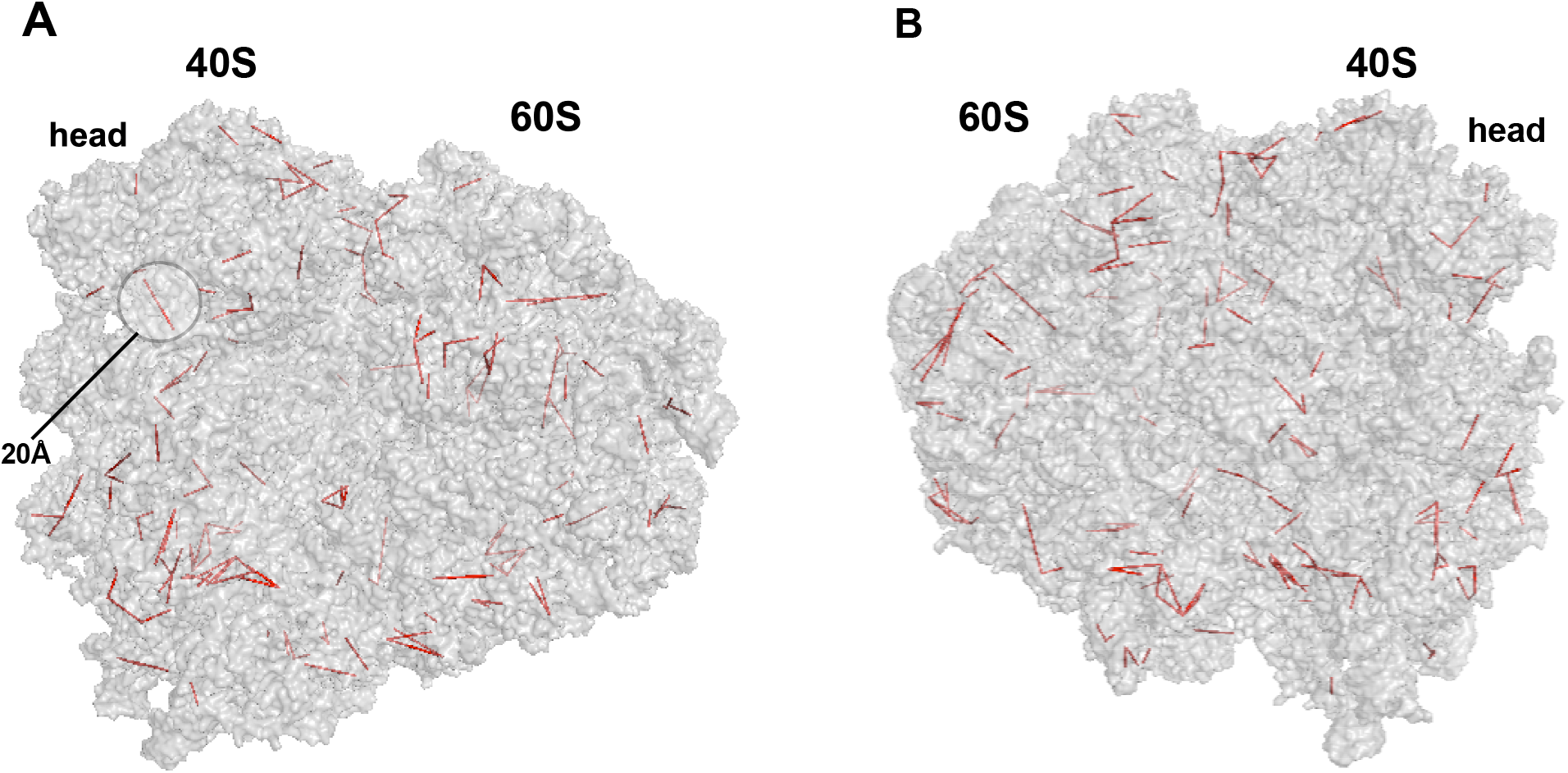
Inter- and intramolecular CL of r-proteins are modeled onto human 80S ribosome structure (5AJ0). Red line represents the CL distance between Cα of r-protein lysine residues. All the rProt - rProt crosslinks are within the distance spanned by the cross-linker BS(PEG)5 when mapped onto the ribosome structure, validating our cross-linking protocol. The circle highlighters CLs within a 20 angstrom distance.

**Figure 7.**
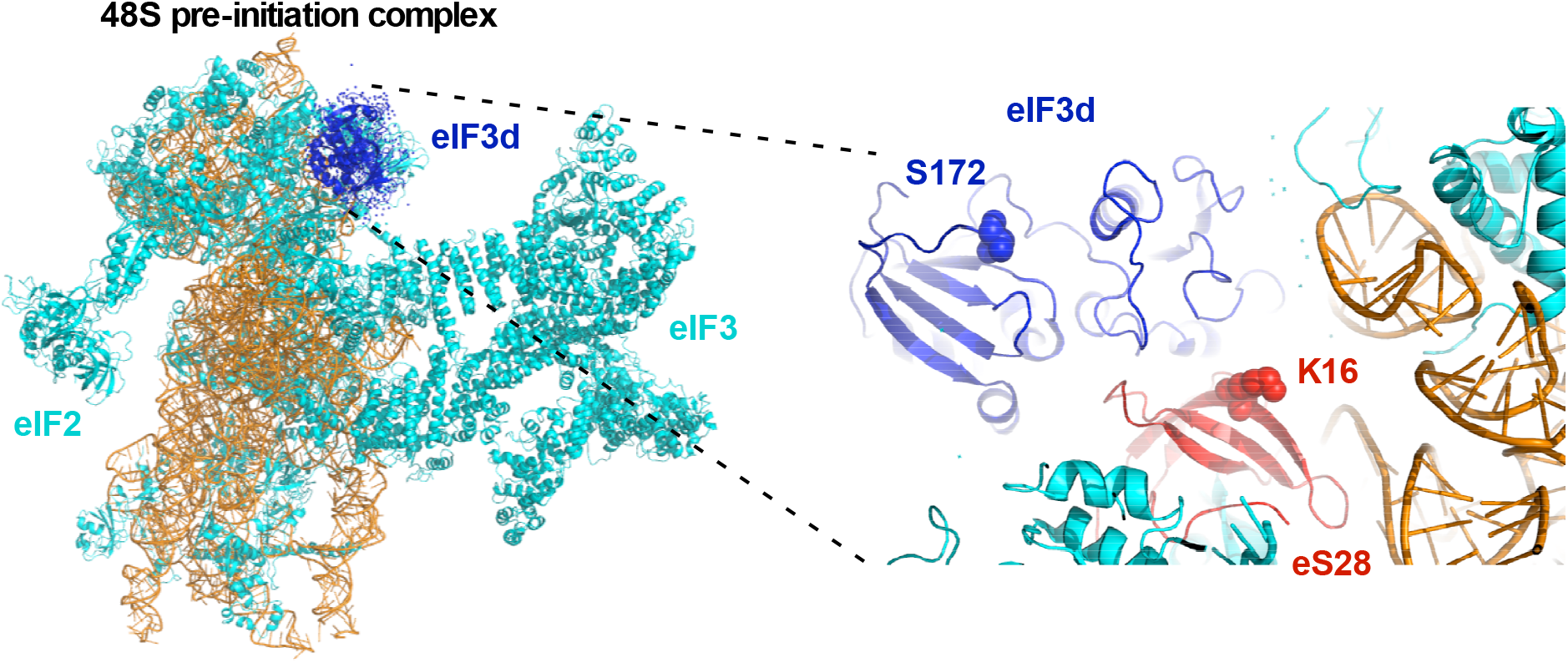
eIF3d location on the 48S pre-initiation complex. Cross-links were mapped to the ribosome-bound structure of eIF3d (PDB id: 6fe). (A) The overall view of 48S pre-initiation complex with eIF3d is highlighted as blue cartoon. (B) eS28 is interacting with eIF3d but the identified CL cannot be visualized as the deposited eIF3d is missing residues 1-171. The CL between eIF3d or G3BP2 and eS28 are as follows: eIF3d (142) – eS28 (16), eIF3d (248) – eS28 (16), G3BP2 (332) – eS28 (16), G3BP2 (248) – eS28 (16). One can imagine that eIF3d (172) chain will be extended toward eS28 (residue 16).

Other functionally relevant cross-links between r-proteins and RBPs with a known role in the regulation of ribosomal activity/translational control include Serbp1 (CL to uS4, eS10, eS12, and RACK1), Pa2g4 (CL to uL29), Ttc5 (CL to uL24), Ccdc124 (CL to eS30) and Btf3 (CL to eL31) in our RNA granule preparations (Table 1). Some of those cross-links could be directly mapped to the existing structures of Serbp1/Pa2g4/Ttc5/Ccdc124/Btf3 on the eukaryotic ribosome where the cross-linked residues are seen to lie within the distance spanned by the spacer arm of the cross-linker (Supplementary Figure S10). With Serbp1, the cross-links to eS10 and eS12 are consistent with the protein’s placement on the head of the 40S subunit as observed by Anger and coworkers (Anger *et al*, 2013). Likewise, the cross-link of Ttc5 to uL24 agrees with the data of Lin and coworkers showing uL24 as the unique protein binding partner of Ttc5 near the peptide exit site (Lin *et al*, 2020). Similarly, the cross-link between K238 of Pa2g4 and K43 of uL29 places Pa2g4 close to the peptide exit site on the 60S subunit and within an interaction distance of eL19, uL23 and uL29 as was observed by translationally inactive 80S ribosome (Bhaskar *et al*, 2021).

In addition to those RBPs, we identified cross-links between r-proteins and proteins involved in ribosome biogenesis (Pelp1, Gtpbp4, Nop58), ubiquitin-dependent protein degradation (Zer1) as well as RNA splicing (Nop58, Hnrnpc, Snu13, Sf3bf3) and stress granule assembly (Casc3).

### Neuronal RNA granules are stalled at a polysomic state

We therefore sought to resolve the structure of an individual ribosomal particle within an nRNAg. For this, a data-set of nRNAg complexes purified from rat cortex-hippocampus tissue and subsequently treated with RNase T1 (to improve the particle separation) was collected. Titan Krios I TEM with Falcon 3 detector in linear mode was used with the magnified pixel size at 1.41 Å. As there was little difference between the cryo-EM maps of intact vs. RNase T1 treated nRNAg, we will discuss the findings of the higher resolution map (RNase T1 treated).

In total, 3644 micrographs were collected with the RNase T1 treated nRNAg sample. Relion 3.1 software (Zivanov *et al*, 2018) was used to process the single-particle cryo-EM data. As seen from the 2D classification step, we were not able to identify any 2D classes of 48S pre-initiation complexes that are thought to be the predominant form of ribosomes in stress granules (Anderson & Kedersha, 2009; Bentmann *et al*., 2013; Buchan *et al*., 2011; Kedersha *et al*., 2016) (Supplementary Figure S11A). Also, proteomic analysis does not show a bias towards the small ribosomal subunit proteins (Supplementary Table S1), further indicating that the purified complexes are not stress granules.

After 2D classification (Supplementary Figure S11A) and multiple 3D classifications (Fig. 8A or Supplementary Figure S12), 3D class Cl2 (62 369 particles) was selected for further data-processing. The other classes (Cl3 and Cl4) did not show significant differences from Cl2 as seen from correlation value and visual inspection (Supplementary Figure S12). The Cl2 was further refined using unmasked and masked refinement, the resolution at FSC 0.143 was reported to be 2.95 Å (Supplementary Figure S11B). Per-particle, reference-based beam-induced motion correction step improved resolution to 2.87 Å and final map is shown in Figure 8B). Since no rat ribosome structure currently exists, we adapted the human ribosome structure (PDB ID 6ole) for refinement as it’s conformation is closest to our structure (Figure 9). Phenix real-space refinement (Liebschner *et al*, 2019) was used to refine human 80S ribosome structure (PDB ID 6ole) against the map. The human ribosomal protein sequences were adapted to the corresponding rat sequences in the refined structure by using Coot software (Emsley *et al*, 2010). Total RNA was extracted from the nRNAg and variable regions of 18S rRNA were sequenced to adopt the sequence differences between human and rat rRNA on structure (Supplementary Figure S13). A similar analysis was performed with the 28S rRNA regions (Supplementary Figure S14). The 5S rRNA and 5.8S rRNA sequences are identical between human and rat when compared using nucleotide blastn suite of NCBI.

**Figure 8.**
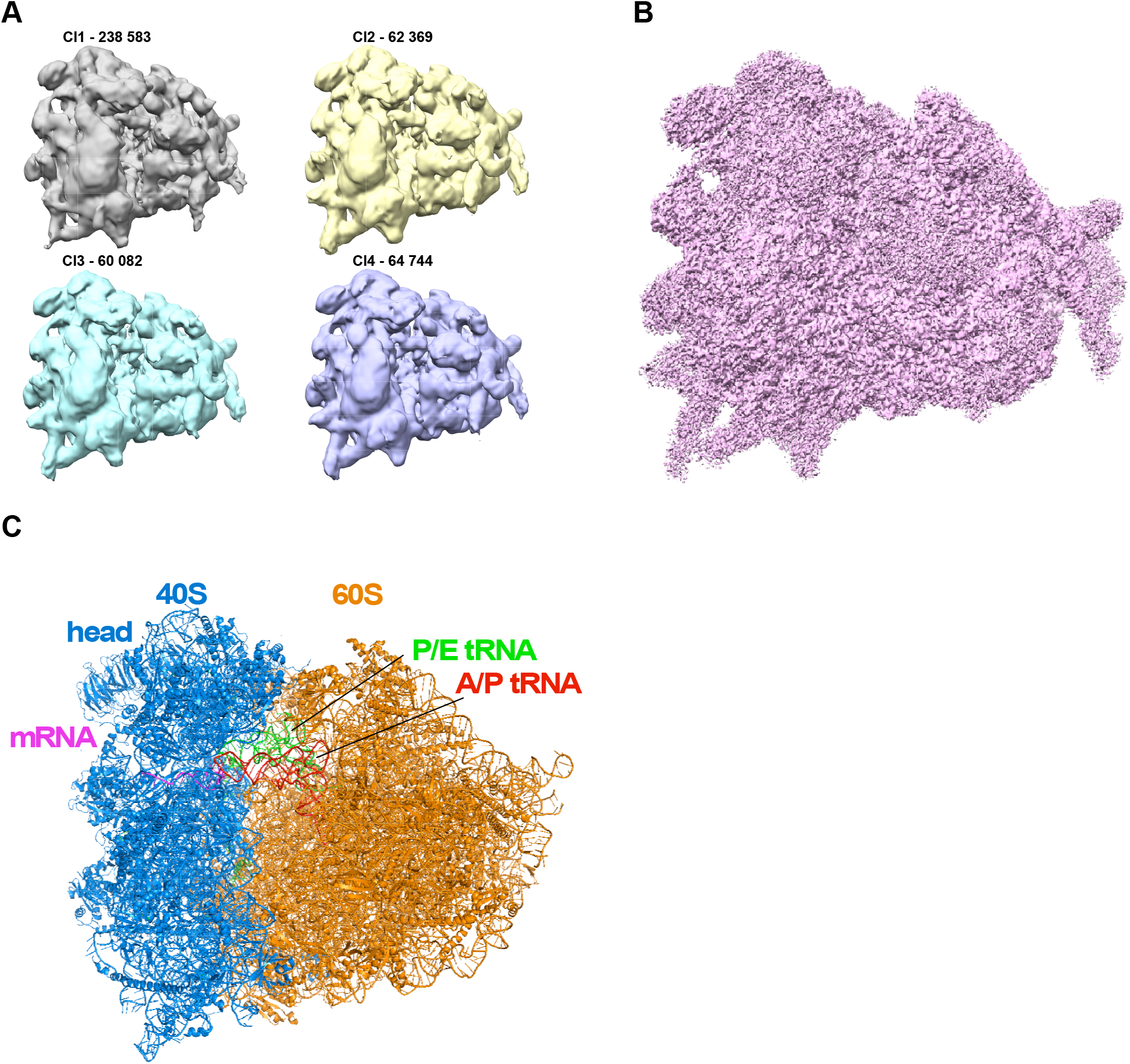
Cryo-EM data-processing workflow of RNase T1 treated nRNAg. (A) The 3D classification of 462 031 particles achieved from 2D classification step were classified into 4 classes. Particle distribution is indicated in brackets. Overall, the 3D classes have similar conformation but have minor differences. Each 3D class was further classified (Supplementary Sigure S12). (B) Cryo-EM map after polishing step at 2.87 Å resolution. (C) The Phenix software was used to refine cryo-EM map against human ribosome structure (PDB: 6ole) using real-space refinement. The 80S ribosome structure is presented in cartoon mode, 40S is in light-blue, 60S orange, mRNA red, A/P tRNA green, and P/E tRNA in magenta color.

**Figure 9.**
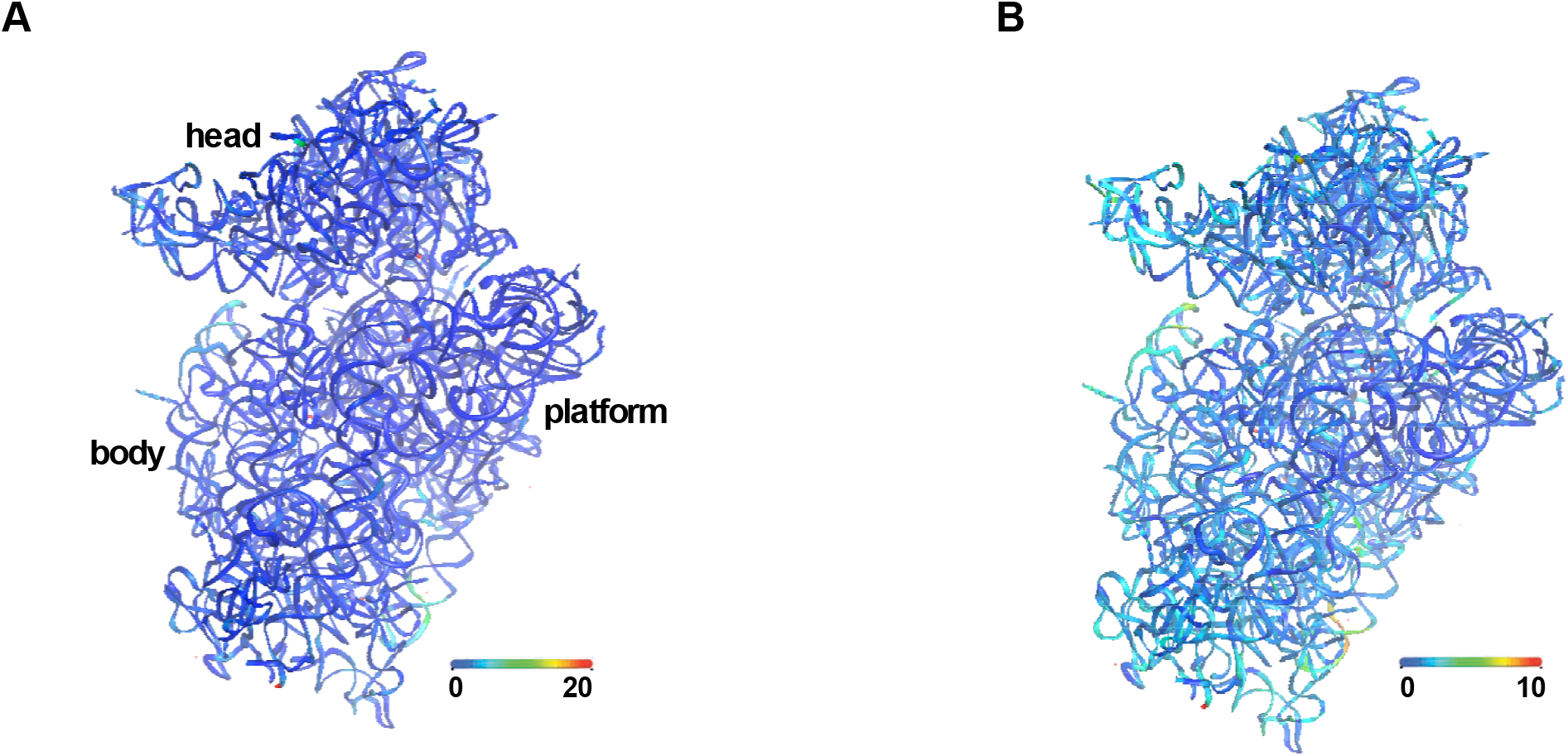
The difference heatmaps of ribosomes showing conformational similarities between two structures. Shifts between equivalent RNA phosphorus atoms and protein Cα atoms in the rat nRNAg ribosome structure (current study) and human 80S complexes stalled at elongation state (Li et al., 2019) are color coded as indicated by the scale. (left) panel represents 0 to 20 Å scale, (right) 0 to 10 Å scale. Ribosomes were superimposed by using the 60S subunit as the frame of reference. Overall, while the structures have similar conformation, slight differences are seen in peripheral parts of the ribosomes that are known to be flexible.

The refined structure resembles polysomic ribosomes with tRNAs in hybrid A/P and P/E states and contains nascent chain (4 residues long) in peptide exit tunnel (Figure 8C). The overall conformation of the ribosome is similar to Li et al., 2019 human 80S complexes stalled at the elongation state with the compound PF846 (Figure 9) (Li *et al*, 2019) or Budkevich et al., 2011 pre-translocation complex Rotated-2 state (Budkevich *et al*, 2011). This kind of rotated state structure precedes the mRNA and tRNA translocation step that moves them to the next codon, presumably impeding the action of eEF2 (Dever & Green, 2012; Li *et al*., 2019). These results indicate that the ribosomes in RNA granules are stalled at the polysomic translation elongation state as proposed by (Graber et al., 2013) and more recently by (Langille *et al*., 2019), where mRNAs are repressed at the elongation stage awaiting translational reactivation upon appropriate synaptic signals. A hybrid A/P state of tRNA may explain why stalled nRNAg complexes can bind puromycin as seen by Graber et al., 2013, as the 60S subunit A-site is not occupied by tRNA. Our results support the idea that the majority of the ribosomes in the complex are in a polysomic state. How the RNA binding proteins stall the ribosomes in nRNAg remains elusive.

As the majority of the ribosomes in our data-set are similar in composition and conformation and as it is known that when macromolecules exhibit continuous molecular motions, it is difficult to separate individual states by classical approaches, as one would need an infinite amount of classes to describe a continuum (Nakane & Scheres, 2021). We assumed that ribosomes in nRNAg should be more heterogenius than what was observed after multiple 3D classification steps (Supplementary Figure S12). The classical approach did not result in classes that significantly differ from the consensus map. The masked or focused refinements provide an efficient way to obtain high-resolution reconstructions. In our case, the masked classification with partial signal subtraction (40S and 60S) did not show any signs of deviation from consensus map.

The 80S ribosomes are intrinsically dynamic and adopt different conformational states during protein synthesis. The process of inter-subunit rotation is one of the principal motions where tRNA’s adopt hybrid configurations and it can occur spontaneously (Budkevich *et al*., 2014). Ribosomal subunits move in respect to each other in a ratchet-like inter-subunit motion (Budkevich *et al*., 2011; Frank & Agrawal, 2000). The ratchet-like motion is coupled with the movement of tRNAs to intersubunit A/P and P/E hybrid states (Agirrezabala *et al*, 2008; Blanchard *et al*, 2004; Budkevich *et al*., 2011; Munro *et al*, 2007). We used Relion multibody refinement approach to deal with continuous structural heterogeneity in cryo-EM (Nakane *et al*, 2018; Nakane & Scheres, 2021). The multibody refinement detected the ratcheting motion as the primary motion in our data-set (Movie S1). The ribosomal subunits rotate 12.3° (∼ 44 Å) with respect to each other. During the ratcheting step, the tRNA acceptor ends (3’ CCA) move from A to P and P to E sites (Movie S2). This amount of movement has also been reported by others and agrees with the notion that the ratcheting is the main principal motion that drives the translocation of the ribosome (Brilot *et al*, 2013; Flis *et al*, 2018).

Whereas the 40S subunit “rolling” motion is described as a 40S subunit rotation ∼6° toward the L1 stalk around the long axis of the small subunit (Budkevich *et al*., 2014). The subunit rolling changes the intersubunit space and causes a reciprocal opening and closing of the A- and E-site regions (Budkevich *et al*., 2014). It has been reported that the distance between the 40S and 60S subunits on the A-site of the 80S ribosome decreases by about 13–15 Å during subunit rolling from the POST state to the classical-1 PRE state (Budkevich et al., 2014). The multibody refinement on our data also detected the “rolling” motion (Movie S3). We measured the “subunit rolling” range in the A-site. When farthest points (28S rRNA U1961 – 18S rRNA A554) with respect to the rolling axis are selected to measure the distance, the widening of the A-site is ∼13 Å, but if more proximal points (28S rRNA U4381 – eS30 K18) are chosen near the A-site tRNA, the opening is 9 Å. The opposite was reported for the E-site, which is narrower in the POST state than in the classical-1 PRE state but due to the smaller distance from the rotation axis, the underlying movements were in the range of 6–7 Å at the E-site (Budkevich *et al*., 2014). Our measurements show 6 Å difference in the E-site when distance is measured between residues eL42 (P75) – uS7 (N186). Our “rolling” movement results are therefore similar to the previously reported results by Budkevich et al., 2014. However, the question remains as to what state the consensus map represents in the “rolling” movement landscape? The Relion multibody refienent software outputs 10 maps (called bins) that represent the specific principal motions (movies are made using these 10 maps). The bin1 and bin10 maps represent the largest differences when compared to each other. Our consensus map that is refined to high resolution lies in the middle of these two extremes.

### 40S head swivel

In addition to the ratcheting and rolling motion of the ribosome, the 40S head can move independently from the rest of the small subunit (body/platform). It was established in a bacterial system that the translocation of a tRNA on the 30S subunit parallels the swiveling of the 30S head and is coupled to the unratcheting of the 30S body (Ratje *et al*, 2010). The head swivel is coupled to the tRNA anticodon stem-loop movement during the translocation step that is mediated by EF-G/eEF2. It was shown in the mammalian system that back-rotation and back-rolling, together with the swivel-like motion of the 40S head, provide a significant shift of the tRNA_2_ -mRNA module relative to the 60S subunit and the 40S body/platform in the direction of translocation, resulting in compacted TI-POST-1 chimeric ap/P and pe/E hybrid states (Flis *et al*., 2018).

Our multibody refinement detected the head swivel and unratcheting of the 40S subunit as well. For the multibody refinement, 40S subunit was divided into two segments (40S head-tRNA and 40S body/platform) and 60S as third body. Interestingly, the multibody refinement detected two principal movement events associated with head swiveling and 40S unratcheting (Movie S4 and S5). In the first principal motion of the 40S head and body, the head moved only slightly ∼ 4.7 - 7.7 Å, whereas body moved as much as 19.4 Å (Movie S4). In the second motion, 40S head moved as much as ∼ 28 Å and body ∼5.4 Å (Movie S5).

### The “rocking” motion

We report a new kind of motion of the ribosome that is opposite to “rolling” motion and call it “rocking” (Movie S6). The “rocking” motion is detected by Relion multibody refinement when the ribosome is divided into 2 bodies (40S-tRNA and 60S). During the “rocking” motion the 40S subunit rotates ∼ 5° (11 Å) toward the 60S subunit around the horizontal axis of tRNA 3’CCA-end (Figure 10). The entire 40S subunit moves in a manner where the 40S subunit head moves toward but the 40S bottom part (spur-body) moves away from the 60S subunit or vice versa. The tRNA T-arm (elbow) becomes twisted from the 60S subunit by ∼ 2 Å. The tRNA’s (A/P and P/E states) anticodon stem-loops move as much as 6.2 Å but the acceptor-ends remain immobile. The intersubunit bridges B1a and B1b/c stretch during the “rocking” motion (Movie S7). Comparing the motion maps with our consensus map reveals that the consensus map is closest to the bin5 map. The bin1 map is more distinct from the consensus map than bin10 map. The question remains as to what kind of functionality is represented by the “rocking” motion? Is it related to the stalling conformation that is required during transport of RNA granules?

**Figure 10.**
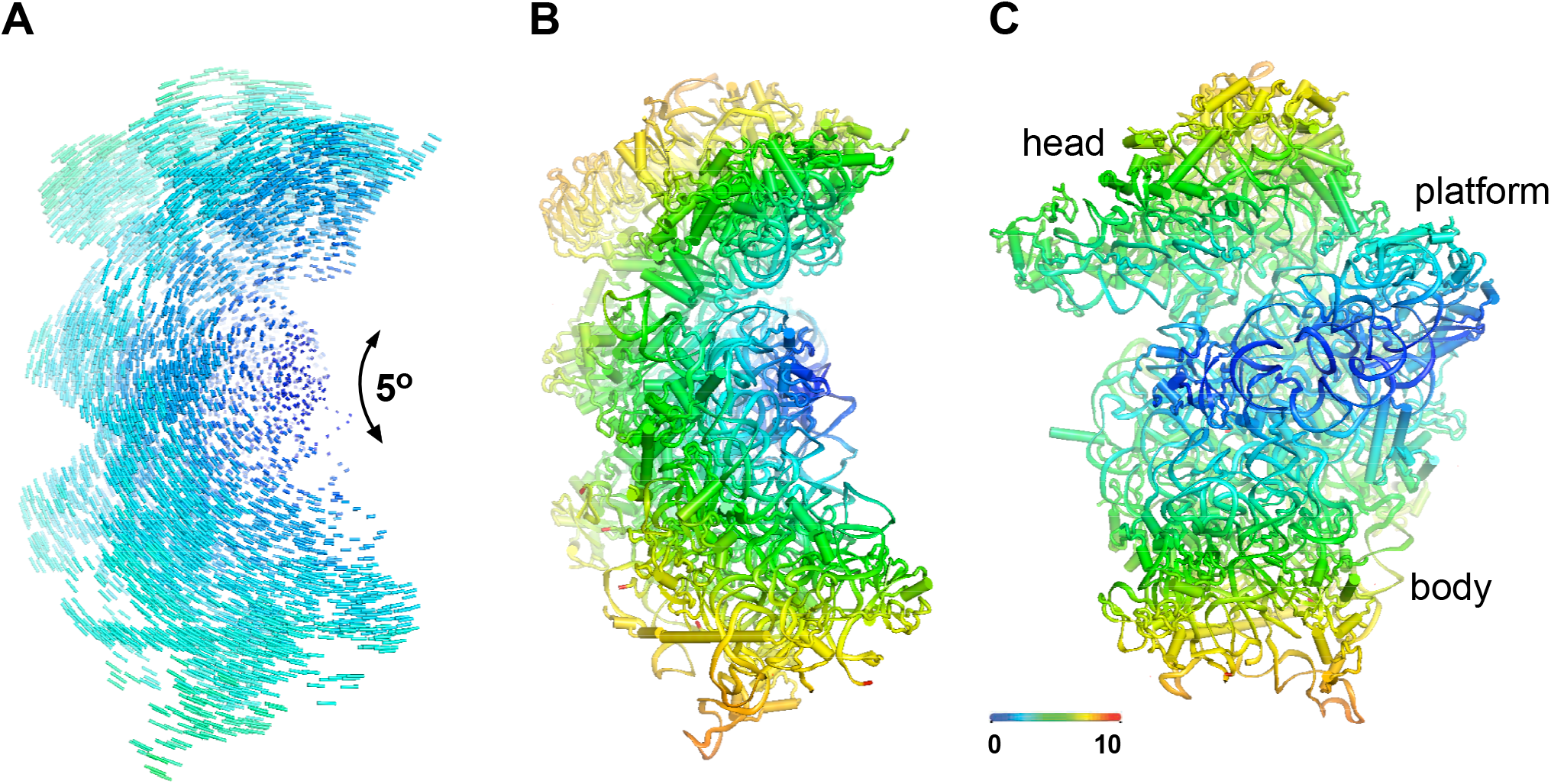
The difference heatmaps of ribosomal “rocking” motion representing the moving parts of the ribosome. The heatmaps were generated using cryo-EM maps outputted by Relion multibody refinement. The two low-resolution maps of “rocking” motion that represent the largest differences (bin1 and bin10) were chosen for Phenix real-space refinement against the high-resolution consensus structure determined in this study. The refined coordinates were aligned based on the 28S rRNA using Pymol and heatmaps were generated using Pymol modevector scripts. (A) Shows the vector distribution of the 40S ribosome from the A-site view. The rotation center is in the middle of the intersubunit face. (B) Same view as in the panel (A) but 40S subunit is shown as cartoon and color-coded based on the movement length in angstroms, represented by the scale bar. (C) The 40S subunit is viewed from the intersubunit side. The blue line running through the small subunit displays the rotational axis of the “rocking” motion.

One possibility is that the “rocking” motion represents the 40S head and body motions that cannot be separated by the rigid 40S subunit and therefore these motions are trapped inside the “rocking” motion. Therefore, we conducted the multibody refinement with 3 bodies (40S-head-tRNA/40S-body/60S). As previously, multibody refinement detected ratchet like motion and head swiveling. The fifth principal motion that was detected displayed a “rocking”-like motion where the 40S body behaved similarly to the “rocking” motion but the 40S head only moved slightly compared to the coupled 40S movement (Movie S8 and S9). The 40S body moved as much as ∼ 14.4 Å and head ∼ 4.3 Å. During the 40S body movement the eS17 helix (residues 70-82) becomes stretched out (Movie S9). The mRNA entrance channel formed by uS3 and 18S rRNA helix 16 (h16) closes and opens during “rocking” motion.

## Discussion

We choose the cortex-hippocampus brain tissue to study neuronal RNA granules as these brain regions are preeminently involved in different memory related tasks. Synaptic plasticity and long-term memory have been linked to the nRNAg components (e.g. Caprin-1) (Nakayama *et al*., 2017). The hippocampus is involved in recognition and spatial memory and transfers the information into cortical regions to give memories meaning and connection to other memories (Huerta *et al*, 2000; Tsien *et al*, 1996). It is also involved in consolidation where new learned information is transformed into long-term memories (from short-term to long-term memory). It was shown recently that the cerebral cortex is really responsible for long-term memories and not hippocampus (Hasan *et al*, 2013). Therefore the hippocampus provides the necessary environmental cues, which are transmitted to the cortex where learning-dependent associations take place. The cerebellum is involved in procedural memory, and motor learning whereas emotional (e.g. fear memory) memory is associated with amygdala.

The molecular mechanisms of memory storage are still elusive. It is thought that memories are distributed in neural networks (Bassett *et al*, 2013; Ferbinteanu, 2019). Long-lasting memories likely result from structural changes such as the growth of new synapses and associated neural networks (Kolb & Gibb, 2014). Memories that last a lifetime are thought to be stored, at least in part, as persistent enhancement of the strength of particular synapses (Jalil *et al*, 2015). Amnesia and dementia, as well as addiction and post-traumatic stress disorder, are associated with deficient or pathophysiological memory induction and maintenance (Jalil *et al*., 2015). The fundamental issue here is how memories can be maintained over extended periods of time despite protein turnover and diffusion (Crick, 1984). “Molecular switch” has been proposed as a solution (Tyson *et al*, 2003; Verdugo *et al*, 2013). A local molecular switch could be placed at the level of post-translational modifications (Lisman, 1985; Routtenberg & Rekart, 2005) or at the level of protein synthesis (translation) (Abraham & Williams, 2008; Belelovsky *et al*, 2005), as translation can occur and be regulated locally in dendritic compartments (Batish *et al*., 2012; Leal *et al*, 2014; Santos *et al*, 2010; Sutton & Schuman, 2006) in the form of neuronal RNA granules.

We show that nRNAg’s purified from the cortices of newborn Wistar rats are morula-like particles that contain ribosomes and RBPs. When analyzed by negative staining electron microscopy, the granules appear morphologically similar to whole brain-derived neuronal RNA granules reported in previous studies (Krichevsky and Kosik 2001; Graber *et al*. 2011; El Fatimy *et al*. 2016) and mostly dissociate into monosomes or smaller polysomes upon RNase treatment (Figure 2). The RNase sensitivity of our cortex-hippocampus-derived RNA granules seems to stand in contrast to the results of Graber *et al*. and El Fatimy *et al*. where the granules were largely resistant to RNase A/MNase treatment under condition similar to the present study. However, a recent study by Anadolu and coworkers indicates that the extent of RNase sensitivity of the granules is affected by *i.a.* monovalent salt concentration. According to Anadolu *et al*., increasing the salt concentration above the physiological level leads to a decompaction of the RNA granules with a concomitant increase in nuclease sensitivity. Though our nuclease sensitivity analysis was only performed at the physiological salt concentration, we note that the particle composition of our nuclease treated samples in ns-EM is broadly similar to that observed by Anadolu *et al*. upon RNase treatment in the absence of a high-salt preincubation of the granules. We therefore surmise that the precise extent of nuclease sensitivity of RNA granules may depend on factors like the degree of compaction of the RNA granules in a particular preparation and/or the type of nuclease used. Indeed, we observed a somewhat higher fraction of nuclease-resistant particles in an MNase-treated sample compared to RNase T1 treatment (Supplementary Figure S2).

We also purified nRNAg from retinoic acid differentiated human neuroblastoma cell line SH-SY5Y. The major hurdle with neuroblastoma is the glycogen contamination as the particles are similar in size. The glycogen contamination does not interfere the identification of nRNAg components, but interferes cryo-EM analysis. Glycogen granules can be reduced by using pull-down strategies targeting specific component of nRNAg’s but this leads to bias toward the specific component.

Caprin-1 together with G3BPs are important RBPs that are present in our nRNAg sample. It has been shown that they work in collaboration in stress granules (Kedersha *et al*., 2016; Solomon *et al*, 2007). Caprin-1 promotes the assembly of RNA granules and is responsible for the transport of its bound mRNAs in cultured cells (Kedersha *et al*., 2016; Shiina *et al*., 2005; Shiina & Tokunaga, 2010). Knockdown and knockout (KO) of Caprin-1 in cultured neurons causes a reduction in the synaptic connections on dendrites and the density of neural networks (Shiina & Tokunaga, 2010; Shiina *et al*, 2010). A heterozygous nonsense mutation in the caprin1 gene has been found in a human patient with autism spectrum disorder (ASD), and heterozygous KO of Rng105 gene in mice causes ASD-like behavior (Jiang *et al*, 2013; Ohashi *et al*., 2016). Caprin-1 has been shown as an essential element of RNA granules for establishing long-term memory, and mediates dendritic localization of mRNAs as an underlying mechanism for AMPA receptor (AMPAR) scaling, synaptic strength and plasticity, and long-term memory formation (Nakayama *et al*., 2017). Caprin-1 abundance is relatively high compared to other RBPs in our proteomic data based on the calculations of relative iBAQ intensities that are compared between proteins present in nRNAg (Supplementary Table S4). According to this analysis there is 1 molecule of Caprin-1 per 9 molecules of ribosomes in nRNAg. It was proposed that Caprin-1 and G3BP-1 were directly or indirectly associated in a stable complex in HEK293T cell extract pull-down experiments and overexpression induce the stress granule formation (Solomon *et al*., 2007). The Caprin-1 region 352 to 380 is responsible for the binding to G3BP-1 NTF-2-like domain (residues 1 – 141) (Solomon *et al*., 2007). We were not able to identify direct cross-links between Caprin-1 and G3BP1 in nRNAg sample, although both of them are present in nRNAg sample (G3BP1 having relative abundance fold value 24 and G3BP2 having 8). As we only identify cross-links between protein amine groups, then one possibile reason, why we do not see the interaction between Caprin-1 and G3BP’s, is that there are no available amine groups within the optimal distance on the surface of the proteins to form the covalent bonds with a cross-linker. At the same time, it was shown by others that Caprin-1 and G3BP1 can bind to each other, but the ability to induce SG formation or enter them did not depend on their association with each other (Solomon *et al*., 2007). Instead, we were able to identify link between Caprin-1 (residue 136) and Pelp1 (residue 1) by using leiker cross-linker (Tan *et al*, 2016). Proline-, glutamic acid- and leucine-rich protein 1 (PELP1) is RBP involved in transcription regulation and 60S ribosomal subunit synthesis and ribosomal RNA transcription (Castle *et al*, 2012; Finkbeiner *et al*, 2011). The two proteins have been shown to be associated with MID1 complex that regulates translation through mTOR and involved in the phosphorylation of tau which hyperphosphorylation induces amyloid- (A ) peptides, and neurofibrillary tangles formation in Alzheimer’s disease (Matthes *et al*, 2018). Our cross-linking results show a so far uncharacterized connection between these two proteins in the neuronal RNA granules.

It has been shown that cells lacking both G3BP1 and G3BP2 cannot form SGs in response to eIF2α phosphorylation or eIF4A inhibition (Kedersha *et al*., 2016). The G3BP1-F33W, a mutant unable to bind G3BP partner proteins Caprin-1 or USP10, can rescue the SG formation (Kedersha *et al*., 2016). Caprin-1 binding to G3BP’s promotes, but USP10 binding inhibits, SG formation (Kedersha *et al*., 2016). USP10 is also present in our nRNAg sample, although much less abundant than G3BPs and Caprin-1 having 124 fold difference to ribosomes (Supplementary Table S4). It remains unclear if the interaction of G3BP’s and Caprin-1 is required for nRNAg formation as we did not see any direct cross-links between these two proteins. It may also indicate that after formation of nRNAg the contacts are no longer maintained. Immunoprecipitation studies show that G3BP1 can interact with 40S ribosomal subunits through its RGG motif, which is also required for G3BP-mediated SG formation (Kedersha *et al*., 2016). More specifically, small ribosomal subunit proteins eS6 and uS12 co-immunoprecipitated with G3BP1 (Kedersha *et al*., 2016). Our cross-linking results show that G3BP2 may interact with small ribosomal protein eS28. eS28 is eukaryotic specific r-protein that interacts with mRNA in the exit tunnel. eS28 residue 16 gives cross-links with G3BP2 residues 248 and 332. This cross-link is specific to G3BP2 as we do not detect G3BP1 cross-links with ribosomal proteins. The identity between human G3BP1 and G3BP2a is 64% and G3BP1 and G3BP2b is 59%. The G3BP2 “binding-site” is special as eIF3d subunit can independently bind to this site and regulate translation of sub-set of mRNA’s (Lee *et al*, 2016). This may indicate that G3BP2 could also mediate translation of specific mRNAs. Beside inhibition, it has been shown that G3BPs can promote protein synthesis of interferon-stimulated genes that are involved in antiviral activity, and in cancer progression and metastasis (Alam & Kennedy, 2019). The possible G3BP2 binding site on the ribosome needs to be further studied.

The microtubule and filament association with nRNAg’s indicate their involvement in transporation that is known phenomenon (Balasanyan & Arnold, 2014; Heisler *et al*., 2011; Hirokawa, 2006). Lesser-known is the lysosome/vesicle involvement in the transporatation of RNA granules. Recent studies have shown that axonal lysosomes co-traffick with RNA granules labeled with G3BP1 (Sahoo *et al*., 2018), TDP-43 (Alami *et al*., 2014; Gopal *et al*., 2017), and Caprin-1 (Nakayama *et al*., 2017). We show that lysosomes/vesicles are in close proximity to nRNAg’s using ns-EM. The proteomics analysis identified Rab proteins that are markers for different membrane bound organelles (Stenmark, 2009; Tang *et al*., 2014). The seven Rab proteins identified in this study in nRNAg sample give a clue as to what kind of membrane containing organells are present. Further experiments are required to investigate the involment of these Rab proteins in nRNAg transport or storage in more detail. It is known that some Rab proteins are involved in neuronal processes. The Rab18, Rab11a and Rab11b (were also identified in our study) are part of recycling endosomes (RE) and neuronal RE’s regulate retrograde neurotrophic signaling, axonal pathway fixation during protein development, renewal and degradation, vesicle recycling, and synaptic plasticity, among other processes (Dittman & Ryan, 2009; Itofusa & Kamiguchi, 2011; Kennedy & Ehlers, 2006; Rozes-Salvador *et al*, 2020; Winckler & Mellman, 2010). Rab7A is involved in the trafficking of endosomes, autophagosomes, and lysosomes, and regulates tau secretion (Rodriguez *et al*, 2017). In Alzheimer’s disease (AD), tau protein accumulation in somato-dendritic compartments is the main cause of neurodegeneration and RAB7A gene expression and protein levels are up-regulated in AD patients (Ginsberg *et al*, 2011; Ginsberg *et al*, 2010). Also, Rab18 is involved in liquid droplet formation and facilitates dengue virus infection (Tang *et al*., 2014).

Our cryo-EM data indicates that majority of ribosomes are stalled in the state that resembles polysomic pre-translocated state where tRNA’s are in A/P or P/E state. The tRNA hybrid states are favored within the mammalian complex (Budkevich *et al*., 2011). Hybrid tRNA positions are defined by a spontaneous movement of A- and P-site tRNAs in the direction of translocation on the large ribosomal subunit, while the anticodon portions of tRNA remain fixed with respect to the small subunit (A/P and P/E hybrid sites, respectively) (Budkevich *et al*., 2011; Flis *et al*., 2018). Hybrid tRNA configurations have been suggested to serve as *bona fide* intermediates in the translocation process by lowering the activation barrier for translocation (Dorner *et al*, 2006; Semenkov *et al*, 2000). The rolling motion of the ribosome converts the POST to the classical PRE state and occurs during the accommodation step of A-site tRNA (Budkevich *et al*., 2011). We discovered a new kind of ribosome conformational change that we call “rocking” motion. The “rocking” motion has been proposed previously by Brimacombe and coworkers using UV cross-linking assay (Stiege *et al*, 1988). According to their intrepetation of the “rocking” motion was that tRNA binding deep in the cleft of small ribosome in the presence of mRNA causes a “rocking” motion of the head region of the subunit, leading to allosteric protections in the helix 18 region. The rocking motion would likely be accompanied by a corresponding slight opening and closing of the cleft, which would effectively create a molecular turnstile”, allowing the tRNA molecules to pass through (Stiege *et al*., 1988). It is hard to interpretate what kind of motion they are positing as rolling motion opens and closes the A- and E-site regions (Budkevich *et al*., 2014). The “rocking” motion does not seem to have an impact on the peptidyl transferase center of the ribosome as the tRNA acceptor ends do not exhibit movement. One possibility is that the “rocking” motion plays a role in the E-site tRNA exiting as this kind of motion can loosen the contacts between 40S head and 60S subunit or the L1-stalk. On the other hand, the anticodon stem-loop of P/E site tRNA changes 6 Å, but the movement of tRNA is not toward the exiting site, instead the anticodon part moves perpendicular to exiting. We argue that the “rocking” motion can be part of the stalling mechanism where ribosomes adopt a conformation that does not support the translation due to their inability to bind translation factors. Also, it may reflect the translation factor binding to the A-site that pushes the 40S body in the outside direction. For example, binding of eEF2 to the ribosome during the elongation step induces the power stroke that unlocks the ribosome to initiate translocation (Chen *et al*, 2016; Noller *et al*, 2017). Although, classification and refinement maps to not show any density for A-site bound translation factors.

The main conclusion of our study is that nRNAg’s contain polysomic ribosomes that can escape the rate limiting translation initiation step of protein synthesis. We hypothesize that G3BP2 targets the ribosomal E-site to stall or activate translation, dependent of mRNA spices.

## Materials and Methods

### Brain tissue isolation and lysis

Cortices along with hippocampi were harvested from 10 newborn (PN 2 - 6) Wistar rats, as described (Brewer & Torricelli, 2007; Nunez, 2008; Wareski *et al*, 2009). Rats were received from University of Tartu Laboratory Animal Centre. During the tissue extraction, brains were kept submerged in ice-cold Hibernate E medium supplemented with 1 × Glutamax and the extracted cortex-hippocampus tissue was stored in a separate container in the same medium until all cortex-hippocampi had been collected (for approximately 1 h). All tissue extraction was performed under a stereomicroscope (VisiScope SZBL350). Brains were stripped of the meninges prior to the extraction of the cortex-hippocampus. The collected tissue was twice washed with 4 mL ice-cold Ca^2+^/Mg^2+^-free 1 × HBSS solution (Gibco) and submerged in 6 mL ice-cold lysis buffer (20 mM Hepes-KOH, pH 7.5, 10 mM K(OAc), 4 mM Mg(OAc)_2_, 0.5 mM Tris(2-carboxyethyl) phosphine (TCEP) supplemented with a protease inhibitor cocktail (Roche,cOmplete Ultra EDTA-free; 1 tablet in 10 mL lysis buffer). The tissue was triturated in the lysis buffer by repeated pipetting with a wide bore 1 ml pipet tip until a homogenous suspension was obtained. The cells were lysed by passing the triturated suspension through a pre-cooled gauge 26 needle in 4 strokes in the presence of 1 mM phenylmethanesulphonyl fluoride (PMSF), 1 μg/mL bestatin (Merck, B8385), 100 U/ml RNase inhibitor (Thermo Scientific RiboLock), 1.25 U/mL DNase I (Promega, M6101) and 0.5 % (v/v) Tween-20 (Merck, P9416). Alternatively, cells in the triturate were lysed by a Dounce homogenization (Kimble, Dounce tissue grinder, 7 ml) through 10 strokes with a loose pestle followed by 5 strokes with a tight pestle under otherwise identical conditions. No significant differences in the yield or morphology of the RNA granules were observed with the two lysis methods (Supplementary Figure)

In a separate set of experiments, cells were lysed (syringe or Dounce lysis) in the presence of 100 μg/mL cycloheximide or 50 μM homoharringtonine. Cycloheximide is a translation inhibitor known to dissociate stress granules while homoharringtonine blocks the transition from initiation to elongation and thus causes a depletion of actively transcribing polysomes. However, an ns-EM analysis did not reveal any noticeable differences in the morphology of RNA granules obtained in the absence or presence of either CHX or HHT during lysis (Supplementary Figure S15).

### SH-SY5Y cell line differentiation

SH-SY5Y (ATCC; REF: CRL-2266) cells were cultivated in DMEM/Ham’s F-12 medium (Corning) supplemented with Fetal Bovine Serum (Corning) and penicillin/streptomycin (DMEM/Ham’s F-12/FBS/pen-strep) at 37° C/5 % CO_2_ under constant humidity. Upon reaching a 40 % confluency, the cells were replated/split to 8 × 15 cm cell culture dishes at an initial seeding density of 8000 cells/cm^2^ and cultivated in DMEM/Ham’s F-12/FBS/pen-strep as described above. Upon reaching a 20 % - 30 % confluency, the culture medium of the cells was replaced with Neurobasal medium (Gibco) supplemented with glutamax (Gibco), B27 supplement (Gibco) and 15 μM all-trans-retinoic acid (ATRA) and the cultivation continued for 5 days. The cells were harvested for RNA granule isolation as described in “RNA Granule isolation”.

### RNA granule isolation from brain tissue

Immediately after lysis, K(OAc) in the lysate was adjusted to 150 mM and the nuclei and mitochondria were removed from the lysate by centrifugation at 2500 × g/4° C for 2 min. The resulting post-mitochondrial supernatant was further clarified by centrifugation at 11 000 × g/4° C for 9 min and the resulting supernatant (5 – 6 mL) loaded on a 2 mL 60 % sucrose cushion in RNA Granule Buffer (20 mM Hepes-KOH, pH 7.5, 150 mM K(OAc), 4 mM Mg(OAc)_2_, 0.5 mM TCEP) in a Beckman SW41 tube. The tube was filled to top with 2 – 3 mL ice-cold RNA Granule Buffer and centrifuged for 2 h in a Beckman SW41 Ti swing-out rotor at 38 000 rpm (= 246942 × g; ω^2^t =1.139 × 10^11^ rad^2^/s). The supernatant liquid on top of the 60 % sucrose cushion was carefully removed with a pipet and the tube walls and the top of the cushion were washed with 1 – 2 mL ice-cold RNA Granule Buffer. After removing the sucrose cushion, the pellet (Pellet 1) was redissolved in ca. 100 μL RNA Granule Buffer and centrifuged through a 700 μL bed-volume Sephacryl-S500 (Cytiva) spin column (equilibrated with RNA Granule Buffer) in a fixed-angle rotor at 1000 × g/4° C for 1 min to obtain the RNA Granule fraction. The RNA granule preparation was aliquoted in 1/10^th^ original volume in “ProtLoBind” microcentrifuge tubes (Eppendorf), flash-frozen in liquid nitrogen and stored at – 80 C.

### RNA granule isolation from neuroblastoma cell line

For RNA granule isolation from the ATRA-differentiated SH-SY5Y cells, cells grown on 8 × 15 cm dishes to a 90 % confluency were washed with 2 × 5 mL ice-cold 1 × HBSS solution and harvested in a total of 4 mL ice-cold lysis buffer by mechanical scraping. The cells were lysed by passing the suspension through a gauge 26 needle in 4 strokes in the presence of 0.1 mM PMSF, bestatin (1 μg/mL), RNase inhibitor (Thermo Scientific RiboLock; 100 U/mL) and 0.5 % (v/v) Tween- The subsequent procedures were identical to that described for the cortex-hippocampus RNA granules.

### Glycogen removal by affinity chromatography

For the removal of glycogen particles from neuroblastoma sample, 20 μL of a Ni-Sepharose resin (Qiagen) was equilibrated with RNA Granule Buffer and incubated with 600 μg His-tagged maltose binding protein at room temperature for 15 min under constant agitation. The resin was pelleted by centrifugation at 100 × g/4° C for 15 s, the supernatant was removed and the resin washed three times with 500 μL RNA Granule Buffer through centrifugation (at 100 × g/4° C/15 s) and resuspension. The lack of a noticeable A_280_ absorbance in the supernatant indicated a near complete binding of the MBP to the resin. The 20 μL aliquots of the MBP-coated Ni-Sepharose resin were incubated with 5, 15, and 30 μL aliquots of an SH-SY5Y-derived RNA Granule preparation (containing 0.13, 0.38, and 0.75 A_260_ units of RNA Granules, respectively) at room temperature for 15 min under constant agitation to remove the contaminating glycogen. The samples were centrifuged at 100 × g/4° C for 15 s and the RNA granule containing supernatant withdrawn. 71 % to 82 % of the original RNA Granule material was recovered in the supernatants. 4 μL aliquots of the glycogen-depleted supernatant were deposited on glow-discharged Quantifoil grids for negative staining and analyzed as described above.

To analyze the nonspecific binding of RNA granule components to the Ni-Separose resin, 0.075 A_260_ units of cortical RNA Granules were incubated with 20 μL MBP-coated Ni-Sepharose resin (810 μg MBP/20 μL resin) as described above. After removing the RNA L ice-cold RNA Granule granule containing supernatant, the resin was washed with 3 × 500 μL Buffer. The resin was subsequently incubated with 50 μL 8 M urea/5 mM TCEP at room temperature for 10 min, followed by centrifugation at 100 × g/4° C/30 s. Eluted protein in the supernantant were anylysed by mass-spectrometry and Western blot.

### Negative staining electron microscopy

RNA granule preparations were diluted with RNA Granule buffer to an optical density A_260_ = 2 U/mL aliquots of the dilution deposited on glow-discharged Quantifoil grids (Quantifoil 1.2/1.3, 300 mesh copper) pre-coated with a continuous layer of 3 nm carbon. Grids were glow-discharged at 10 mA for 15 s (PELCO easiGlow Glow Discharge Unit). The grids were incubated in a custom-made humidity chamber at room temperature for 3 min and rinsed in 3 × 50 μL drops of RNA Granule Buffer for a total of 30 s. The excess buffer was blotted away from the grid by gently touching the side of the grid with a Whatman filter paper and the grids were negatively stained by incubation with 2 × 4 μL 2 % U(OAc)_2_ in two consecutive 1 min incubation steps at room temperature. The grids were dried on air and stored in glass dishes at room temperature in a dessicator. The EM images were aquired on a Tecnai 12 Bio-Twin transmission electron microscope (ThermoFicherScientific) operated at 120 kV using a room temperature side entry holder. Images were collected in a Gatan Orius camera system at a nominal magnification of 11 000 × at a defocus ranging from 2.5 – 3.5 μm.

### RNase treatment of RNA granules

All nuclease units are given in Kunitz units. For the RNase A sensitivity assay nRNAg were incubated with RNase A at a 0.084 activity units per 1 A_260_ unit of nRNAg in RNA Granule Buffer at room temperature for 30 min. The sample was immediately deposited to a glow-discharged Quantifoil grid coated with a 3 nm continuous carbon layer and analyzed as described above. For the RNase T1 assay, RNA granules were incubated with 37 U of RNase T1 per 1 A_260_ unit nRNAg in RNA Granule Buffer at room temperature for 30 min and thereafter immediately deposited to a glow-discharged Quantifoil grid coated with a 3 nm continuous carbon layer. For the MNase sensitivity assay, RNA granules were incubated with 25 or 200 U of MNase per 1 A_260_ unit of nRNAg in RNA Granule Buffer in the presence of 0.5 mM CaCl_2_ at room temperature for 30 min. The MNase digestion was stopped by the addition of EGTA to an f.c. of 1.5 mM and the sample was immediately deposited to a glow-discharged Quantifoil grid coated with a 3 nm continuous carbon layer.

### Immunoblotting

0.13 – 0.15 A_260_ units of RNA granules in 1 × SDS Gel Loading buffer were incubated at 95 C for 5 min and loaded to a 10 % SDS-PAGE gel. The proteins were resolved by electrophoresis at 180 V/50 mA at room temperature. The resolved proteins were transferred onto a 0.45 μm PVDF membrane (Immobilon, Merck) in ice-cold transfer buffer (25 mM Tris, 192 mM glycine; 600 mM MeOH ) at 80V/4 C for 85 min under continuous stirring. The membranes were blocked with 5 % milk (nonfat dried Milk, AppliChem) in Tris Buffered Saline with Tween (TBST) (50 mM Tris-HCl, pH 7.5, 150 mM NaCl, 0.05 % (v/v) Tween-20) at room temperature for 1 h with constant shaking. The membranes were incubated in 0.5 % milk in TBST for 1 h with the following antibodies : rabbit anti-rpl7a (Bethyl # A300-749A-T; at a 4 : 10 000 dilution), anti-caprin-1 (Invitrogen, PA5-96857; at a 7 : 10 000 dilution), anti-G3BP-1 (Millipore, 07-1801; at a 10 : 10 000 dilution), and anti-G3BP-2 (Invitrogen, PA5-53776; at a 6 : 10 000 dilution), followed by three 2-min washes with room-temperature TBST. The antibodies were detected by incubation with an HRP conjugated antibody (Goat Anti-Rabbit IgG (H +L) Peroxidase Conjugated, Pierce 31466; at a 1 : 10 000 dilution) in 0.5 % milk/TBST for 1 h at room temperature, followed by three 2-min washes with room-temperature TBST and incubation in 2 mL ECL (Cytiva, ECL Western Blotting Analysis System) developing solution. The detection and visualization of the protein bands was completed with Hyperfilm ECL (Cytiva).

### Chemical crosslinking of RNA Granules

For a lysine-specific cross-linking of RNA granule proteins with succinimidyl-based cross-linkers (BS3, BS(PEG)5, DSG, DSS) we adopted the protocol of Leitner and coworkers (Leitner *et al*., 2012) with some modifications. RNA granule preparation corresponding to 1 mg total protein (based on the A_280_ absorbance and 1 A_280_ = 1 mg/mL protein) was incubated with 1 mM cross-linker in RNA Granule Buffer at room temperature for 30 min in a “ProtLoBind” tube wrapped in an aluminium foil. The cross-linking reaction was quenched by the addition of NH_4_HCO_3_ to an f.c. of 50 mM and incubated at room temperature for 20 min. RNase A was added to the sample to an f.c. of 0.01 μg/μ evaporated in an Eppendorf Centricon Plus Concentrator at 30° C over 1 h. Proteins were denatured by incubation in Denaturation Buffer (8 M Urea, 100 mM Tris-HCl, pH 8.5, 20 mM CH_3_NH_2_, 0.5 mM TCEP) at 50 C for 30 min. Cysteines were blocked with 10 mM iodoacetamide at room temperature for 30 min and any remaining iodoacetamide scavenged with 10 mM dithiothreiol at room temperature for 10 min. Urea in the sample was diluted to 3 M with the addition of 100 mM Tris-HCl, pH 8.5 and RNA digested with 125 U Benzonase (Merck, E1014 ) at 37° C for 1 h. Following the nuclease treatment, urea in the sample was diluted to 1.5 M with 100 mM Tris-HCl, pH 8.5 and proteomics grade trypsin (Promega, V5280) was added to the sample at an enzyme-to-substrate ratio of 1:70 and the sample was incubated in the presence of 1 mM CaCl_2_ at 37° C overnight. After overnight digestion, the sample was acidified to 0.5 % trifluoroacetic acid and the tryptic peptides were purified by solid-phase extraction using Pierce C18 pipet tips (100 μL) and elution in 70 % acetonitrile/0.1 % formic acid according to manufacturer’s specifications. The eluate (40 μL) was evaporated to near dryness in an Eppendorf Centricon Plus vacuum centrifuge at 30° C over 1 h. The remaining 4 μL concentrated sample was diluted in 18 μL SEC Buffer (H_2_O:Acetonitrile:TFA at a 70 : 30:0.1 ratio) and the peptides were fractionated on a Superdex Peptide PC 3.2/200 size-exclusion column (equilibrated with the SEC buffer) with the SEC buffer at a flow rate of 0.05 mL/min on a GE Healthcare ÄKTA micro system. 75 μL fractions were collected in “ProtLoBind” microcentrifuge tubes and the fractions eluting between 0.4 and 0.6 column volume were pooled, evaporated to dryness and redissolved in 17 μL 0.5 % trifluoroacetic acid for the subsequent proteomics analysis.

The lysine-specific cross-linking of RNA granule proteins with a biotin-containing enrichable cross-linker Leiker (J & K Scientific) was based on the protocol of Tan and coworkers (Tan *et al*., 2016). A minimal amount of the cross-linker was dissolved in anhydrous DMSO (Invitrogen) immediately prior to use and the concentration of the cross-linker determined based on the A_260_ absorbance of the N-hydroxy-succinimidyl group using a molar extinction coefficient 9 700 M^-1^cm^-1^ (Miron & Wilchek, 1982). RNA granule preparation corresponding to 300 µg total protein was incubated with 0.23 mM Leiker at room temperature for 30 min protected from light. The cross-linking reaction was quenched by incubation with 90 mM Tris at room temperature for 20 min. The cross-linked proteins were precipitated with 4 volumes of acetone at – 20° C for 30 min, followed by centrifugation at 21 000 × g/4° C for 20 min, removal of the supernatant and evaporation of the residual solvent at 37° C for 4 min. The protein pellet was redissolved by incubation in 32 µL Denaturation Buffer at 50° C for 20 min. Cysteines were blocked with 10 mM iodoacetamide at room temperature for 30 min and any remaining iodoacetamide scavenged with 10 mM dithiothreiol at room temperature for 10 min. The sample was diluted 6.5-fold with 100 mM Tris-HCl (pH 8.5) and proteomics grade trypsin was added to the sample at an enzyme-to-substrate ration of 1 : 75, followed by an overnight digestion at 37° C in the presence of 1 mM CaCl_2_. The trypsinization was inhibited by the addition of PMSF to an f.c. of 0.1 mM and the sample diluted 2-fold with 20 mM Hepes-KOH (pH 8.0). The sample was subsequently incubated with 80 µL strept-actin resin (Strep-Tactin Superflow Plus; QIAGEN) at room-temperature for 2 h with constant agitation on a horizontal roller mixer. The resin with the bound peptides was pelleted by a quick spin and the supernatant was removed. The resin was successively washed with 3 × 500 µL 20 mM Hepes-KOH (pH 8.0)/1 M KCl, 1 × 500 µ L doubly-distilled water, 3 × 500 µ L 10 % acetonitrile in doubly-distilled water and finally with 3 × 500 µL doubly-distilled water. The cross-linked peptides were eluted from the resin by a dithionate-induced reductive cleavage of the azo group between the cross-linker and the biotin moiety by incubating the resin in 100 µL Cleavage Buffer (10 mM Hepes-KOH, pH 8.0, 6 M urea, 2 M thiourea, 0.3 M Na_2_S_2_O_4_) at 37° C for 30 min. The progress of the cleavage reaction was visually monitored by the disappearance of the yellowish color of the resin. The resin was briefly spun and the supernatant containing the cross-linked peptides withdrawn to a “ProtLoBind” tube. The resin was twice eluted with 50 µ L doubly-distilled water through a brief centrifugation and the supernatants combined with the original supernatant and sent for proteomics analysis.

### Mass-spectrometry analysis of cross-linked peptides

Peptides were injected to an Ultimate 3000 RSLCnano system (Dionex) using a 0.3 × 5 mm trap-column (5 µm C18 particles, Dionex) and an in-house packed (3 µm C18 particles, Dr Maisch) analytical 50 cm × 75 µm emitter-column (New Objective). Peptides were eluted at 250 nL/min with an 5-40% (4 h) A to B gradient (buffer A: 0.1% (v/v) formic acid; buffer B: 80% (v/v) acetonitrile + 0.1% (v/v) formic acid) to a quadrupole-orbitrap Q Exactive Plus (Thermo Fisher Scientific) MS/MS via a nano-electrospray source (positive mode, spray voltage of 2.5 kV). The MS was operated with a top-10 data-dependent acquisition strategy. Briefly, one 350-1,400 m/z MS scan at a resolution setting of R = 70,000 was followed by higher-energy collisional dissociation fragmentation (normalized collision energy of 26) of the 10 most intense ions (z: +2 to +6) at R = 17,500. MS and MS/MS ion target values were 3,000,000 and 50,000 ions with 50 ms injection times, respectively. Dynamic exclusion was limited to 70 s. MS .raw files were converted to .mgf format (top 100 peaks per 100 Da) using ProteoWizard MSConvertGUI (Chambers *et al*, 2012).

### Identification of cross-linked peptides using pLink

Cross-linked peptides were identified using pLink software as described previously (Yang et al., 2012), with the following modifications. Cross-linker was set to Leiker_clv, DSS, BS3 or BS(PEG)_5_. BS(PEG)_5_ was defined as having LinkerMass 302.36, MonoMass 320.38. The minimum peptide length was set to 5 amino acids, oxidation on Met was set as a variable modification, and carbamidomethyl Cys was set as fixed modification. Missed cleavage sites were chosen 3, PeptideMass 500 to 7000, PeptideLength 5 to 60, PrecursorTolerance 10 ppm, FragmentTolerance 20 ppm. False discovery rate was 5%. The search databases consisted of the Rattus norvegicus protein sequences that were identified previously by proteomics as nRNAg components (r-proteins, translation factors, RBP etc.). The sequences were downloaded from Uniprot. The resulted excel files of cross-linked specters are added in Supplementary Table S6.

### Nano-LC/MS/MS for protein identification

Samples were injected to an Ultimate 3000 RSLCnano system (Dionex) using a C18 trap-column (Dionex) and an in-house packed (3 µm C18 particles, Dr Maisch) analytical 50 cm x 75 µm ID emitter-column (New Objective). Peptides were eluted at 250 nl/min with a 5-35% B 120 min gradient (buffer B: 80% acetonitrile + 0.1% formic acid, buffer A: 0.1% formic acid) to a Q Exactive Plus (Thermo Fisher Scientific) mass spectrometer (MS) using a nano-electrospray source (spray voltage of 2.5 kV). The MS was operated with a top-10 data-dependent acquisition strategy. Briefly, one 350-1400 m/z MS scan at a resolution setting of R=70 000 at 200 m/z was followed by higher-energy collisional dissociation fragmentation (normalized collision energy of 27) of 10 most intense ions (z: +2 to +6) at R=17 500. MS and MS/MS ion target values were 3e6 and 5e4 with 50 ms injection times. Dynamic exclusion was limited to 40 s.

### LC/MS/MS raw data processing

Mass spectrometric raw files were analyzed the with the MaxQuant software (version 1.6.15.0) (Cox & Mann, 2008). The methionine oxidation, was set as variable modifications. Cysteine carbamidomethylation was defined as a fixed modification in both searches. Searches were performed against the UniProt (www.uniprot.org) *Rattus norvegicus* or *Homo sapiens* reference proteome database using the tryptic digestion rule (including cleavages after proline). Transfer of identifications between runs was enabled. iBAQ feature was also enabled, which normalizes protein intensities by the number of theoretically observable peptides and enables rough intra-sample estimation of protein abundance. Peptide-spectrum match and protein false discovery rate (FDR) was kept below 1% using a target-decoy approach. All other parameters were default.

### Protein abundance calculation based iBAQ intensity value

To estimate the ribosome-to RBP stoichiometry/ratio, the MS data in Supplementary Table S1 was first used to calculate the average iBAQ intensity value for the ribosome. For this, majority of r-proteins representing the most abundant population was selected and average iBAQ value calculated in Excel. The r-proteins that were excluded from the calculations deviated over 2-fold from the mean iBAQ value. Excluded r-proteins from CH-tissue and SH-SY5Y are listed in Supplementary Table S7. The average of the iBAQ intensities of the remaining r-proteins was set equal to 1 equimole of 80S ribosome. The ribosome-to-RBP stoichiometry was then calculated by dividing the „ribosome iBAQ“ value by the iBAQ value of a given RBP in each replicate

For the rankordered r-protein lists in Supplementary Table S3, the relative abundance of each identified r-protein was calculated by normalizing its individual iBAQ value to the mean over the iBAQ values of all identified r-proteins. For Supplementary Figure S5 and S6, the relative abundances for the 40S and 60S subunit r-proteins were calculated separately from the iBAQ data of Waterbeemd et al. 2018 (i.e. the iBAQ value of a 40S r-protein was normalized to the mean over the iBAQ values of all 40S r-proteins and likewise for the 60S r-proteins).

In the original MS data (reported in Supplementary Table S1), some identified r-proteins (Rps2, Rps3, Rps12, Rps18, Rps20, Rps24, Rps25, Rpl10, Rpl21, and Rpl36) were present in duplicates with one of the duplicates having only 1 or 2 corresponding razor + unique peptides. Out of those duplicates, the entry with 1 or 2 peptides was excluded from the subsequent analysis.

### Protein Purification

G3BP-1, G3BP-2, and Caprin-1 were expressed using a pET-derived expression vector (Addgene, pET His6 MBP Asn10 TEV LIC, 1C) in *E.coli* Rosetta as N-terminal fusions to an N-terminally His6-tagged *E.coli* maltose-binding protein (MBP). An N-terminally His6-tagged variant of MBP was expressed from the pET-derived expression vector identically to the fusion proteins. For protein expression, cells were grown in 1 L 2 × YT medium at 37° C under constant shaking at 140 rpm. Upon reaching a mid-log phase (OD_600_ ≈ 0.6) IPTG was added to the cultures to an f.c. of 1 mM and the incubation continued for 3 h. Cells were harvested by centrifugation in a Herolab A6.9 rotor at 2000 × g/4 C for 10 min. The supernatant was decanted and the cell pellet flash-frozen in liquid nitrogen and stored at – 80 C until further processing. The cells were thawed on ice and resuspended in 35 mL ice-cold lysis buffer (40 mM Hepes-KOH, pH 7.5, 500 mM KCl, 10 mM imidazole, 2 mM beta-mercaptoethanol) in the presence of 0.1 mM PMSF and a protease inhibitor cocktail (cOmplete, EDTA-free; Roche). Triton X-100 (f.c. 0.1 % (v/v)), lyzozyme (f.c. 1 mg/mL) and DNase I (f.c 43 U/mL) were added to the suspension and the suspension was incubated at 4° C on an end-over-end shaker for 15 min. The cells were subsequently lyzed by sonication in a Bandelin Sonoplus sonicator at 55 % power through 10 cycles (10 s each) on ice and the lysate was clarified by centrifugation in a Sorval SS-34 rotor at 34 540 × g/4° C for 40 min. The lysate supernatant was additionally filtered through a 0.22 µm syringe filter. All subsequent chromatographic procedures were performed at 4° C.

With G3BP-1, G3BP-2, and Caprin-1, the clarified lysate was applied to a Ni-Sepharose affinity column (HisTrap^TM^ 5 ml; Cytiva) at a flow rate of 1 mL/min using a peristaltic pump. After washing the column with 3 column volumes of HisTrap buffer [20 mM Hepes-KOH, pH 7.5, 500 mM KCl (for G3BP-1, Caprin-1)/800 mM KCl (for G3BP-2), 20 mM imidazole, 2 mM beta-mercaptoethanol] at a flow rate of 1 mL/min, the column was connected to a GE Healthcare ÄKTA Purifier system and washed with additional 3 column volumes of HisTrap buffer at a flow rate of 1 mL/min. Bound protein was eluted from the Ni-Sepharose column using a linear gradient of imidazole in HisTrap buffer up to 300 mM (for G3BP-1, Caprin-1) or 200 mM (G3BP-2) at a flow rate of 2 mL/min. Fractions corresponding to the major peak were pooled and applied to an MBP affinity column (MBPTrap^TM^ 10 ml; Cytiva) at a flow rate of 2 mL/min. After washing the MBP column with 3 column volumes of MBP buffer [20 mM Hepes-KOH, pH 7.5, 500 mM KCl (for G3BP-1, Caprin-1)/600 mM KCl (for G3BP-2), 2 mM beta-mercaptoethanol], bound proteins were eluted with 10 mM D-maltose in MBP buffer at a flow rate of 2 mL/min. The eluted fractions were pooled and incubated with TEV protease at an enzyme-to-substrate ratio ranging from 1 : 100 to 1 :150 at 4° C overnight on an end-over-end shaker. G3BP-1, G3BP-2 and Caprin-1 were purified from MBP by passing the TEV cleaved sample through the Ni-Sepharose column at 20 mM imidazole in HisTrap buffer at a flow rate of 2 mL/min. Under those conditions the non-tagged proteins elute in the low imidazole buffer while the N-terminally His_6_-tagged MBP is retained on the column.

G3BP-1 containing fractions from the second (subtractive) Ni-Sepharose affinity chromatography were pooled and concentrated by centrifugation on a 30 kDa MWCO Amicon Ultra centrifugal filter unit at 3220 × g/4° C. KCl in the retentate was diluted to 50 mM with 20 mM Hepes-KOH (pH 7.5). The G3BP-1 containing retentate was subsequently fractionated by cation exchange on a sulphopropyl column (HiScreen HP; Cytiva) at a flow rate of 0.5 mL/min using a stepwise KCl gradient (from 50 mM to 1 M KCl) in SP buffer [20 mM Hepes-KOH (pH 7.5), 2 mM beta-mercaptoethanol]. Two partially overlapping peaks eluted at 140 – 200 mM KCl, followed by a single large peak at 300 mM KCl. According to an SDS-PAGE analysis, the peaks eluting at 140 – 200 mM KCl correspond to a partially cleaved G3BP-1 while the full-length protein is mostly found in the peak eluting at 300 mM KCl. Fractions corresponding to the 300 mM KCl peak were pooled and concentrated by centrifugation at 3200 × g/4° C on an Amicon Ultra Centrifugal filter unit. The concentrated sample was fractionated on a Sephacryl S200 size exclusion column in S200 buffer [20 mM Hepes-KOH (pH 7.5), 500 mM KCl, 2 mM beta-mercaptoethanol] at a flow rate of 0.4 mL/min. During the fractionation, only a broad peak eluting in the void volume (0.15 – 0.3 CV) was observed, likely due to an aggregation of G3BP-1. The presence of G3BP-1 in this peak was confirmed by an SDS-PAGE analysis. Fractions corresponding to the peak were pooled, concentrated by centrifugation at 3200 × g/4° C on an Amicon Ultra Centrifugal filter unit and stored at – 80° C in a buffer containing 20 mM Hepes-KOH, pH 7.5, 131 mM KCl, 10 % (v/v) glycerol and 1.75 mM beta-mercaptoethanol.

G3BP-2 containing fractions from the second Ni-Sepharose chromatography were pooled and concentrated by centrifugation on a 30 kDa MWCO Amicon Ultra centrifugal filter unit at 3220 × g/4° C. The G3BP-2 containing retentate was fractionated on a HiLoad 16/600 Superdex 200pg size-exclusion column and the fractions corresponding to G3BP-2 were pooled, concentrated by centrifugation at 3200 × g/4 C on an Amicon Ultra Centrifugal filter unit and stored at – 80° C in a buffer containing 20 mM Hepes-KOH, pH 7.5, 514 mM KCl, 2 % (v/v) glycerol, 5.5 % (v/v) trehalose, 1.7 mM beta-mercaptoethanol and 0.14 mM TCEP.

Caprin-1 containing fractions from the second Ni-Sepharose affinity chromatography were pooled, buffer exchanged for S200 buffer (20 mM Hepes-KOH, pH 7.5, 500 mM KCl, 2 mM beta-mercaptoethanol) on a HiTrap 26/10 desalting column and concentrated by centrifugation on a 30 kDa MWCO Amicon Ultra centrifugal filter unit at 3220 × g/4 C. The preparation was stored at – 80° C in a buffer containing 20 mM Hepes-KOH (pH 7.5), 500 mM KCl, 40 % (v/v) glycerol and 2 mM beta-mercaptoethanol.

For maltose-binding protein (MBP) purification, the clarified lysate was fractionated on a Ni-Sepharose column as described above. Fractions containing MBP were pooled and buffer-exchanged for Q buffer [20 mM Hepes-KOH (pH 7.5), 50 mM KCl, 2 mM beta-mercaptoethanol] on a HiTrap 26/10 desalting column at 1.5 mL/min. The MBP-containing fractions were pooled and further purified on an HiScreen Q HP anion-exchange column (Cytiva) at a flow rate of 0.5 mL/min in a 0.05 – 1 M KCl linear gradient in Q buffer. Fractions containing MBP were pooled and buffer-exhanged for B buffer [20 mM Hepes-KOH (pH 7.5), 150 mM KCl, 2 mM beta-mercaptoethanol] by centrifugation at 3200 × g/4° C on an Amicon Ultra Centrifugal filter unit. and stored at – 80° C in a buffer containing 20 mM Hepes-KOH (pH 7.5), 131 mM KCl, 10 % (v/v) glycerol and 1.75 mM beta-mercaptoethanol.

All protein concentrations were measured via 10-fold dilutions in 6 M GuHCl using known extinction coefficients. Before storage, all proteins were aliquoted and flash-frozen in liquid nitrogen.

### Sample preparation for cryo-EM

Two different nRNAg samples were made for cryo-EM. Both samples were made of nRNAg purified from rat cortical-hippocampal tissue. Samples were diluted into the concentration of 2 U/ml (A_260_) in Buffer A (20 mM HEPS, pH7.5/150 mM KOAc/ 4 mM MgOAc/0.5 mM TCEP). The only difference was that one sample was treated with RNase T1 0.03 (U/ μl) for 30 min at RT and other not. 3 μl of sample was applied onto glow-discharged C-Flat 1.2/1.3- pp) made in-house. Cryo-EM grids were vitrified using a FEI Vitrobot Mark IV at 10°C and 100% humidity. Sample was incubated 150 sec on the grid, and blotted with the force of 5 for 1 to 2 sec. After blotting the grids were plunged into liquid ethane at 93 K and subsequently into liquid nitrogen for storage.

### Cryo-EM data collection and processing

Two independent data sets were collected on Talos Arctica or Titan Krios electron microscopes (ThermoFisher) equipped with Falcon III direct electron detector (FEI). Intact nRNAg dataset was collected on Talos Arctica operated at 200kV at a magnification of 92 000x and at pixel sizes of 1.62 A_/pix. 20 frames per movie were collected for 1.2 s in integrated mode with defocus range from −1 μm to -3.0 μm, and flux of 22 e/A□^2^/s. 70 μm objective aperture was used.

RNase T1 treated dataset was collected on Titan Krios (Diamond cryo-EM facility, eBIC) Falcon III detector, and EPU software was used for data-collection. Nominal magnification 59kx that gave 1.41 A /pix, and 4 shots per hole. 39 frames per movie, dose per frame 1.2 e/ A□^2^. In total, 3644 micrographs were collected. Relion 3.1 software (2) was used to process the single-particle cryo-EM data (Scheres, 2012). Motion correction was performed using RELION 3’s own implementation (Zivanov *et al*., 2018). Movies were aligned using 5×5 patches with dose-weighting. CTF was estimated using CTFFIND4 (Rohou & Grigorieff, 2015). The cryo-EM map of human 80S ribosome (PDB id: 6ole) (Li *et al*., 2019) was used as reference map after lowpass filtering to 70 A□ for the initial 3D classification. After focused classification on various regions of the map and 3D refinement, Bayesian polishing in RELION was used for beam-induced motion correction (Zivanov *et al*, 2019). All reported resolutions are based on the gold-standard Fourier shell correlation (FSC) = 0.143 criterion (Chen *et al*, 2013)

After 2D classification and multiple 3D classifications (Figure 8), 3D class of 62 369 particles was selected for further data-processing. After unmasked and masked refinement, the resolution at FSC 0.143 was reported to be 2.95 Å (Supplementary Figure S11B). Per-particle, reference-based beam-induced motion correction step improved resolution to 2.87 Å.

The predominant molecular motions were accounted by multi-body refinement and flexibility analysis in RELION (Nakane *et al*., 2018; Nakane & Scheres, 2021). For the multi-body refinement, consensus map determined in this study was used to prepare the bodies using Chimera Segger segmentation tool (Pettersen *et al*, 2004; Pintilie *et al*, 2010) as described by Nakane, and Scheres, 2021. Ribosomes were segmented into two bodies (40S and 60S), or three bodies (40S-head, 40S-body, and 60S). Bodies/segmentations were saved as .mrc maps in Chimera and Relion MaskCreate tool was used to generate masks with Extended binary map by 3 pixels and Soft-edge by 6 pixels. The body STAR file was made to determine the movement relationships between various bodies. For the two-body refinement, angular priors were set 5 degrees for 60S and 25 degrees for 40S. The widths of the translation priors were set 2 pixels for 60S and 5 pixels for 40S.

For the three-body refinement angular priors were set 5 degrees for 60S, 20 degrees for 40S-body, and 20 degrees for 40S-head. The widths of the translation priors were set 2 pixels for 60S, 5 pixels for 40S-body and 40S-head. The body reconstructions were analysed using Chimera and “Volume Series” tool was used to analyse the principal motions to prepare the movies. The three cycles of each principal motion were recorded and saved as .mp4 file. Chimera segmentation tool was used to color-code the maps for movies. Movie editing software VEED.IO was used to add labels to movies.

### Molecular model determination

The Phenix real-space refinement was used to determine the molecular structure of ribosome bound to nRNAg (Afonine *et al*, 2018). The human cryo-EM structure of (PDB: 6ole) was used as initial model to build the rat 80S ribosome structure. Secondary structure restrains were applied during refinement. In order to construct rat ribosome structure, r-protein sequences were aligned between human and rat. Sequences were adopted from Uniprot and NCBI global protein blast suit was used. The list of changes made to human ribosome proteins are in Supplementary Table S7. Coot software was used to manually insert the differences into coordinates and real-space refined (Emsley *et al*., 2010). Model was evaluated using MolProbity (Chen *et al*, 2010). Maps and models were visualized using Pymol (version 2.3.2) or Chimera software (Pettersen *et al*., 2004).

### rRNA sequencing

In order to sequence variable regions of ribosomal 18S rRNA and 28S rRNA, the rRNA were extracted from Wistar rat cerebral cortices neuronal RNA granules using QIAGEN RNeasy mini-kit as stated in manufacturing protocol. A mixture containing reverse transcription primer, reaction buffer (50 mM Tris-HCl, pH 8.3, 50 mM KCl, 10 mM DTT), 1 µL of RNA (100ng/µL) was incubated at 70◦C for 5 minutes. Following incubation step, dNTPs at a final concentration of 1 mM, 50 mM MgCl_2_, *Solis BioDyne^TM^* FIRESCRIPT enzyme mix were added to the mixture, and the reverse transcription was performed under the conditions shown in the Supplementary Table S9. The obtained cDNA were then amplified by PCR using *Thermo Scientific Phusion High-Fidelity DNA Polymerase*, different primers were used for amplification (primer list below), only the regions that were known to have differences in nucleotides between human rRNA and rat rRNA molecules were amplified and sequences (18S rRNA regions: 41-414, 340 - 937, 733 - 1275; 28S rRNA: 211 - 574, 1294 - 1905, 3113 - 3899, 4362 - 4548). The PCR products were applied on 1% TAE agarose gel. DNA was extracted using Thermo scientific Gene jet gel extraction kit according to the protocol provided. The samples were sent for sequencing in Core Facility of Genomics, University of Tartu and the sequencing results were obtained and analyzed using Ape (plasmid editor) or SnapGene® software (from Insightful Science; available at snapgene.com). The human 28S rRNA sequence (Homo sapiens RNA, 28S ribosomal N2, ribosomal RNA, NCBI Reference Sequence: NR_146148.1) and rat (Rattus norvegicus 28S ribosomal RNA, ribosomal RNA, NCBI Reference Sequence: NR_046246.2) were used as reference sequences in an alignment. The human 18 S rRNA sequence (Homo sapiens RNA, 18S ribosomal N2, ribosomal RNA, NCBI Reference Sequence: NR_146146.1) rat 18S rRNA sequence (Rattus norvegicus 18S ribosomal RNA, ribosomal RNA, NCBI Reference Sequence: NR_046237.2). The 18S rRNA sequence was identical to the reference rat 18S rRNA in sequenced regions.

### The following list of primers were used for cDNA amplification

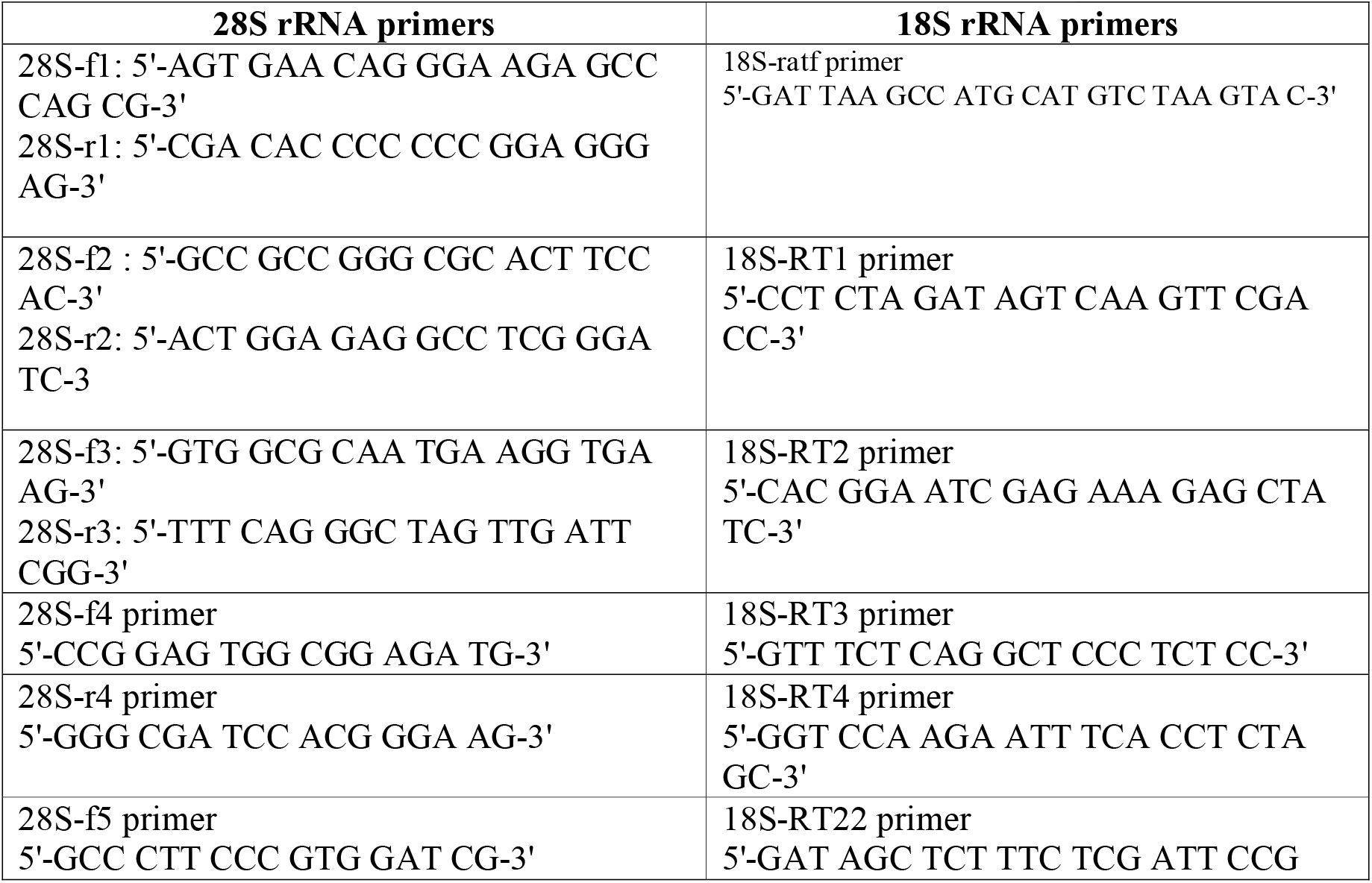
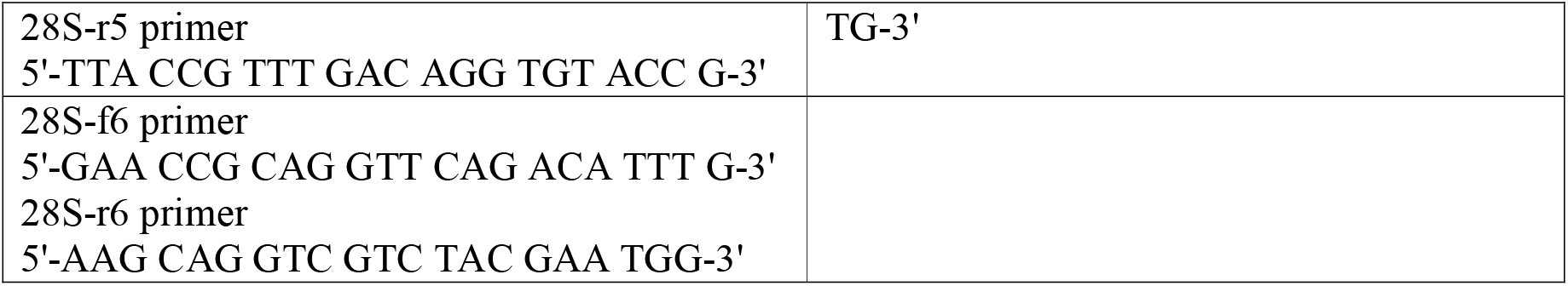

### The following list of primers were used for reverse transcription

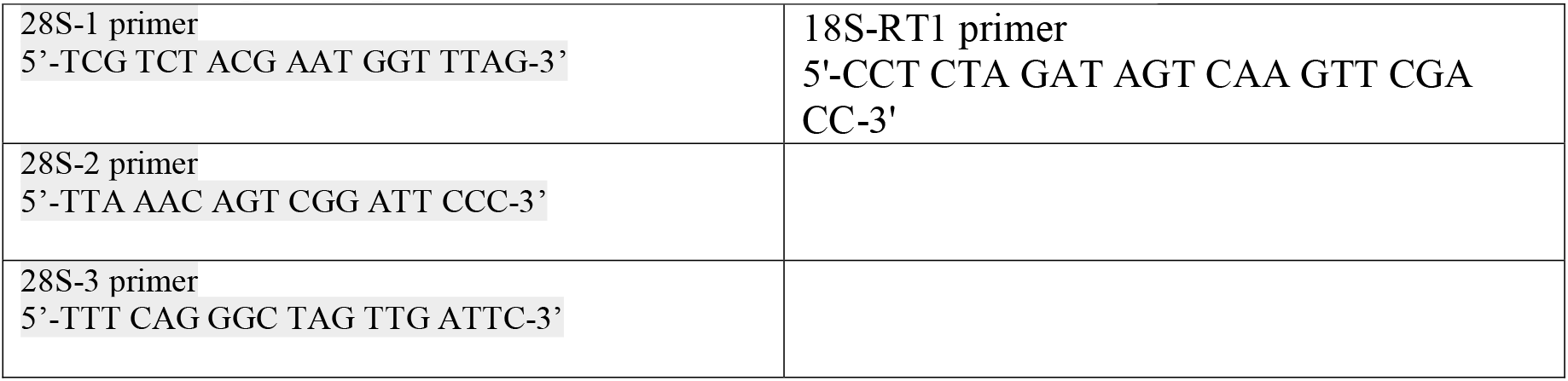

## Ethical standards

The experiments comply with the current laws of Estonia, where they were performed.

## Acknowledgment

We thank Dr. Sergo Kasvandik, and Merilin Saarma for advice and support for mass spectrometry analysis in Proteomics core facility in University of Tartu; EMBL Proteomics core facility for mass-spectrometry data collection and analysis. We also thank Dr. Allen Kaasik, Marko Piirsoo and Ulla Peterson for the advice and support for rat brain tissue dissection. We appreciate the collaboration with Dr. Alexy Amunts lab in Stockholm University and Victor Tobiasson for collecting intact RNA granule cryo-EM data. Dr. Alistair Siebert in the Diamond eBIC cryo-EM facility for data collection that was supported by iNext PID8541 program; Dr. Kärt Padari for electron microscopy support at University of Tartu. We are grateful to Dr. Ülo Maiväli for his thoughtful comments and data analysis advice of the manuscript. This work was supported by Estonian Research Council (grant PSG87 and MOBTP44), and by the EMBO installation grant Nr. 3911. Data and materials availability: EM maps have been uploaded to the Electron Microscopy Data Bank with accession code EMD-13954 (RNase T1 treated) and EMD-13975 (intact). Protein coordinate has been deposited in the Protein Data Bank (PDB) with ID 7QGG. Mass-spectrometry coupled cross-linking data is deposited to ProXL database and can be accessed using URL: https://www.yeastrc.org/proxl_public/viewProject.do?project_id=527

## Author contributions

KK and AP performed rat brain dissection, purification of neuronal RNA granules, negative staining EM experiments, and protein purification. KK perfomed the cell culture experiments. KK and AM performed chemical cross-linking experiments. AP prepared the cryo-EM grids. AP performed cryo-EM data analysis. AP conducted the cross-linking data analysis using pLink. AM performed rRNA sequencing assay. AP and KK prepared the manuscript, with contributions and edits from AM.

---

Supplementary Table S5. pLink software was used to process the mass-spectrometry CL data (Yang *et al*, 2012).

Supplementary Figure S1. Neuronal RNA granules contaminated with larger polysomes. (A) Pellet-1 fraction of rat cortex-hippocampus tissue imaged by ns-EM. The sample contains spherical tightly packed nRNAg’s and polysomes. (B) nRNAg fraction of rat cortex-hippocampus tissue imaged by ns-EM. Pellet-1 fraction is enriched with nRNAg’s by Sephacryl-S500 chromatography step. The scale bar on the bottom corner represents 500 nm distance.

Supplementary Figure S2. Nuclease sensitivity of cortex-hippocampus RNA granules. RNA granules were incubated with MNase or RNase T1 at room temperature for 30 min at 150 mM monovalent salt and deposited on glow-discharged continuous carbon grids for negative staining. Scale bar – 500 nm (A – C, E - F), 200 nm (D).

Supplementary Figure S3. The RNA granules are detectable in the lysate of rat CH tissue without the pelleting step. (A to C) The lysate of the rat CH tissue was partially purified by using Sephacryl-S500 spin-column. The larger particles (including nRNAg) will move through the resin and no pelleting occurs. The dense particles visible in ns-EM images resemble the purified nRNAg’s. The scale bar on the bottom corner represents 500 nm distance. (D) The nRNAg’s purified from the rat CH tissue using two-layer (90 % and 60%) sucrose cushion. The CH tissue lysate was loaded on the two-layer sucrose cushion and the nRNAg material was collected in the interface of two sucrose layers to avoid pelleting the sample. The collected fraction was concentrated using stirred cell (Merck Millipore) that keeps particles in dissolved state. ns-EM images show the presence of nRNAg particles. The scale bar on the bottom corner represents 200 nm distance.

Supplementary Figure S4. Negative stain TEM images of recombinant human G3BP1 purified from E. coli (A-B) 200 nM G3BP1 sample was loaded onto a continuous carbon coated Quantifoil TEM grid and visualized by negative staining. G3BP1 tends to form liquid-like droplets characteristic of various RBP’s. (C) 200 nM MBP-G3BP1 (N-terminally fused G3BP1 with MBP). Liquid –like droplet formation is diminished when an MBP-tag is added to G3BP1. The scale bar on the bottom corner represents 200 nm distance.

Supplementary Figure S5. Relative abundance of r-proteins in rat corticohippocampal RNA granules and mouse embryonic stem cells. Data for mouse embryonic stem cells is from Slavov et al. 2015. The green zone corresponds to r-proteins whose relative abundance relative to the mean does not exceed 2-fold.

Supplementary Figure S6. Relative abundance of r-proteins in rat corticohippocampal RNA granules, SH-SY5Y RNA granules and HEK293-6E 80S ribosomes. Data for the HEK293-6E 80S ribosomes is from Supplementary Data 1 of Waterbeemd et al. 2018. The green zone corresponds to r-proteins whose relative abundance with respect to the mean does not exceed 2-fold.

Supplementary Figure S7. Analysis of RBPs in RNA granules by Western blot. The presence of G3BP-1, G3BP-2, and Caprin-1 in cortex-hippocampus RNA granules (0.13 – 0.15 A260 units/lane) was analyzed by immunoblotting using RBP-specific antibodies. Negative staining EM micrographs of the RNA granules used for the immunoblotting are shown. Scale bar – 200 nm.

Supplementary Figure S8. Correlation between r-protein abundances. The number on the figure indicates the Pearson correlation coefficient between two datasets. (A) r-protein abundances in rat corticohippocampal RNA granules versus SH-SY5Y RNA granules. (B) r-protein abundances in rat corticohippocampal RNA garnules versus HEK293-6 ( 80S ribosomes. Panel (C) r-protein abundances in SH-SY5Y RNA granules versus HEK293 80S ribosomes.

Supplementary Figure S9. Nonspecific binding of RNA granule components to Ni-Sepharose affinity resin. MBP-coated Ni-Sepharose resin incubated with 1.4 A_260_ units cortex-hippocampus RNA granules was washed with RNA Granule Buffer and incubated with SDS Gel Loading Buffer at 95 C for 5 min to elute nonspecifically bound proteins. The eluted proteins were analyzed by immunoblotting using Caprin-1 and G3BP-1-specific antibodies.

Supplementary Figure S10. Structural validation of the cross-linking data on the ribosome complexes. (A) Ttc5 (blue) cross-link with uL24 (red) is modeled on the cryo-EM structure of Ttc5 bound on ribosome (PDB; 6t59). Cross-linking distances between 10 to 30 Å can be expected with BS(PEG)_5_ chemical cross-linker. Dashed line represents the distance in angstroms (Å) between Ca atoms. (B) Pa2G4 (EBP1) (blue) cross-link with uL29 (red) is modeled on the cryo-EM structure of Pa2G4 bound on a ribosome (PDB; 7bhp). (C) Btf3 (blue) cross-link with eL31 (red) is modeled on the cryo-EM structure of Btf3 bound on a ribosome (PDB; 6t59). We identified CL with Btf3 residue K13 with eL31 (K70 and K91), but Btf3 structure is missing N-terminal region, and therefor the most N-terminal residue (V53) is modeled.

Supplementary Figure S11. nRNAg cryo-EM data-processing workflow. (A) 2D classification results using Relion softwar. The image shows all the 2D classes (bad classes included). Well aligned 2D classes (74 classes, containing 462 031 particles) were selected, and further classified by 3D classification. (B) FSC curve of a cryo-EM map after polishing step. The gold-standard FSC at 0.143 gives average resolution estimate of 2.87 Å.

Supplementary Figure S12. 3D classification scheme of the RNase T1 treated nRNAg particles purified from rat cortex-hippocampus tissue. A total of 425 778 particles were selected from the 2D classification step using Relion software (Scheres, 2012). In the first 3D classification step, particles were classified into 4 classes. The particle numbers in each class are indicated on the top of the class. The second 3D class (CL2) represents the consensus map that was chosen as a reference map to fit the rest of the maps using Chimera “Fit in Map” tool (Pettersen *et al*., 2004). The map correlation values are indicated on the left on each class where second class has value 1.0, meaning perfect correlation. Class 1 was classified into 8 classes, and additional classification runs were made to separate any interesting classes. For example, CL1-CL4 was classified into 8 separate classes, and CL1-CL5 into 4 classes.

Supplementary Figure S13. 18S rRNA alignment between rat and human origin. SnapGene local-alignment (Smith-Waterman) was used with default parameters. Identical bases are indicated with “dot” and differences with “base” substitution.

Supplementary Figure S14. 28S rRNA alignment between rat and human. SnapGene multiple-dna sequence alignement (MUSCLE) tool was used with default parameters. “Rat 28S seq” sequence represents modified rat 28S rRNA sequence acquired by sequencing various regions of 28S rRNA purified from nRNAg. The sequences that were not sequences are also included in the “Rat 28S seq” sequence, see Materials and Methods.

Supplementary Figure S15. Effect of translation inhibitors on RNA granules. Negative staining EM micrographs of cortex-hippocampus RNA granules isolated in the presence of translation inhibitors homoharringtonine (B) or cycloheximide (C). The inhibitors were present throughout tissue extraction and cell lysis. Panel A – RNA granules isolated in the absence of inhibitor. Sacle bar – 200 nm (A,B) or 500 nm (C).

Supplementary Table S1. Mass-spectrometry analysis of the nRNAg sample purified from Wistar P0 (biological replicate 1, sheet 1), P4 (biological replicate 2, sheet 2) puppies or from human neuroblastoma cell line SH-SY5Y (biological replicate 1 and 2, sheets 3 and 4, respectively). The protein groups are sorted using iBAQ intensities. Most abundant proteins are ribosomal proteins and following RBPs, like Caprin-1 and G3BP2.

Supplementary Table S2. List of uncharacterized proteins identified by proteomic analysis of nRNAg’s purified from rat cortices. Identity was evaluated based NCBI Protein BLAST alignment.

Supplementary Table S3. Relative abundances of r-proteins in rat and SH-SY5Y RNA granules. The proteins have been rankordered according to their relative abundance (defined as the ratio of the iBAQ value of an individual r-protein to the mean over the iBAQ values of all identified r-proteins). The data represents averages of two replicates for the rat and SH-SY5Y RNA granules, respectively. Low-abundance r-proteins shared between the rat and SH-SY5Y datasets have been highlighted in red. To facilitate comparison with previously published data, the relative abundances are presented in log2 units.

Supplementary Table S4. Relative abundance of RBPs in nRNAg purified from rat cortex-hippocampus tissue or human neuroblastoma SH-SY5Y. The relative abundance values were calculated based on the iBAQ intensity values listed in Supplementary Table S1. Ribosomal proteins average iBAQ intensity value was taken as reference and was assigned 1 equimole. The relative abundance of RBPs were calculated with respect to the ribosome. Calculations were done based on 2 biological replicates and standard deviations was calculated.

Supplementary Table S5. Non-specific binding of RNAg components to the MBP-Ni-Sepharose resin. Rat brain nRNAg sample was incubated with MBP-coated Ni-Sepharose resin and the nonspecifically bound proteins were analyzed by mass-spectrometry after elution of the resin in 8 M urea. The protein groups are sorted using iBAQ intensities. In addition to MBP, nonspecifically bound RBPs and r-proteins are also present in abundance in the urea-eluted fraction.

Supplementary Table S6. pLink (Yang *et al*., 2012) results of cross-linking coupled mass-spectrometry analysis of nRNAg’s purified from rat CH tissue. Amine-reactive cross-linkers BS(PEG)_5_, DSG, DSS, and Leiker were used. Biological replicates with different cross-linkers are listed in excel sheets. Summary of the cross-linking results is in the Table 1.

Supplementary Table S7. List of r-proteins that were excluded from the calculations of average iBAQ intensity value for ribosome. Two biological replicates from rat CH-tissue or neuroblastoma are indicated.

Supplementary Table S8. Ribosomal protein primary sequence differences between human and Rattus norvegicus that were modeled into structure. Listed are only the r-proteins and residues that exist in the structure. Old and new nomenclature of r-protein names are listed (Ban *et al*, 2014).

Supplementary Table S9. Reaction conditions for reverse transcription and subsequent PCR of rRNA purified from rat CH nRNAg.

## References

Abraham WC, Williams JM (2008) LTP maintenance and its protein synthesis-dependence. Neurobiol Learn Mem 89: 260–268

Afonine PV, Poon BK, Read RJ, Sobolev OV, Terwilliger TC, Urzhumtsev A, Adams PD (2018) Real-space refinement in PHENIX for cryo-EM and crystallography. Acta Crystallogr D Struct Biol 74: 531–544

Agirrezabala X, Lei J, Brunelle JL, Ortiz-Meoz RF, Green R, Frank J (2008) Visualization of the hybrid state of tRNA binding promoted by spontaneous ratcheting of the ribosome. Mol Cell 32: 190–197

Alam U, Kennedy D (2019) G3BP1 and G3BP2 regulate translation of interferon-stimulated genes: IFITM1, IFITM2 and IFITM3 in the cancer cell line MCF7. Mol Cell Biochem 459: 189–204

Alami NH, Smith RB, Carrasco MA, Williams LA, Winborn CS, Han SSW, Kiskinis E, Winborn B, Freibaum BD, Kanagaraj A et al (2014) Axonal transport of TDP-43 mRNA granules is impaired by ALS-causing mutations. Neuron 81: 536–543

Alves MJ, Goksel M, Kaya B, Mostafa H, Gygli P, Stephens J, Fair S, Otero JJ, Czeisler CM (2019) CCNA2 Ablation in Aged Mice Results in Abnormal rRNA Granule Accumulation in Hippocampus. The American journal of pathology 189: 426–439

Anadolu MN, Kailasam S, Simbriger K, Sun J, Markova T, Jafarnejad SM, Lefebvre F, Ortega J, Gkogkas CG, Sossin WS (2021) Ribosomes in RNA granules are stalled on mRNA sequences that are consensus sites for FMRP association. bioRxiv: 2021.2002.2022.432349

Anderson P, Kedersha N (2009) Stress granules. Curr Biol 19: R397–398

Anger AM, Armache JP, Berninghausen O, Habeck M, Subklewe M, Wilson DN, Beckmann R (2013) Structures of the human and Drosophila 80S ribosome. Nature 497: 80–85

Arike L, Valgepea K, Peil L, Nahku R, Adamberg K, Vilu R (2012) Comparison and applications of label-free absolute proteome quantification methods on Escherichia coli. J Proteomics 75: 5437–5448

Aschrafi A, Cunningham BA, Edelman GM, Vanderklish PW (2005) The fragile X mental retardation protein and group I metabotropic glutamate receptors regulate levels of mRNA granules in brain. Proceedings of the National Academy of Sciences of the United States of America 102: 2180–2185

Balasanyan V, Arnold DB (2014) Actin and myosin-dependent localization of mRNA to dendrites. PLoS One 9: e92349

Ban N, Beckmann R, Cate JH, Dinman JD, Dragon F, Ellis SR, Lafontaine DL, Lindahl L, Liljas A, Lipton JM et al (2014) A new system for naming ribosomal proteins. Curr Opin Struct Biol 24: 165–169

Bandeira F, Lent R, Herculano-Houzel S (2009) Changing numbers of neuronal and non-neuronal cells underlie postnatal brain growth in the rat. Proceedings of the National Academy of Sciences of the United States of America 106: 14108–14113

Bassell GJ, Warren ST (2008) Fragile X syndrome: loss of local mRNA regulation alters synaptic development and function. Neuron 60: 201–214

Bassett DS, Wymbs NF, Rombach MP, Porter MA, Mucha PJ, Grafton ST (2013) Task-based core-periphery organization of human brain dynamics. PLoS Comput Biol 9: e1003171

Batish M, van den Bogaard P, Kramer FR, Tyagi S (2012) Neuronal mRNAs travel singly into dendrites. Proceedings of the National Academy of Sciences of the United States of America 109: 4645–4650

Belelovsky K, Elkobi A, Kaphzan H, Nairn AC, Rosenblum K (2005) A molecular switch for translational control in taste memory consolidation. Eur J Neurosci 22: 2560–2568

Bentmann E, Haass C, Dormann D (2013) Stress granules in neurodegeneration--lessons learnt from TAR DNA binding protein of 43 kDa and fused in sarcoma. The FEBS journal 280: 4348–4370

Besford QA, Sullivan MA, Zheng L, Gilbert RG, Stapleton D, Gray-Weale A (2012) The structure of cardiac glycogen in healthy mice. Int J Biol Macromol 51: 887–891

Bhaskar V, Desogus J, Graff-Meyer A, Schenk AD, Cavadini S, Chao JA (2021) Dynamic association of human Ebp1 with the ribosome. RNA 27: 411–419

Bidet K, Dadlani D, Garcia-Blanco MA (2014) G3BP1, G3BP2 and CAPRIN1 are required for translation of interferon stimulated mRNAs and are targeted by a dengue virus non-coding RNA. PLoS Pathog 10: e1004242

Blanchard SC, Kim HD, Gonzalez RL, Jr., Puglisi JD, Chu S (2004) tRNA dynamics on the ribosome during translation. Proceedings of the National Academy of Sciences of the United States of America 101: 12893–12898

Bosco DA, Lemay N, Ko HK, Zhou H, Burke C, Kwiatkowski TJ, Jr., Sapp P, McKenna-Yasek D, Brown RH, Jr., Hayward LJ (2010) Mutant FUS proteins that cause amyotrophic lateral sclerosis incorporate into stress granules. Human molecular genetics 19: 4160–4175

Bounedjah O, Desforges B, Wu TD, Pioche-Durieu C, Marco S, Hamon L, Curmi PA, Guerquin-Kern JL, Pietrement O, Pastre D (2014) Free mRNA in excess upon polysome dissociation is a scaffold for protein multimerization to form stress granules. Nucleic Acids Res 42: 8678–8691

Bramham CR, Wells DG (2007) Dendritic mRNA: transport, translation and function. Nature reviews Neuroscience 8: 776–789

Brewer GJ, Torricelli JR (2007) Isolation and culture of adult neurons and neurospheres. Nat Protoc 2: 1490–1498

Brilot AF, Korostelev AA, Ermolenko DN, Grigorieff N (2013) Structure of the ribosome with elongation factor G trapped in the pretranslocation state. Proceedings of the National Academy of Sciences of the United States of America 110: 20994–20999

Brito Querido J, Sokabe M, Kraatz S, Gordiyenko Y, Skehel JM, Fraser CS, Ramakrishnan V (2020) Structure of a human 48S translational initiation complex. Science 369: 1220–1227

Broadbelt K, Ramprasaud A, Jones LB (2006) Evidence of altered neurogranin immunoreactivity in areas 9 and 32 of schizophrenic prefrontal cortex. Schizophrenia research 87: 6–14

Buchan JR, Yoon JH, Parker R (2011) Stress-specific composition, assembly and kinetics of stress granules in Saccharomyces cerevisiae. J Cell Sci 124: 228–239

Budkevich T, Giesebrecht J, Altman RB, Munro JB, Mielke T, Nierhaus KH, Blanchard SC, Spahn CM (2011) Structure and dynamics of the mammalian ribosomal pretranslocation complex. Mol Cell 44: 214–224

Budkevich TV, Giesebrecht J, Behrmann E, Loerke J, Ramrath DJ, Mielke T, Ismer J, Hildebrand PW, Tung CS, Nierhaus KH et al (2014) Regulation of the mammalian elongation cycle by subunit rolling: a eukaryotic-specific ribosome rearrangement. Cell 158: 121–131

Burguete AS, Almeida S, Gao FB, Kalb R, Akins MR, Bonini NM (2015) GGGGCC microsatellite RNA is neuritically localized, induces branching defects, and perturbs transport granule function. eLife 4: e08881

Buxbaum AR, Wu B, Singer RH (2014) Single beta-actin mRNA detection in neurons reveals a mechanism for regulating its translatability. Science 343: 419–422

Cajigas IJ, Tushev G, Will TJ, tom Dieck S, Fuerst N, Schuman EM (2012) The local transcriptome in the synaptic neuropil revealed by deep sequencing and high-resolution imaging. Neuron 74: 453–466

Castle CD, Cassimere EK, Denicourt C (2012) LAS1L interacts with the mammalian Rix1 complex to regulate ribosome biogenesis. Mol Biol Cell 23: 716–728

Chambers MC, Maclean B, Burke R, Amodei D, Ruderman DL, Neumann S, Gatto L, Fischer B, Pratt B, Egertson J et al (2012) A cross-platform toolkit for mass spectrometry and proteomics. Nat Biotechnol 30: 918–920

Chen C, Cui X, Beausang JF, Zhang H, Farrell I, Cooperman BS, Goldman YE (2016) Elongation factor G initiates translocation through a power stroke. Proceedings of the National Academy of Sciences of the United States of America 113: 7515–7520

Chen PB, Kawaguchi R, Blum C, Achiro JM, Coppola G, O’Dell TJ, Martin KC (2017) Mapping Gene Expression in Excitatory Neurons during Hippocampal Late-Phase Long-Term Potentiation. Frontiers in molecular neuroscience 10: 39

Chen S, McMullan G, Faruqi AR, Murshudov GN, Short JM, Scheres SH, Henderson R (2013) High-resolution noise substitution to measure overfitting and validate resolution in 3D structure determination by single particle electron cryomicroscopy. Ultramicroscopy 135: 24–35

Chen VB, Arendall WB, 3rd, Headd JJ, Keedy DA, Immormino RM, Kapral GJ, Murray LW, Richardson JS, Richardson DC (2010) MolProbity: all-atom structure validation for macromolecular crystallography. Acta Crystallogr D Biol Crystallogr 66: 12–21

Chu JF, Majumder P, Chatterjee B, Huang SL, Shen CJ (2019) TDP-43 Regulates Coupled Dendritic mRNA Transport-Translation Processes in Co-operation with FMRP and Staufen1. Cell Rep 29: 3118–3133 e3116

Cioni JM, Lin JQ, Holtermann AV, Koppers M, Jakobs MAH, Azizi A, Turner-Bridger B, Shigeoka T, Franze K, Harris WA et al (2019) Late Endosomes Act as mRNA Translation Platforms and Sustain Mitochondria in Axons. Cell 176: 56–72 e15

Coffee RL, Jr., Williamson AJ, Adkins CM, Gray MC, Page TL, Broadie K (2012) In vivo neuronal function of the fragile X mental retardation protein is regulated by phosphorylation. Human molecular genetics 21: 900–915

Conicella AE, Zerze GH, Mittal J, Fawzi NL (2016) ALS Mutations Disrupt Phase Separation Mediated by alpha-Helical Structure in the TDP-43 Low-Complexity C-Terminal Domain. Structure 24: 1537–1549

Costa S, Almeida A, Castro A, Domingues L (2014) Fusion tags for protein solubility, purification and immunogenicity in Escherichia coli: the novel Fh8 system. Front Microbiol 5: 63

Costa-Mattioli M, Sossin WS, Klann E, Sonenberg N (2009) Translational control of long-lasting synaptic plasticity and memory. Neuron 61: 10–26

Cox J, Mann M (2008) MaxQuant enables high peptide identification rates, individualized p.p.b.-range mass accuracies and proteome-wide protein quantification. Nat Biotechnol 26: 1367–1372

Crick F (1984) Memory and molecular turnover. Nature 312: 101

Darnell JC, Van Driesche SJ, Zhang C, Hung KY, Mele A, Fraser CE, Stone EF, Chen C, Fak JJ, Chi SW et al (2011) FMRP stalls ribosomal translocation on mRNAs linked to synaptic function and autism. Cell 146: 247–261

de Araujo ME, Huber LA, Stasyk T (2008) Isolation of endocitic organelles by density gradient centrifugation. Methods Mol Biol 424: 317–331

DeJesus-Hernandez M, Mackenzie IR, Boeve BF, Boxer AL, Baker M, Rutherford NJ, Nicholson AM, Finch NA, Flynn H, Adamson J et al (2011) Expanded GGGGCC hexanucleotide repeat in noncoding region of C9ORF72 causes chromosome 9p-linked FTD and ALS. Neuron 72: 245–256

Dever TE, Green R (2012) The elongation, termination, and recycling phases of translation in eukaryotes. Cold Spring Harb Perspect Biol 4: a013706

Dictenberg JB, Swanger SA, Antar LN, Singer RH, Bassell GJ (2008) A direct role for FMRP in activity-dependent dendritic mRNA transport links filopodial-spine morphogenesis to fragile X syndrome. Developmental cell 14: 926–939

Dittman J, Ryan TA (2009) Molecular circuitry of endocytosis at nerve terminals. Annu Rev Cell Dev Biol 25: 133–160

Dobra I, Pankivskyi S, Samsonova A, Pastre D, Hamon L (2018) Relation Between Stress Granules and Cytoplasmic Protein Aggregates Linked to Neurodegenerative Diseases. Current neurology and neuroscience reports 18: 107

Dorner S, Brunelle JL, Sharma D, Green R (2006) The hybrid state of tRNA binding is an authentic translation elongation intermediate. Nat Struct Mol Biol 13: 234–241

Dynes JL, Steward O (2012) Arc mRNA docks precisely at the base of individual dendritic spines indicating the existence of a specialized microdomain for synapse-specific mRNA translation. J Comp Neurol 520: 3105–3119

El Fatimy R, Davidovic L, Tremblay S, Jaglin X, Dury A, Robert C, De Koninck P, Khandjian EW (2016) Tracking the Fragile X Mental Retardation Protein in a Highly Ordered Neuronal RiboNucleoParticles Population: A Link between Stalled Polyribosomes and RNA Granules. PLoS Genet 12: e1006192

Eliseev B, Yeramala L, Leitner A, Karuppasamy M, Raimondeau E, Huard K, Alkalaeva E, Aebersold R, Schaffitzel C (2018) Structure of a human cap-dependent 48S translation pre-initiation complex. Nucleic Acids Res 46: 2678–2689

Elvira G, Wasiak S, Blandford V, Tong XK, Serrano A, Fan X, del Rayo Sanchez-Carbente M, Servant F, Bell AW, Boismenu D et al (2006) Characterization of an RNA granule from developing brain. Molecular & cellular proteomics : MCP 5: 635–651

Emsley P, Lohkamp B, Scott WG, Cowtan K (2010) Features and development of Coot. Acta Crystallogr D Biol Crystallogr 66: 486–501

Ferbinteanu J (2019) Memory systems 2018 - Towards a new paradigm. Neurobiol Learn Mem 157: 61–78

Finkbeiner E, Haindl M, Raman N, Muller S (2011) SUMO routes ribosome maturation. Nucleus 2: 527–532

Flis J, Holm M, Rundlet EJ, Loerke J, Hilal T, Dabrowski M, Burger J, Mielke T, Blanchard SC, Spahn CMT et al (2018) tRNA Translocation by the Eukaryotic 80S Ribosome and the Impact of GTP Hydrolysis. Cell Rep 25: 2676–2688 e2677

Frank J, Agrawal RK (2000) A ratchet-like inter-subunit reorganization of the ribosome during translocation. Nature 406: 318–322

Ginsberg SD, Mufson EJ, Alldred MJ, Counts SE, Wuu J, Nixon RA, Che S (2011) Upregulation of select rab GTPases in cholinergic basal forebrain neurons in mild cognitive impairment and Alzheimer’s disease. J Chem Neuroanat 42: 102–110

Ginsberg SD, Mufson EJ, Counts SE, Wuu J, Alldred MJ, Nixon RA, Che S (2010) Regional selectivity of rab5 and rab7 protein upregulation in mild cognitive impairment and Alzheimer’s disease. J Alzheimers Dis 22: 631–639

Glock C, Biever A, Tushev G, Nassim-Assir B, Kao A, Bartnik I, Tom Dieck S, Schuman EM (2021) The translatome of neuronal cell bodies, dendrites, and axons. Proceedings of the National Academy of Sciences of the United States of America 118

Gonatopoulos-Pournatzis T, Niibori R, Salter EW, Weatheritt RJ, Tsang B, Farhangmehr S, Liang X, Braunschweig U, Roth J, Zhang S et al (2020) Autism-Misregulated eIF4G Microexons Control Synaptic Translation and Higher Order Cognitive Functions. Mol Cell 77: 1176–1192 e1116

Gopal PP, Nirschl JJ, Klinman E, Holzbaur EL (2017) Amyotrophic lateral sclerosis-linked mutations increase the viscosity of liquid-like TDP-43 RNP granules in neurons. Proceedings of the National Academy of Sciences of the United States of America 114: E2466–E2475

Graber TE, Freemantle E, Anadolu MN, Hebert-Seropian S, MacAdam RL, Shin U, Hoang HD, Alain T, Lacaille JC, Sossin WS (2017) UPF1 Governs Synaptic Plasticity through Association with a STAU2 RNA Granule. The Journal of neuroscience : the official journal of the Society for Neuroscience 37: 9116–9131

Graber TE, Hebert-Seropian S, Khoutorsky A, David A, Yewdell JW, Lacaille JC, Sossin WS (2013) Reactivation of stalled polyribosomes in synaptic plasticity. Proceedings of the National Academy of Sciences of the United States of America 110: 16205–16210

Hafner AS, Donlin-Asp PG, Leitch B, Herzog E, Schuman EM (2019) Local protein synthesis is a ubiquitous feature of neuronal pre- and postsynaptic compartments. Science 364

Hasan MT, Hernandez-Gonzalez S, Dogbevia G, Trevino M, Bertocchi I, Gruart A, Delgado-Garcia JM (2013) Role of motor cortex NMDA receptors in learning-dependent synaptic plasticity of behaving mice. Nat Commun 4: 2258

Heisler FF, Loebrich S, Pechmann Y, Maier N, Zivkovic AR, Tokito M, Hausrat TJ, Schweizer M, Bahring R, Holzbaur EL et al (2011) Muskelin regulates actin filament- and microtubule-based GABA(A) receptor transport in neurons. Neuron 70: 66–81

Hirokawa N (2006) mRNA transport in dendrites: RNA granules, motors, and tracks. The Journal of neuroscience : the official journal of the Society for Neuroscience 26: 7139–7142

Huerta PT, Sun LD, Wilson MA, Tonegawa S (2000) Formation of temporal memory requires NMDA receptors within CA1 pyramidal neurons. Neuron 25: 473–480

Huttlin EL, Bruckner RJ, Navarrete-Perea J, Cannon JR, Baltier K, Gebreab F, Gygi MP, Thornock A, Zarraga G, Tam S et al (2020) Dual Proteome-scale Networks Reveal Cell-specific Remodeling of the Human Interactome. bioRxiv: 2020.2001.2019.905109

Itofusa R, Kamiguchi H (2011) Polarizing membrane dynamics and adhesion for growth cone navigation. Mol Cell Neurosci 48: 332–338

Jalil SJ, Sacktor TC, Shouval HZ (2015) Atypical PKCs in memory maintenance: the roles of feedback and redundancy. Learning & memory 22: 344–353

Jiang YH, Yuen RK, Jin X, Wang M, Chen N, Wu X, Ju J, Mei J, Shi Y, He M et al (2013) Detection of clinically relevant genetic variants in autism spectrum disorder by whole-genome sequencing. Am J Hum Genet 93: 249–263

Jobert L, Argentini M, Tora L (2009) PRMT1 mediated methylation of TAF15 is required for its positive gene regulatory function. Experimental cell research 315: 1273–1286

Junutula JR, De Maziere AM, Peden AA, Ervin KE, Advani RJ, van Dijk SM, Klumperman J, Scheller RH (2004) Rab14 is involved in membrane trafficking between the Golgi complex and endosomes. Mol Biol Cell 15: 2218–2229

Kanai Y, Dohmae N, Hirokawa N (2004) Kinesin transports RNA: isolation and characterization of an RNA-transporting granule. Neuron 43: 513–525

Kedersha N, Cho MR, Li W, Yacono PW, Chen S, Gilks N, Golan DE, Anderson P (2000) Dynamic shuttling of TIA-1 accompanies the recruitment of mRNA to mammalian stress granules. J Cell Biol 151: 1257–1268

Kedersha N, Ivanov P, Anderson P (2013) Stress granules and cell signaling: more than just a passing phase? Trends Biochem Sci 38: 494–506

Kedersha N, Panas MD, Achorn CA, Lyons S, Tisdale S, Hickman T, Thomas M, Lieberman J, McInerney GM, Ivanov P et al (2016) G3BP-Caprin1-USP10 complexes mediate stress granule condensation and associate with 40S subunits. J Cell Biol 212: 845–860

Kennedy MJ, Ehlers MD (2006) Organelles and trafficking machinery for postsynaptic plasticity. Annu Rev Neurosci 29: 325–362

Kiebler MA, Hemraj I, Verkade P, Kohrmann M, Fortes P, Marion RM, Ortin J, Dotti CG (1999) The mammalian staufen protein localizes to the somatodendritic domain of cultured hippocampal neurons: implications for its involvement in mRNA transport. The Journal of neuroscience : the official journal of the Society for Neuroscience 19: 288–297

Knowles RB, Sabry JH, Martone ME, Deerinck TJ, Ellisman MH, Bassell GJ, Kosik KS (1996) Translocation of RNA granules in living neurons. The Journal of neuroscience : the official journal of the Society for Neuroscience 16: 7812–7820

Koenig E, Martin R, Titmus M, Sotelo-Silveira JR (2000) Cryptic peripheral ribosomal domains distributed intermittently along mammalian myelinated axons. The Journal of neuroscience : the official journal of the Society for Neuroscience 20: 8390–8400

Kolb B, Gibb R (2014) Searching for the principles of brain plasticity and behavior. Cortex 58: 251–260

Koppers M, Cagnetta R, Shigeoka T, Wunderlich LC, Vallejo-Ramirez P, Qiaojin Lin J, Zhao S, Jakobs MA, Dwivedy A, Minett MS et al (2019) Receptor-specific interactome as a hub for rapid cue-induced selective translation in axons. eLife 8

Kovalevich J, Langford D (2013) Considerations for the use of SH-SY5Y neuroblastoma cells in neurobiology. Methods Mol Biol 1078: 9–21

Krichevsky AM, Kosik KS (2001) Neuronal RNA granules: a link between RNA localization and stimulation-dependent translation. Neuron 32: 683–696

LaMonte BH, Wallace KE, Holloway BA, Shelly SS, Ascano J, Tokito M, Van Winkle T, Howland DS, Holzbaur EL (2002) Disruption of dynein/dynactin inhibits axonal transport in motor neurons causing late-onset progressive degeneration. Neuron 34: 715–727

Langdon EM, Gladfelter AS (2018) A New Lens for RNA Localization: Liquid-Liquid Phase Separation. Annu Rev Microbiol 72: 255–271

Langille JJ, Ginzberg K, Sossin WS (2019) Polysomes identified by live imaging of nascent peptides are stalled in hippocampal and cortical neurites. Learning & memory 26: 351–362

Leal G, Comprido D, Duarte CB (2014) BDNF-induced local protein synthesis and synaptic plasticity. Neuropharmacology 76 Pt C: 639–656

Lebeau G, Miller LC, Tartas M, McAdam R, Laplante I, Badeaux F, DesGroseillers L, Sossin WS, Lacaille JC (2011) Staufen 2 regulates mGluR long-term depression and Map1b mRNA distribution in hippocampal neurons. Learning & memory 18: 314–326

Lee AS, Kranzusch PJ, Doudna JA, Cate JH (2016) eIF3d is an mRNA cap-binding protein that is required for specialized translation initiation. Nature 536: 96–99

Leitner A, Faini M, Stengel F, Aebersold R (2016) Crosslinking and Mass Spectrometry: An Integrated Technology to Understand the Structure and Function of Molecular Machines. Trends Biochem Sci 41: 20–32

Leitner A, Reischl R, Walzthoeni T, Herzog F, Bohn S, Forster F, Aebersold R (2012) Expanding the chemical cross-linking toolbox by the use of multiple proteases and enrichment by size exclusion chromatography. Molecular & cellular proteomics : MCP 11: M111 014126

Li W, Ward FR, McClure KF, Chang ST, Montabana E, Liras S, Dullea RG, Cate JHD (2019) Structural basis for selective stalling of human ribosome nascent chain complexes by a drug-like molecule. Nat Struct Mol Biol 26: 501–509

Li YC, Cheng CX, Li YN, Shimada O, Atsumi S (2005) Beyond the initial axon segment of the spinal motor axon: fasciculated microtubules and polyribosomal clusters. J Anat 206: 535–542

Liao YC, Fernandopulle MS, Wang G, Choi H, Hao L, Drerup CM, Patel R, Qamar S, Nixon-Abell J, Shen Y et al (2019) RNA Granules Hitchhike on Lysosomes for Long-Distance Transport, Using Annexin A11 as a Molecular Tether. Cell 179: 147–164 e120

Liebschner D, Afonine PV, Baker ML, Bunkoczi G, Chen VB, Croll TI, Hintze B, Hung LW, Jain S, McCoy AJ et al (2019) Macromolecular structure determination using X-rays, neutrons and electrons: recent developments in Phenix. Acta Crystallogr D Struct Biol 75: 861–877

Lin Z, Gasic I, Chandrasekaran V, Peters N, Shao S, Mitchison TJ, Hegde RS (2020) TTC5 mediates autoregulation of tubulin via mRNA degradation. Science 367: 100–104

Lisman JE (1985) A mechanism for memory storage insensitive to molecular turnover: a bistable autophosphorylating kinase. Proceedings of the National Academy of Sciences of the United States of America 82: 3055–3057

Liu-Yesucevitz L, Bassell GJ, Gitler AD, Hart AC, Klann E, Richter JD, Warren ST, Wolozin B (2011) Local RNA translation at the synapse and in disease. The Journal of neuroscience : the official journal of the Society for Neuroscience 31: 16086–16093

Mallardo M, Deitinghoff A, Muller J, Goetze B, Macchi P, Peters C, Kiebler MA (2003) Isolation and characterization of Staufen-containing ribonucleoprotein particles from rat brain. Proceedings of the National Academy of Sciences of the United States of America 100: 2100–2105

Mateu-Regue A, Christiansen J, Bagger FO, Winther O, Hellriegel C, Nielsen FC (2019) Single mRNP Analysis Reveals that Small Cytoplasmic mRNP Granules Represent mRNA Singletons. Cell Rep 29: 736–748 e734

Matthes F, Hettich MM, Schilling J, Flores-Dominguez D, Blank N, Wiglenda T, Buntru A, Wolf H, Weber S, Vorberg I et al (2018) Inhibition of the MID1 protein complex: a novel approach targeting APP protein synthesis. Cell Death Discov 4: 4

Mikl M, Vendra G, Kiebler MA (2011) Independent localization of MAP2, CaMKIIalpha and beta-actin RNAs in low copy numbers. EMBO reports 12: 1077–1084

Miron T, Wilchek M (1982) A spectrophotometric assay for soluble and immobilized N-hydroxysuccinimide esters. Anal Biochem 126: 433–435

Miyakawa T, Yared E, Pak JH, Huang FL, Huang KP, Crawley JN (2001) Neurogranin null mutant mice display performance deficits on spatial learning tasks with anxiety related components. Hippocampus 11: 763–775

Munro JB, Altman RB, O’Connor N, Blanchard SC (2007) Identification of two distinct hybrid state intermediates on the ribosome. Mol Cell 25: 505–517

Nagaraj N, Wisniewski JR, Geiger T, Cox J, Kircher M, Kelso J, Paabo S, Mann M (2011) Deep proteome and transcriptome mapping of a human cancer cell line. Mol Syst Biol 7: 548

Nakane T, Kimanius D, Lindahl E, Scheres SH (2018) Characterisation of molecular motions in cryo-EM single-particle data by multi-body refinement in RELION. eLife 7

Nakane T, Scheres SHW (2021) Multi-body Refinement of Cryo-EM Images in RELION. Methods Mol Biol 2215: 145–160

Nakayama K, Ohashi R, Shinoda Y, Yamazaki M, Abe M, Fujikawa A, Shigenobu S, Futatsugi A, Noda M, Mikoshiba K et al (2017) RNG105/caprin1, an RNA granule protein for dendritic mRNA localization, is essential for long-term memory formation. eLife 6

Narayanan U, Nalavadi V, Nakamoto M, Pallas DC, Ceman S, Bassell GJ, Warren ST (2007) FMRP phosphorylation reveals an immediate-early signaling pathway triggered by group I mGluR and mediated by PP2A. The Journal of neuroscience : the official journal of the Society for Neuroscience 27: 14349–14357

Nastasijevic B, Becker NA, Wurster SE, Maher LJ, 3rd (2008) Sequence-specific binding of DNA and RNA to immobilized Nickel ions. Biochem Biophys Res Commun 366: 420–425

Noller HF, Lancaster L, Mohan S, Zhou J (2017) Ribosome structural dynamics in translocation: yet another functional role for ribosomal RNA. Q Rev Biophys 50: e12

Nunez J (2008) Primary Culture of Hippocampal Neurons from P0 Newborn Rats. J Vis Exp

O’Reilly FJ, Rappsilber J (2018) Cross-linking mass spectrometry: methods and applications in structural, molecular and systems biology. Nat Struct Mol Biol 25: 1000–1008

Oesterlin LK, Goody RS, Itzen A (2012) Posttranslational modifications of Rab proteins cause effective displacement of GDP dissociation inhibitor. Proceedings of the National Academy of Sciences of the United States of America 109: 5621–5626

Ohashi R, Shiina N (2020) Cataloguing and Selection of mRNAs Localized to Dendrites in Neurons and Regulated by RNA-Binding Proteins in RNA Granules. Biomolecules 10

Ohashi R, Takao K, Miyakawa T, Shiina N (2016) Comprehensive behavioral analysis of RNG105 (Caprin1) heterozygous mice: Reduced social interaction and attenuated response to novelty. Scientific reports 6: 20775

Okada Y, Yamazaki H, Sekine-Aizawa Y, Hirokawa N (1995) The neuron-specific kinesin superfamily protein KIF1A is a unique monomeric motor for anterograde axonal transport of synaptic vesicle precursors. Cell 81: 769–780

Olink-Coux M, Hollenbeck PJ (1996) Localization and active transport of mRNA in axons of sympathetic neurons in culture. The Journal of neuroscience : the official journal of the Society for Neuroscience 16: 1346–1358

Ostroff LE, Santini E, Sears R, Deane Z, Kanadia RN, LeDoux JE, Lhakhang T, Tsirigos A, Heguy A, Klann E (2019) Axon TRAP reveals learning-associated alterations in cortical axonal mRNAs in the lateral amgydala. eLife 8

Park HY, Lim H, Yoon YJ, Follenzi A, Nwokafor C, Lopez-Jones M, Meng X, Singer RH (2014) Visualization of dynamics of single endogenous mRNA labeled in live mouse. Science 343: 422–424

Pazo A, Perez-Gonzalez A, Oliveros JC, Huarte M, Chavez JP, Nieto A (2019) hCLE/RTRAF-HSPC117-DDX1-FAM98B: A New Cap-Binding Complex That Activates mRNA Translation. Front Physiol 10: 92

Pettersen EF, Goddard TD, Huang CC, Couch GS, Greenblatt DM, Meng EC, Ferrin TE (2004) UCSF Chimera--a visualization system for exploratory research and analysis. J Comput Chem 25: 1605–1612

Pintilie GD, Zhang J, Goddard TD, Chiu W, Gossard DC (2010) Quantitative analysis of cryo-EM density map segmentation by watershed and scale-space filtering, and fitting of structures by alignment to regions. J Struct Biol 170: 427–438

Pushpalatha KV, Besse F (2019) Local Translation in Axons: When Membraneless RNP Granules Meet Membrane-Bound Organelles. Front Mol Biosci 6: 129

Quiocho FA, Spurlino JC, Rodseth LE (1997) Extensive features of tight oligosaccharide binding revealed in high-resolution structures of the maltodextrin transport/chemosensory receptor. Structure 5: 997–1015

Ratje AH, Loerke J, Mikolajka A, Brunner M, Hildebrand PW, Starosta AL, Donhofer A, Connell SR, Fucini P, Mielke T et al (2010) Head swivel on the ribosome facilitates translocation by means of intra-subunit tRNA hybrid sites. Nature 468: 713–716

Reineke LC, Lloyd RE (2013) Diversion of stress granules and P-bodies during viral infection. Virology 436: 255–267

Rodriguez L, Mohamed NV, Desjardins A, Lippe R, Fon EA, Leclerc N (2017) Rab7A regulates tau secretion. J Neurochem 141: 592–605

Rohou A, Grigorieff N (2015) CTFFIND4: Fast and accurate defocus estimation from electron micrographs. J Struct Biol 192: 216–221

Routtenberg A, Rekart JL (2005) Post-translational protein modification as the substrate for long-lasting memory. Trends Neurosci 28: 12–19

Rozes-Salvador V, Gonzalez-Billault C, Conde C (2020) The Recycling Endosome in Nerve Cell Development: One Rab to Rule Them All? Front Cell Dev Biol 8: 603794

Russo A, Scardigli R, La Regina F, Murray ME, Romano N, Dickson DW, Wolozin B, Cattaneo A, Ceci M (2017) Increased cytoplasmic TDP-43 reduces global protein synthesis by interacting with RACK1 on polyribosomes. Human molecular genetics 26: 1407–1418

Sahoo PK, Lee SJ, Jaiswal PB, Alber S, Kar AN, Miller-Randolph S, Taylor EE, Smith T, Singh B, Ho TS et al (2018) Axonal G3BP1 stress granule protein limits axonal mRNA translation and nerve regeneration. Nat Commun 9: 3358

Salapa HE, Hutchinson C, Popescu BF, Levin MC (2020) Neuronal RNA-binding protein dysfunction in multiple sclerosis cortex. Ann Clin Transl Neurol 7: 1214–1224

Santos AR, Comprido D, Duarte CB (2010) Regulation of local translation at the synapse by BDNF. Prog Neurobiol 92: 505–516

Scheres SH (2012) A Bayesian view on cryo-EM structure determination. J Mol Biol 415: 406–418

Schroter CJ, Braun M, Englert J, Beck H, Schmid H, Kalbacher H (1999) A rapid method to separate endosomes from lysosomal contents using differential centrifugation and hypotonic lysis of lysosomes. J Immunol Methods 227: 161–168

Schwanhausser B, Busse D, Li N, Dittmar G, Schuchhardt J, Wolf J, Chen W, Selbach M (2011) Global quantification of mammalian gene expression control. Nature 473: 337–342

Semenkov YP, Rodnina MV, Wintermeyer W (2000) Energetic contribution of tRNA hybrid state formation to translocation catalysis on the ribosome. Nat Struct Biol 7: 1027–1031

Shepherd JD, Rumbaugh G, Wu J, Chowdhury S, Plath N, Kuhl D, Huganir RL, Worley PF (2006) Arc/Arg3.1 mediates homeostatic synaptic scaling of AMPA receptors. Neuron 52: 475–484

Shigeoka T, Jung H, Jung J, Turner-Bridger B, Ohk J, Lin JQ, Amieux PS, Holt CE (2016) Dynamic Axonal Translation in Developing and Mature Visual Circuits. Cell 166: 181–192

Shiina N (2019) Liquid- and solid-like RNA granules form through specific scaffold proteins and combine into biphasic granules. J Biol Chem 294: 3532–3548

Shiina N, Shinkura K, Tokunaga M (2005) A novel RNA-binding protein in neuronal RNA granules: regulatory machinery for local translation. The Journal of neuroscience : the official journal of the Society for Neuroscience 25: 4420–4434

Shiina N, Tokunaga M (2010) RNA granule protein 140 (RNG140), a paralog of RNG105 localized to distinct RNA granules in neuronal dendrites in the adult vertebrate brain. J Biol Chem 285: 24260–24269

Shiina N, Yamaguchi K, Tokunaga M (2010) RNG105 deficiency impairs the dendritic localization of mRNAs for Na+/K+ ATPase subunit isoforms and leads to the degeneration of neuronal networks. The Journal of neuroscience : the official journal of the Society for Neuroscience 30: 12816–12830

Slavov N, Semrau S, Airoldi E, Budnik B, van Oudenaarden A (2015) Differential Stoichiometry among Core Ribosomal Proteins. Cell Rep 13: 865–873

Solomon S, Xu Y, Wang B, David MD, Schubert P, Kennedy D, Schrader JW (2007) Distinct structural features of caprin-1 mediate its interaction with G3BP-1 and its induction of phosphorylation of eukaryotic translation initiation factor 2alpha, entry to cytoplasmic stress granules, and selective interaction with a subset of mRNAs. Mol Cell Biol 27: 2324–2342

Sotelo JR, Kun A, Benech JC, Giuditta A, Morillas J, Benech CR (1999) Ribosomes and polyribosomes are present in the squid giant axon: an immunocytochemical study. Neuroscience 90: 705–715

Spaulding EL, Burgess RW (2017) Accumulating Evidence for Axonal Translation in Neuronal Homeostasis. Front Neurosci 11: 312

Spencer GE, Syed NI, van Kesteren E, Lukowiak K, Geraerts WP, van Minnen J (2000) Synthesis and functional integration of a neurotransmitter receptor in isolated invertebrate axons. J Neurobiol 44: 72–81

Stenmark H (2009) Rab GTPases as coordinators of vesicle traffic. Nat Rev Mol Cell Biol 10: 513–525

Steward O, Falk PM (1986) Protein-synthetic machinery at postsynaptic sites during synaptogenesis: a quantitative study of the association between polyribosomes and developing synapses. The Journal of neuroscience : the official journal of the Society for Neuroscience 6: 412–423

Steward O, Levy WB (1982) Preferential localization of polyribosomes under the base of dendritic spines in granule cells of the dentate gyrus. The Journal of neuroscience : the official journal of the Society for Neuroscience 2: 284–291

Steward O, Ribak CE (1986) Polyribosomes associated with synaptic specializations on axon initial segments: localization of protein-synthetic machinery at inhibitory synapses. The Journal of neuroscience : the official journal of the Society for Neuroscience 6: 3079–3085

Stiege W, Stade K, Schuler D, Brimacombe R (1988) Covalent cross-linking of poly(A) to Escherichia coli ribosomes, and localization of the cross-link site within the 16S RNA. Nucleic Acids Res 16: 2369–2388

Sudhakaran IP, Hillebrand J, Dervan A, Das S, Holohan EE, Hulsmeier J, Sarov M, Parker R, VijayRaghavan K, Ramaswami M (2014) FMRP and Ataxin-2 function together in long-term olfactory habituation and neuronal translational control. Proceedings of the National Academy of Sciences of the United States of America 111: E99–E108

Sullivan MA, Vilaplana F, Cave RA, Stapleton D, Gray-Weale AA, Gilbert RG (2010) Nature of alpha and beta particles in glycogen using molecular size distributions. Biomacromolecules 11: 1094–1100

Sutton MA, Schuman EM (2006) Dendritic protein synthesis, synaptic plasticity, and memory. Cell 127: 49–58

Tan D, Li Q, Zhang MJ, Liu C, Ma C, Zhang P, Ding YH, Fan SB, Tao L, Yang B et al (2016) Trifunctional cross-linker for mapping protein-protein interaction networks and comparing protein conformational states. eLife 5

Tang WC, Lin RJ, Liao CL, Lin YL (2014) Rab18 facilitates dengue virus infection by targeting fatty acid synthase to sites of viral replication. J Virol 88: 6793–6804

Tourriere H, Chebli K, Zekri L, Courselaud B, Blanchard JM, Bertrand E, Tazi J (2003) The RasGAP-associated endoribonuclease G3BP assembles stress granules. J Cell Biol 160: 823–831

Tsien JZ, Huerta PT, Tonegawa S (1996) The essential role of hippocampal CA1 NMDA receptor-dependent synaptic plasticity in spatial memory. Cell 87: 1327–1338

Tyson JJ, Chen KC, Novak B (2003) Sniffers, buzzers, toggles and blinkers: dynamics of regulatory and signaling pathways in the cell. Curr Opin Cell Biol 15: 221–231

Urbanska AS, Janusz-Kaminska A, Switon K, Hawthorne AL, Perycz M, Urbanska M, Bassell GJ, Jaworski J (2017) ZBP1 phosphorylation at serine 181 regulates its dendritic transport and the development of dendritic trees of hippocampal neurons. Scientific reports 7: 1876

Urnavicius L, Zhang K, Diamant AG, Motz C, Schlager MA, Yu M, Patel NA, Robinson CV, Carter AP (2015) The structure of the dynactin complex and its interaction with dynein. Science 347: 1441–1446

van de Waterbeemd M, Tamara S, Fort KL, Damoc E, Franc V, Bieri P, Itten M, Makarov A, Ban N, Heck AJR (2018) Dissecting ribosomal particles throughout the kingdoms of life using advanced hybrid mass spectrometry methods. Nat Commun 9: 2493

Verdugo A, Vinod PK, Tyson JJ, Novak B (2013) Molecular mechanisms creating bistable switches at cell cycle transitions. Open Biol 3: 120179

Walker IH, Hsieh PC, Riggs PD (2010) Mutations in maltose-binding protein that alter affinity and solubility properties. Appl Microbiol Biotechnol 88: 187–197

Wareski P, Vaarmann A, Choubey V, Safiulina D, Liiv J, Kuum M, Kaasik A (2009) PGC-1{alpha} and PGC-1{beta} regulate mitochondrial density in neurons. J Biol Chem 284: 21379–21385

Weng FJ, Garcia RI, Lutzu S, Alvina K, Zhang Y, Dushko M, Ku T, Zemoura K, Rich D, Garcia-Dominguez D et al (2018) Npas4 Is a Critical Regulator of Learning-Induced Plasticity at Mossy Fiber-CA3 Synapses during Contextual Memory Formation. Neuron 97: 1137–1152 e1135

White JP, Cardenas AM, Marissen WE, Lloyd RE (2007) Inhibition of cytoplasmic mRNA stress granule formation by a viral proteinase. Cell Host Microbe 2: 295–305

Winckler B, Mellman I (2010) Trafficking guidance receptors. Cold Spring Harb Perspect Biol 2: a001826

Yang B, Wu YJ, Zhu M, Fan SB, Lin J, Zhang K, Li S, Chi H, Li YX, Chen HF et al (2012) Identification of cross-linked peptides from complex samples. Nat Methods 9: 904–906

Zerial M, McBride H (2001) Rab proteins as membrane organizers. Nat Rev Mol Cell Biol 2: 107–117

Zivanov J, Nakane T, Forsberg BO, Kimanius D, Hagen WJ, Lindahl E, Scheres SH (2018) New tools for automated high-resolution cryo-EM structure determination in RELION-3. eLife 7

Zivanov J, Nakane T, Scheres SHW (2019) A Bayesian approach to beam-induced motion correction in cryo-EM single-particle analysis. IUCrJ 6: 5–17

Zois CE, Favaro E, Harris AL (2014) Glycogen metabolism in cancer. Biochem Pharmacol 92: 3–11

