## Supplementary figures and images for "Neuronal RNA granules are ribosome complexes stalled at the pre-translocation state"

### Supplementary Figure S0

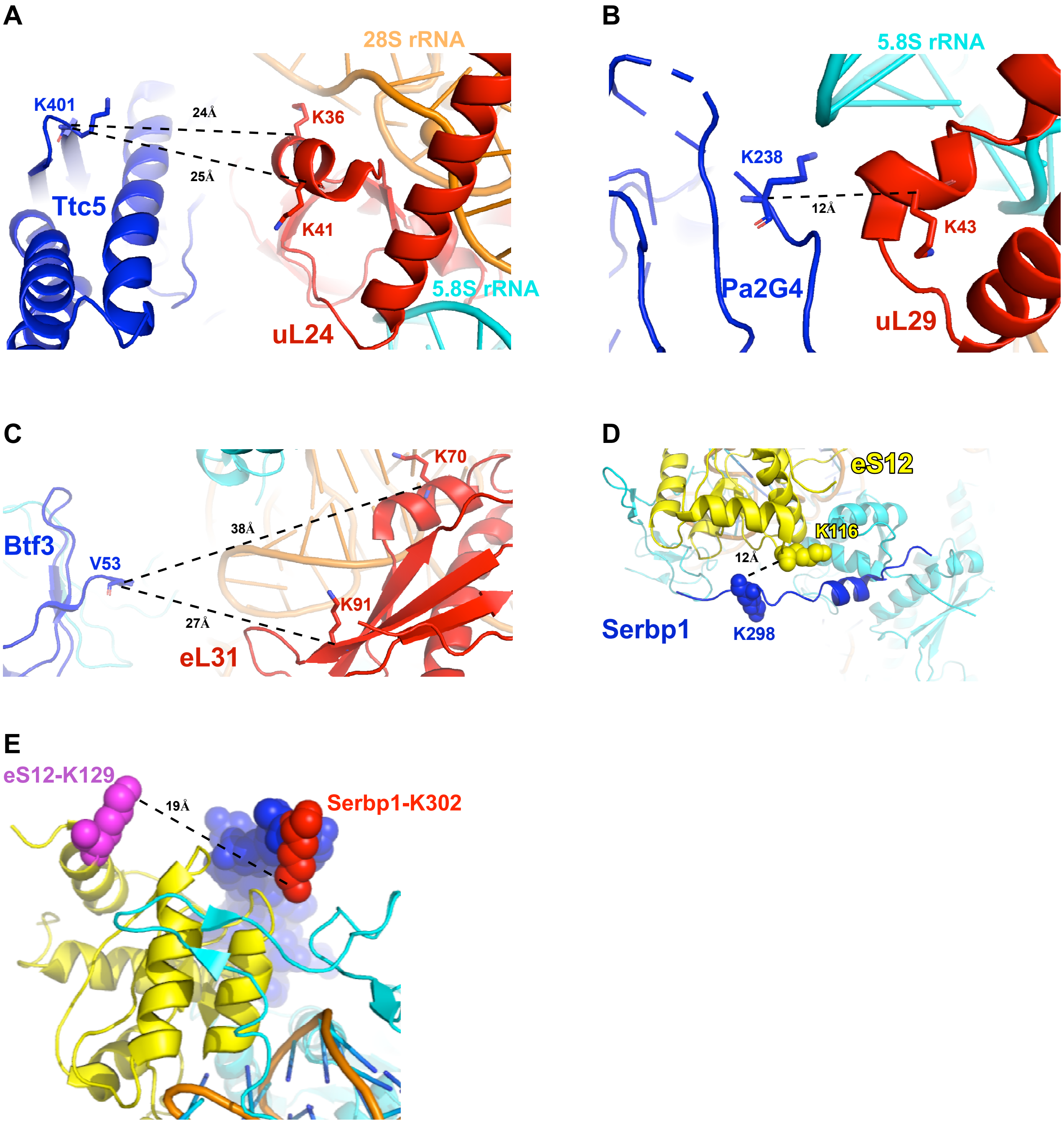


**Supplementary Figure S10**

### Supplementary Figure S1

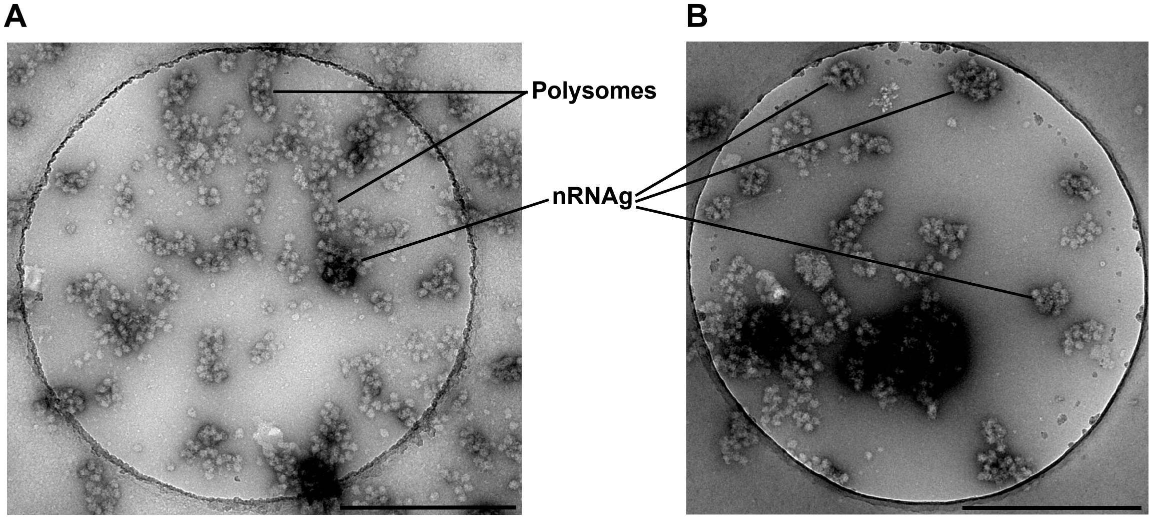


Supplementary Figure 1.

### Supplementary Figure S2

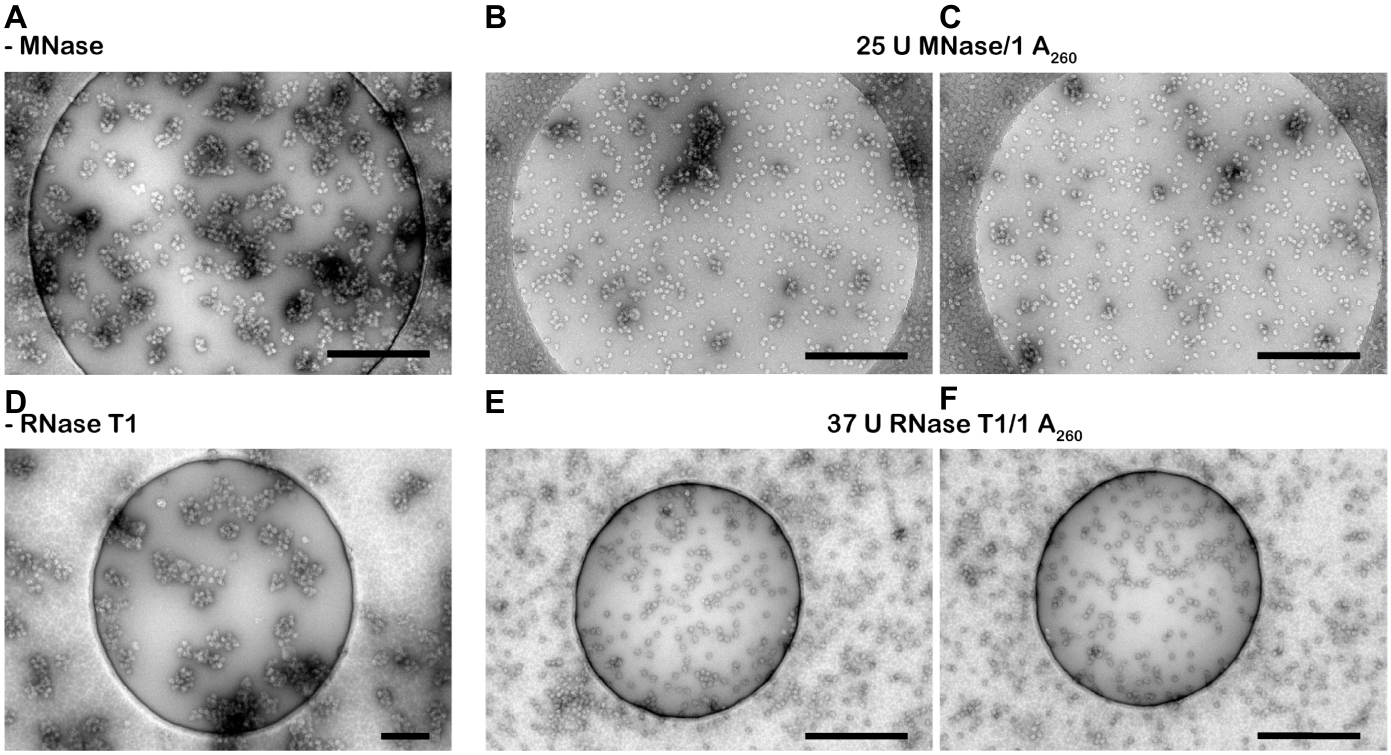


Supplementary Figure S2

### Supplementary Figure S3

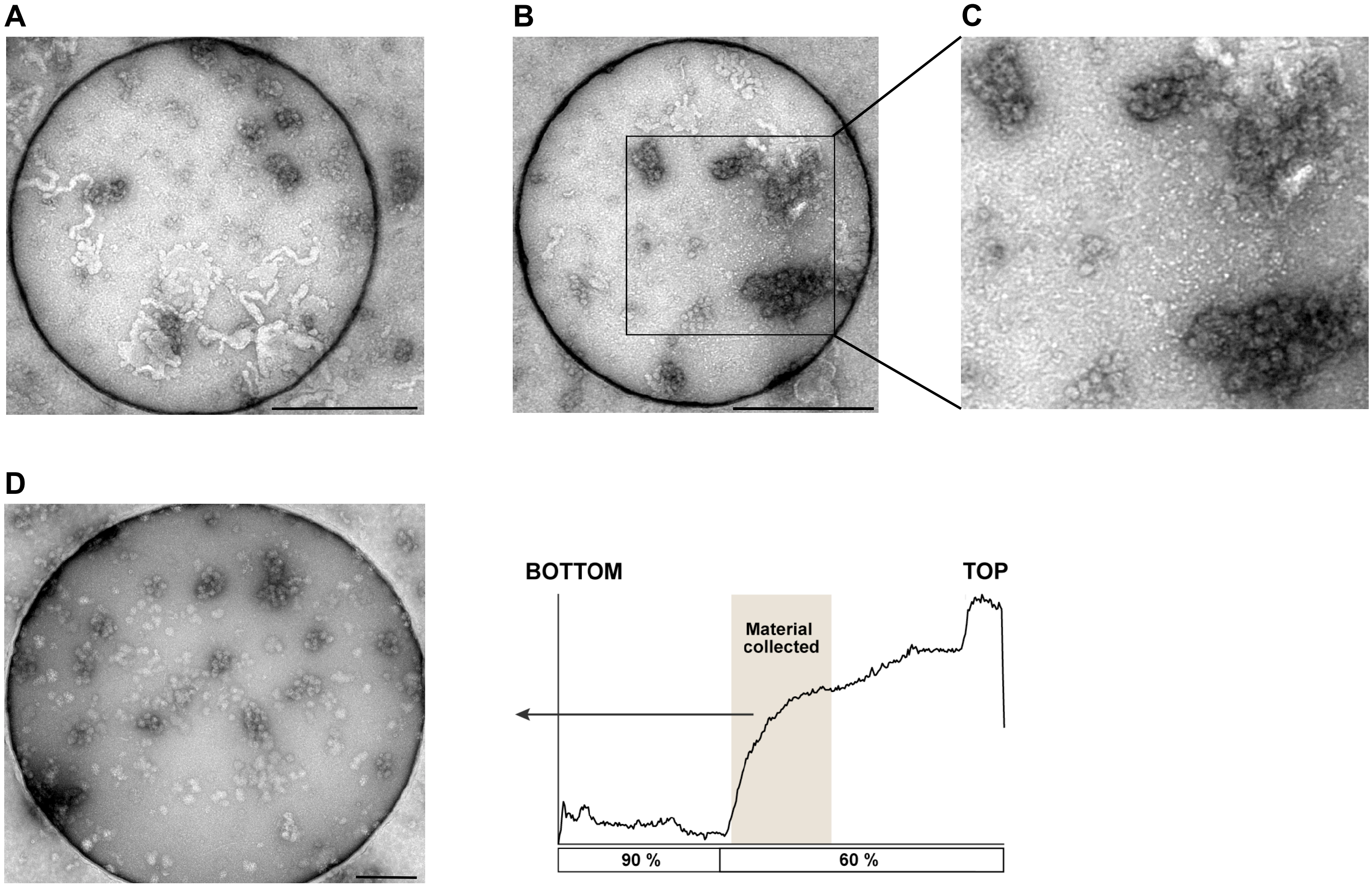


Supplementary Figure S3.

### Supplementary Figure S4

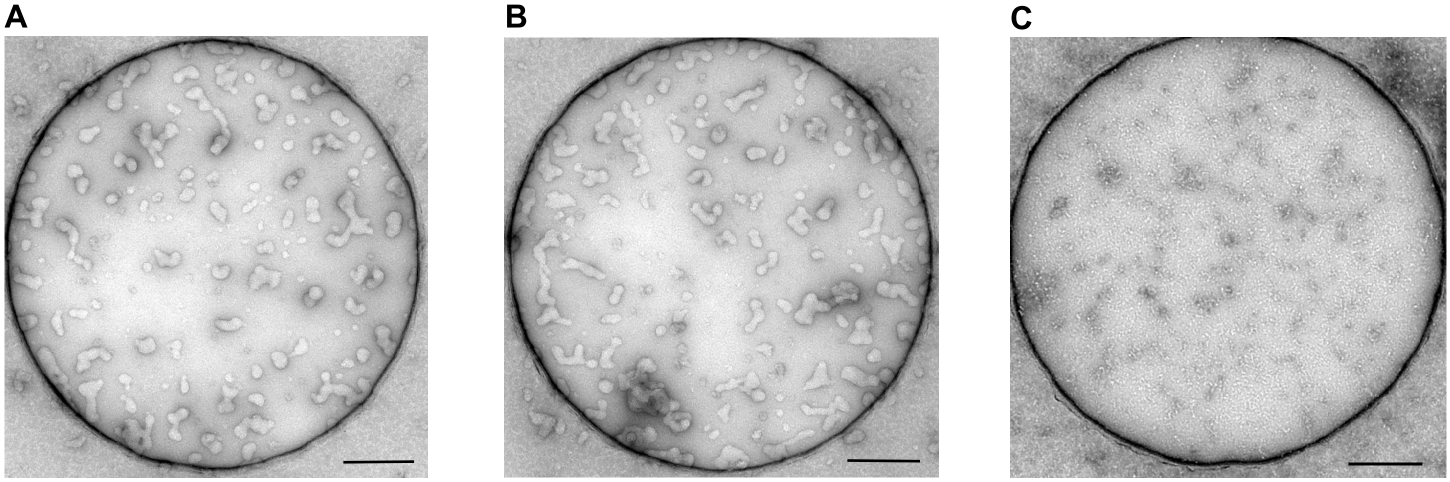

Supplementary Figure S4.

### Supplementary Figure S5 S6 S8

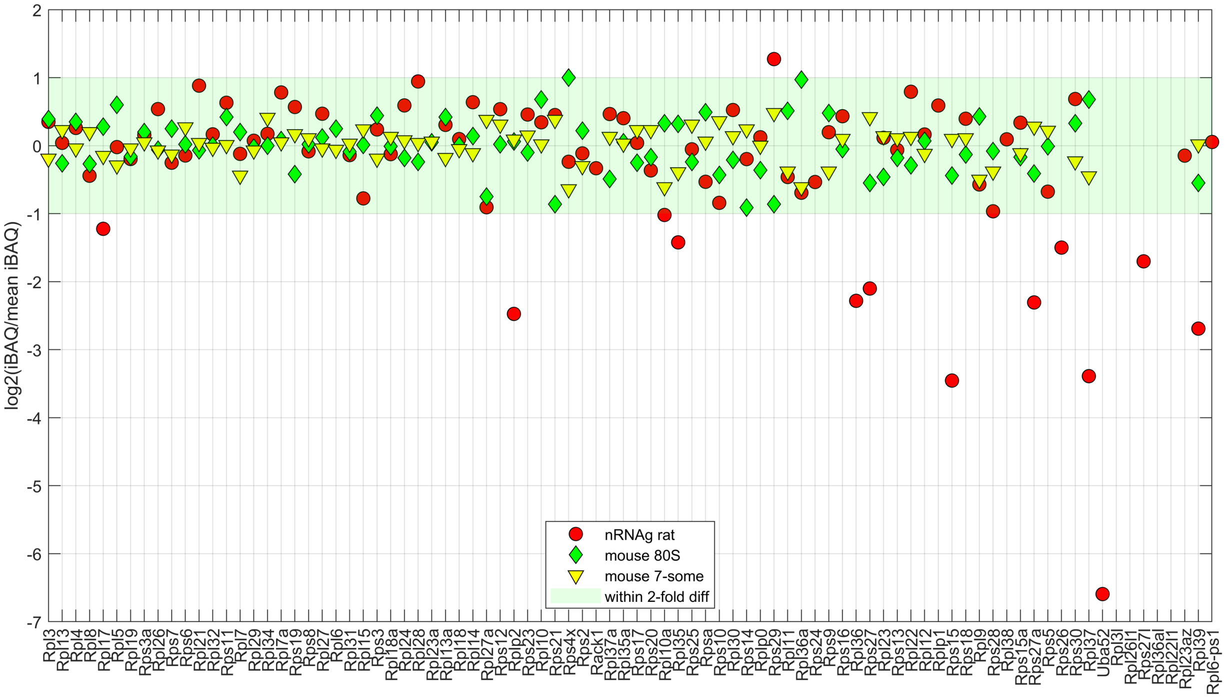


Supplementary Figure S5.

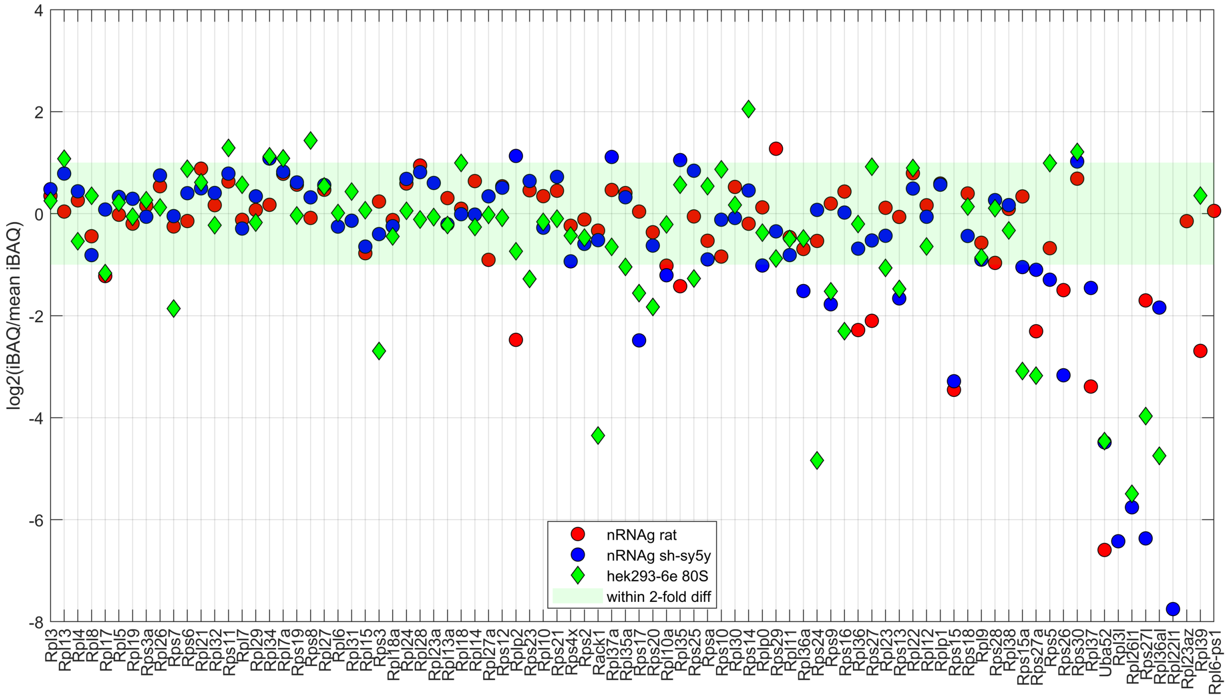


Supplementary Figure S6.

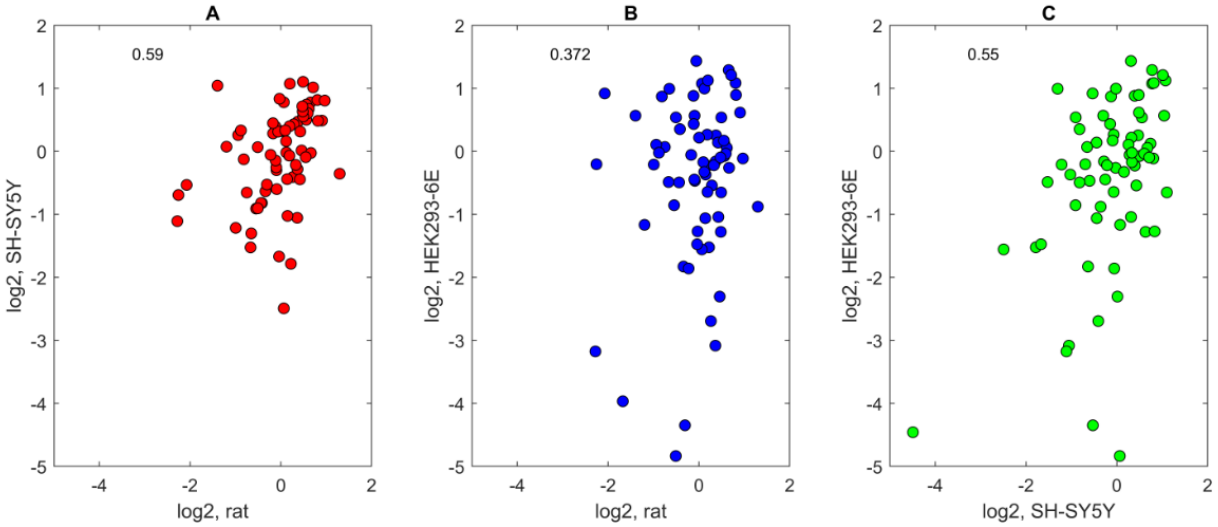


Supplementary Figure S8.

### Supplementary Figure S7

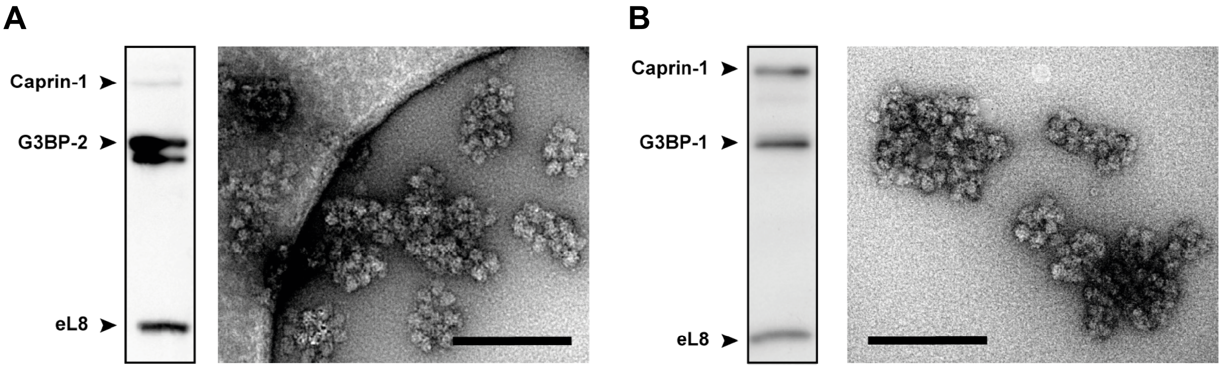


Supplementary Figure S7.

### Supplementary Figure S9

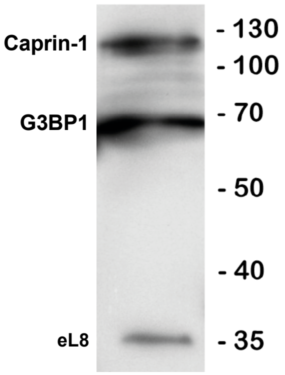


Supplementary Figure S9

### Supplementary Figure S11

A

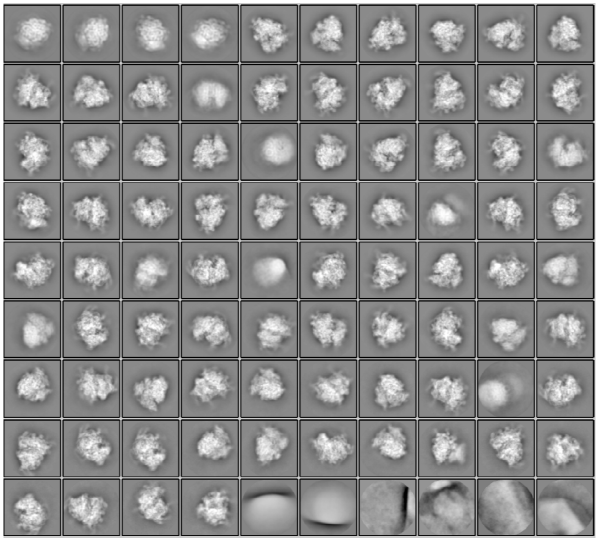

B

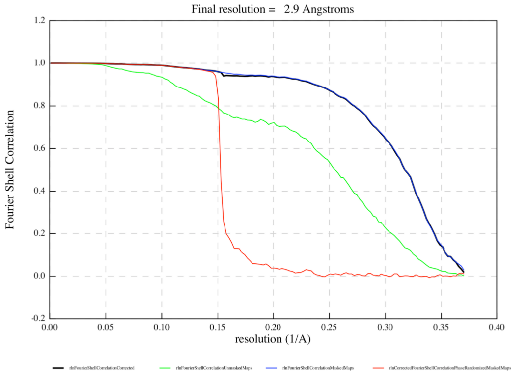


Supplementary Figure S11

### Supplementary Figure S12

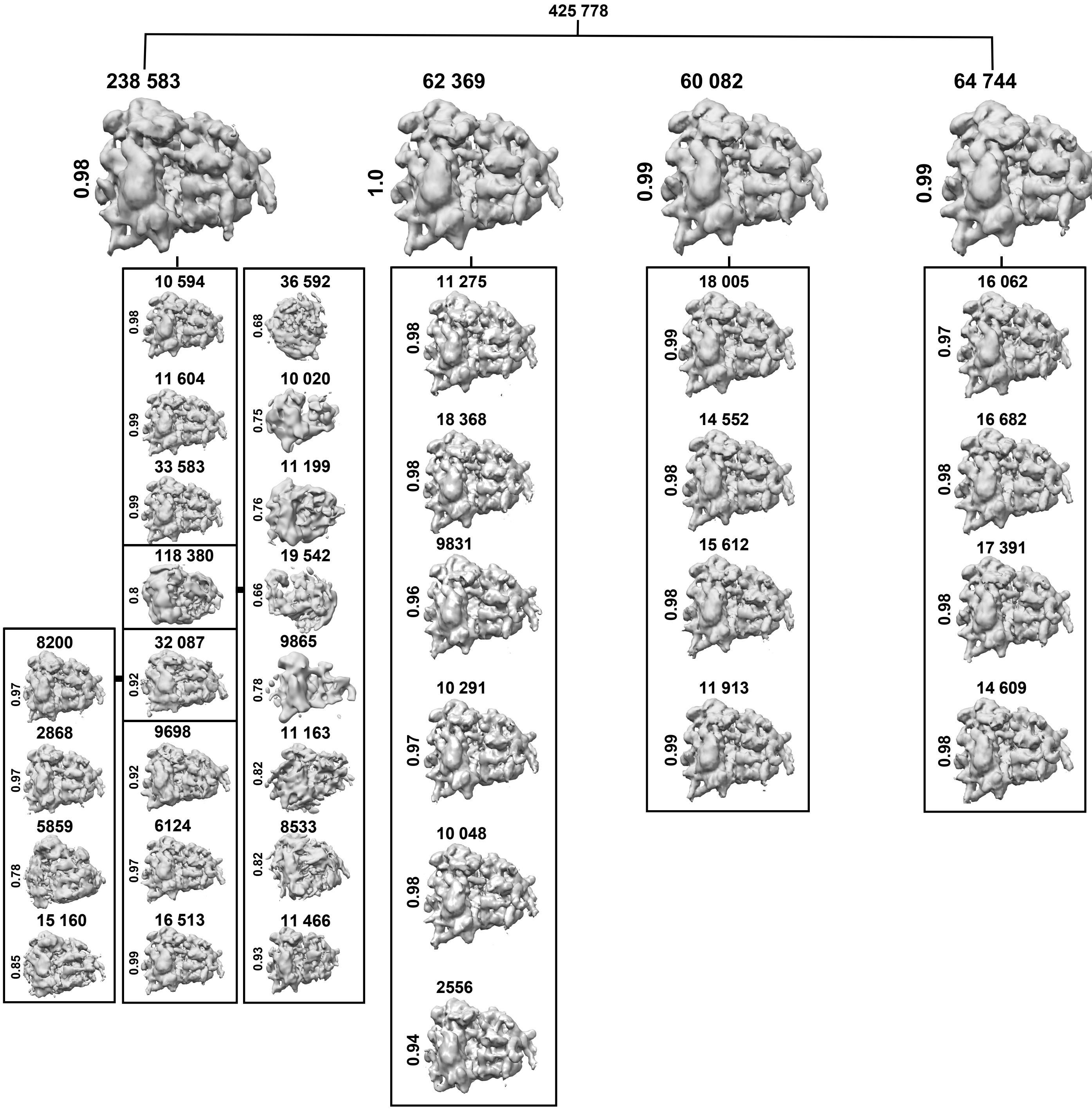


**Supplementary Figure S12**

### Supplementary Figure S15

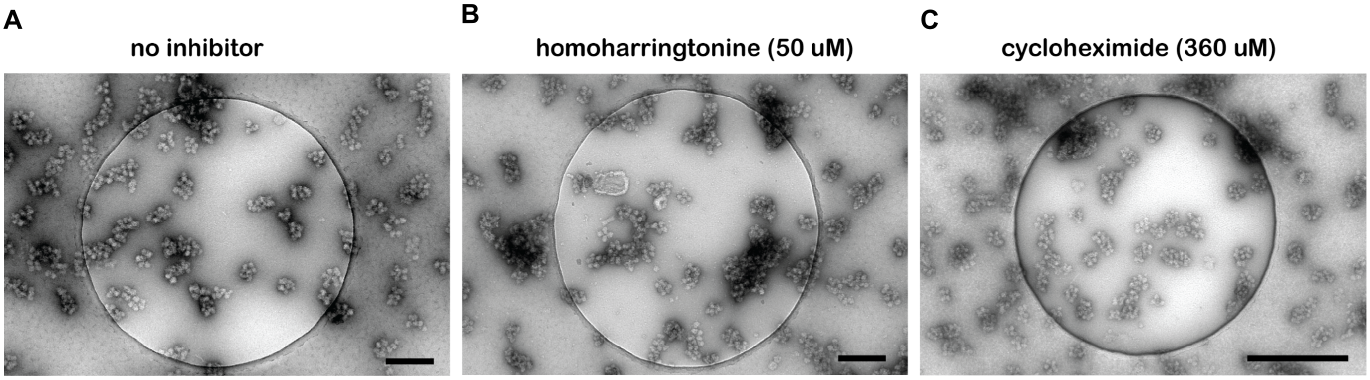


Supplementary Figure S15.
