## Supplementary Figure S13 for "Neuronal RNA granules are ribosome complexes stalled at the pre-translocation state"

**Consensus**

**1.** Rat 18S
**2.** Human 18S

```
1    1    TACCTGGTTGATCCTGCCAGTAGCATATGCTTGTCTCAAAGATTAAGCCATGCATGTCTA  60
2    1    ............................................................  60

1    61   AGTACGCACGGCCGGTACAGTGAAACTGCGAATGGCTCATTAAATCAGTTATGGTTCCTT  120
2    61   ............................................................  120

1    121  TGGTCGCTCGCTCCTCTCCTACTTGGATAACTGTGGTAATTCTAGAGCTAATACATGCCG  180
2    121  ............................................................  180

1    181  ACGGGCGCTGACCCCCCTTCCCGTGGGGGGACGCGTGCATTTATCAGATCAAAACCAAC  240
2    181  ................T.---..C........T...........................  237

1    241  CCGGTCAGCCCC-CTCCCGGCTCCGGCCGGGGGTCGGGCGCCGGCGGCTTTGGTGACTCT  299
2    238  ............T..-.....C...........G..........................  296

1    300  AGATAACCTCGGGCCGATCGCACGCCCTCCGTGGCGGCGACGACCCATTCGAACGTCTGC  359
2    297  ...........................C................................  356

1    360  CCTATCAACTTTCGATGGTAGTCGCCGTGCCTACCATGGTGACCACGGGTGACGGGGAAT  419
2    357  ............................................................  416

1    420  CAGGGTTCGATTCCGGAGAGGGAGCCTGAGAAACGGCTACCACATCCAAGGAAGGCAGCA  479
2    417  ............................................................  476

1    480  GGCGCGCAAATTACCCACTCCCGACCCGGGGAGGTAGTGACGAAAAATAACAATACAGGA  539
2    477  ............................................................  536

1    540  CTCTTTCGAGGCCCTGTAATTGGAATGAGTCCACTTTAAATCCTTTAACGAGGATCCATT  599
2    537  ............................................................  596

1    600  GGAGGGCAAGTCTGGTGCCAGCAGCCGCGGTAATTCCAGCTCCAATAGCGTATATTAAAG  659
2    597  ............................................................  656
```

```
1   660  TTGCTGCAGTTAAAAAGCTCGTAGTTGGATCTTGGGAGCGGGCGGGCGGTCCGCCGCGAG  719
2   657  ............................................................  716

1   720  GCGAGTCACCGCCCGTCCCCGCCCCTTGCCTCTCGGCGCCCCCTCGATGCTCTTAGCTGA  779
2   717  .....C......................................................  776

1   780  GTGTCCCGCGGGGCCCGAAGCGTTTACTTTGAAAAAATTAGAGTGTTCAAAGCAGGCCCG  839
2   777  ............................................................  836

1   840  AGCCGCCTGGATACCGCAGCTAGGAATAATGGAATAGGACCGCGGTTCTATTTTGTTGGT  899
2   837  ............................................................  896

1   900  TTTCGGAACTGAGGCCATGATTAAGAGGGACGGCCGGGGGCATTCGTATTGCGCCGCTAG  959
2   897  ............................................................  956

1   960  AGGTGAAATTCTTGGACCGGCGCAAGACGGACCAGAGCGAAAGCATTTGCCAAGAATGTT  1019
2   957  ............................................................  1016

1   1020 TTCATTAATCAAGAACGAAAGTCGGAGGTTCGAAGACGATCAGATACCGTCGTAGTTCCG  1079
2   1017 ............................................................  1076

1   1080 ACCATAAACGATGCCGACTGGCGATGCGGCGGCGTTATTCCCATGACCCGCCGGGCAGCT  1139
2   1077 ..................C.........................................  1136

1   1140 TCCGGGAAACCAAAGTCTTTGGGTTCCGGGGGGAGTATGGTTGCAAAGCTGAAACTTAAA  1199
2   1137 ............................................................  1196

1   1200 GGAATTGACGGAAGGGCACCACCAGGAGTGGAGCCTGCGGCTTAATTTGACTCAACACGG  1259
2   1197 ............................................................  1256

1   1260 GAAACCTCACCCGGCCCGGACACGGACAGGATTGACAGATTGATAGCTCTTTCTCGATTC  1319
2   1257 ............................................................  1316

1   1320 CGTGGGTGGTGGTGCATGGCCGTTCTTAGTTGGTGGAGCGATTTGTCTGGTTAATTCCGA  1379
2   1317 ............................................................  1376
```

```
1   1380 TAACGAACGAGACTCTGGCATGCTAACTAGTTACGCGACCCCCGAGCGGTCGGCGTCCCC 1439
2   1377 ............................................................ 1436

1   1440 CAACTTCTTAGAGGGACAAGTGGCGTTCAGCCACCCGAGATTGAGCAATAACAGGTCTGT 1499
2   1437 ............................................................ 1496

1   1500 GATGCCCTTAGATGTCCGGGGCTGCACGCGCGCTACACTGACTGGCTCAGCGTGTGCCTA 1559
2   1497 ............................................................ 1556

1   1560 CCCTACGCCGGCAGGCGCGGGTAACCCGTTGAACCCCATTCGTGATGGGGATCGGGGATT 1619
2   1557 ............................................................ 1616

1   1620 GCAATTATTCCCCATGAACGAGGAATTCCCAGTAAGTGCGGGTCATAAGCTTGCGTTGAT 1679
2   1617 ............................................................ 1676

1   1680 TAAGTCCCTGCCCTTTGTACACACCGCCCGTCGCTACTACCGATTGGATGGTTTAGTGAG 1739
2   1677 ............................................................ 1736

1   1740 GCCCTCGGATCGGCCCCGCCGGGGTCGGCCCACGGCCCTGGCGGAGCGCTGAGAAGACGG 1799
2   1737 ............................................................ 1796

1   1800 TCGAACTTGACTATCTAGAGGAAGTAAAAGTCGTAACAAGGTTTCCGTAGGTGAACCTGC 1859
2   1797 ............................................................ 1856

1   1860 GGAAGGATCATTA 1872
2   1857 ............. 1869
```
