## Supplementary Figure S14 for "Neuronal RNA granules are ribosome complexes stalled at the pre-translocation state"

**Consensus**

1. Human 28S
2. Rat 28S
3. Rat 28S seq

```
  CGCGACCTCAGATCAGACGTGGCGACCCGCTGAATTTAAGCATATTAGTCAGCGGAGGAA
1 CGCGACCTCAGATCAGACGTGGCGACCCGCTGAATTTAAGCATATTAGTCAGCGGAGGAA 60
2 CGCGACCTCAGATCAGACGTGGCGACCCGCTGAATTTAAGCATATTAGTCAGCGGAGGAA 60
3 CGCGACCTCAGATCAGACGTGGCGACCCGCTGAATTTAAGCATATTAGTCAGCGGAGGAA 60

  AAGAAACTAACCAGGATTCCCTCAGTAACGGCGAGTGAACAGGGAAGAGCCCAGCGCCGA
1 AAGAAACTAACCAGGATTCCCTCAGTAACGGCGAGTGAACAGGGAAGAGCCCAGCGCCGA 120
2 AAGAAACTAACCAGGATTCCCTCAGTAACGGCGAGTGAACAGGGAAGAGCCCAGCGCCGA 120
3 AAGAAACTAACCAGGATTCCCTCAGTAACGGCGAGTGAACAGGGAAGAGCCCAGCGCCGA 120

  ATCCCCGCCGCGCGCCGCGGCGCGGGAAATGTGGCGTACGGAAGACCCACTCCCCGGCGC
1 ATCCCCGCCCCGCGGCGGGGCGCGGGACATGTGGCGTACGGAAGACCCGCTCCCCGGCGC 180
2 ATCCCCGCCGCGCGCCGCGGCGCGGGAAATGTGGCGTACGGAAGACCCACTCCCCGGCGC 180
3 ATCCCCGCCGCGCGCCGCGGCGCGGGAAATGTGGCGTACGGAAGACCCACTCCCCGGCGC 180

  CGCTCGTGGGGGGCCCAAGTCCTTCTGATCGAGGCCCAGCCCGTGGACGGTGTGAGGCCG
1 CGCTCGTGGGGGGCCCAAGTCCTTCTGATCGAGGCCCAGCCCGTGGACGGTGTGAGGCCG 240
2 CGCTCGTGGGGGGCCCAAGTCCTTCTGATCGAGGCCCAGCCCGTGGACGGTGTGAGGCCG 240
3 CGCTCGTGGGGGGCCCAAGTCCTTCTGATCGAGGCCCAGCCCGTGGACGGTGTGAGGCCG 240

  GTAGCGGCCCCCGGCGCGCCGGGCCCGGGTCTTCCCGGAGTCGGGTTGCTTGGGAATGCA
1 GTAGCGGCCCCCGGCGCGCCGGGCCCGGGTCTTCCCGGAGTCGGGTTGCTTGGGAATGCA 300
2 GTAGCGGCCCCCGGCGCGCCGGGCCCGGGTCTTCCCGGAGTCGGGTTGCTTGGGAATGCA 300
3 GTAGCGGCCCCCGGCGCGCCGGGCCCGGGTCTTCCCGGAGTCGGGTTGCTTGGGAATGCA 300
```

GCCCAAAGCGGGTGGTAAACTCCATCTAAGGCTAAATACCGGCACGAGACCGATAGCCAA

1 GCCCAAAGCGGGTGGTAAACTCCATCTAAGGCTAAATACCGGCACGAGACCGATAGTCAA 360
2 GCCCAAAGCGGGTGGTAAACTCCATCTAAGGCTAAATACCGGCACGAGACCGATAGCCAA 360
3 GCCCAAAGCGGGTGGTAAACTCCATCTAAGGCTAAATACCGGCACGAGACCGATAGCCAA 360

CAAGTACCGTAAGGGAAAGTTGAAAAGAACTTTGAAGAGAGAGTTCAAGAGGGCGTGAAA

1 CAAGTACCGTAAGGGAAAGTTGAAAAGAACTTTGAAGAGAGAGTTCAAGAGGGCGTGAAA 420
2 CAAGTACCGTAAGGGAAAGTTGAAAAGAACTTTGAAGAGAGAGTTCAAGAGGGCGTGAAA 420
3 CAAGTACCGTAAGGGAAAGTTGAAAAGAACTTTGAAGAGAGAGTTCAAGAGGGCGTGAAA 420

CCGTTAAGAGGTAAACGGGTGGGGTCCGCGCAGTCCGCCCGGAGGATTCAACCCGGCGGC

1 CCGTTAAGAGGTAAACGGGTGGGGTCCGCGCAGTCCGCCCGGAGGATTCAACCCGGCG-- 478
2 CCGTTAAGAGGTAAACGGGTGGGGTCCGCGCAGTCCGCCCGGAGGATTCAACCCGGCGGC 480
3 CCGTTAAGAGGTAAACGGGTGGGGTCCGCGCAGTCCGCCCGGAGGATTCAACCCGGCGGC 480

GCGCGTCCGGCCGTGCCGGTGGTCCCGGCGGATCTTTCCCGCTCCCCGTTCCTCCCGACC

1 GCGGGTCCGGCCGTGTCGGCGG-CCCGGCGGATCTTTCCCGCCCCCGTTCCTCCCGACC 537
2 GCGCGTCCGGCCGTGCCGGTGGTCCCGGCGGATCTTTCCCGCTCCCCGTTCCTCCCGACC 540
3 GCGCGTCCGGCCGTGCCGGTGGTCCCGGCGGATCTTTCCCGCTCCCCGTTCCTCCCGACC 540

CCTCCACCCGCGCGTCTCTCTCCCCCCTCCCCGCGTCCCGCCGTCGCCGTCCCCGCTCCT

1 CCTCCACCCGC----------CCTCCCTTCCC-----------CCGCCGCCCCTCCTCCT 576
2 CCTCCACCCGCGCGTCTCTCTCCCCCCTCCCCGCGTCCCGCCGTCGCCGTCCCCGCTCCT 600
3 CCTCCACCCGCGCGTCTCTCTCCCCCCTCCCCGCGTCCCGCCGTCGCCGTCCCCGCTCCT 600

CCCTCCGGGGGGGTGTCGGCGGGCGCTCCGGCGGCGGGCGCGGGGTGTGGTGGGGGCGCG

1 CCTCCCCGGAGGG----GGCGGGCTC-----CGGCGGGTGC----------GGGGGTGGG 617
2 CCCTCCGGGGGGGTGTCGGCGGGCGCTCCGGCGGCGGGCGCGGGGTGTGGTGGGGGCGCG 660
3 CCCTCCGGGGGGGTGTCGGCGGGCGCTCCGGCGGCGGGCGCGGGGTGTGGTGGGGGCGCG 660

```
    CGGGCGGGCCGGGGTGGGTCGGCGGGGACCGNCCCCCGGTCGGCGACCGGCCGCCG
1   CGGGCGGGCCGGGGTGGGTCGGCGGGGACCGTCCCCCGACCGGCGACCGGCCGCCG   677
2   CGGGCGGGCCGGGGTGGGTCGGCGGGGACCGCCCCCCGGTCGGCGACCGGCCGCCG   720
3   CGGGCGGGCCGGGGTGGGTCGGCGGGGACCG-CCCCCGGTCGGCGACCGGCCGCCG   719
```

```
    CCGGGCGCACTTCCACCGTGGCGGTGCGCCGCGACCGGCTCCGGGACGGCTGGGAAGGCC
1   CCGGGCGCATTTCCACCGCGGCGGTGCGCCGCGACCGGCTCCGGGACGGCTGGGAAGGCC   737
2   CCGGGCGCACTTCCACCGTGGCGGTGCGCCGCGACCGGCTCCGGGACGGCTGGGAAGGCC   780
3   CCGGGCGCACTTCCACCGTGGCGGTGCGCCGCGACCGGCTCCGGGACGGCTGGGAAGGCC   779
```

```
    CGGCGGGGAAGGTGGCTCGGGGGG------------------------------------
1   CGGCGGGGAAGGTGGCTCGGGGGGCCCCGTCCGTCCGTCCGTCCGTCCTCCTCCTCCCCC   797
2   CGGCGGGGAAGGTGGCTCGGGGGG------------------------------------   804
3   CGGCGGGGAAGGTGGCTCGGGGGG------------------------------------   803
```

```
    ------------------------------------------------------------
1   GTCTCCGCCCCCCGGCCCCGCGTCCTCCCTCGGGAGGGCGCGCGGGTCGGGGCGGCGGCG   857
2   ------------------------------------------------------------   804
3   ------------------------------------------------------------   803
```

```
    -----GGCGGCGTCACCCGTGGGCGCCGGACC--ACCCCGCCCCGAGTGTTACAG-CCCC
1   GCGGTGGCGGCGGCGGCGGCGGCGGCGGACCGAAACCCCCCCCGAGTGTTACAGCCCCC   917
2   -----GGCGGCGTCACCCGTGGGCGCCGGACC--ACCCCGCCCCGAGTGTTACAG-CCCC   856
3   -----GGCGGCGTCACCCGTGGGCGCCGGACC--ACCCCGCCCCGAGTGTTACAG-CCCC   855
```

```
    CCGGCAGCAGCGCTCGCCGAATCCCGGGGCCGAGGGAGCCGGATACCCGTCGCCGCGCTC
1   CCGGCAGCAGCACTCGCCGAATCCCGGGGCCGAGGGAGC--GAGACCCGTCGCCGCGCTC   975
2   CCGGCAGCAGCGCTCGCCGAATCCCGGGGCCGAGGGAGCCGGATACCCGTCGCCGCGCTC   916
3   CCGGCAGCAGCGCTCGCCGAATCCCGGGGCCGAGGGAGCCGGATACCCGTCGCCGCGCTC   915
```

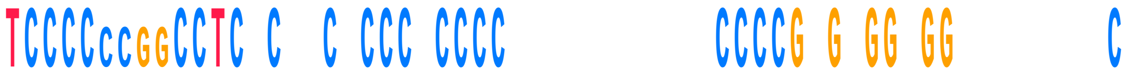

```
  TCCCCCCGGCCTCTCCCCTCCCGCCCC----------TCCCCGTGGGGTGG--------C
1 TCCCC----CCTCCCGGCGCCCACCCCCGCGGGGAATCCCCCGCGAGGGGGGTCTCCCCC 1031
2 TCCCCCCGGCCTCTCCCCTCCCGCCCC----------TCCCCGTGGGGTGG--------C 958
3 TCCCCCCGGCCTCTCCCCTCCCGCCCC----------TCCCCGTGGGGTGG--------C 957
```

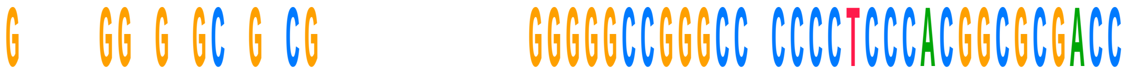

```
  GGAAAGGGGGGCGGTCGC----------GGGGGCCGGGCCGCCCCTCCCACGGCGCGACC
1 GCGGGGGCGCGCCGGCGTCTCCTCGTGGGGGGGCCGGGCCACCCCTCCCACGGCGCGACC 1091
2 GGAAAGGGGGGCGGTCGC----------GGGGGCCGGGCCGCCCCTCCCACGGCGCGACC 1008
3 GGAAAGGGGGGCGGTCGC----------GGGGGCCGGGCCGCCCCTCCCACGGCGCGACC 1007
```

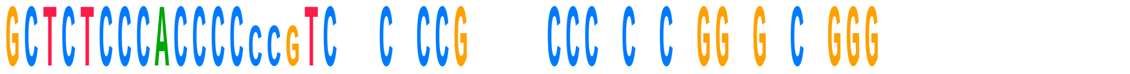

```
  GCTCTCCCACCCCCCGTCGCCTCCGTCGTCCCTCTCGGGGGTCCGGG-------------
1 GCTCTCCCACCCC---TCCTCCCCGCGCCCCCGCCCCGGCGACGGGGGGGGTGCCGCGCG 1148
2 GCTCTCCCACCCCCCGTCGCCTCCGTCGTCCCTCTCGGGGGTCCGGG------------- 1055
3 GCTCTCCCACCCCCCGTCGCCTCCGTCGTCCCTCTCGGGGGTCCGGG------------- 1054
```

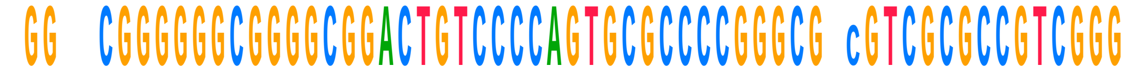

```
  -GGCCCGGGGGGCGGGGCGGACTGTCCCCAGTGCGCCCCGGGCGTCGTCGCGCCGTCGGG
1 CGGGTCGGGGGGCGGGGCGGACTGTCCCCAGTGCGCCCCGGGCGG-GTCGCGCCGTCGGG 1207
2 -GGCCCGGGGGGCGGGGCGGACTGTCCCCAGTGCGCCCCGGGCGTCGTCGCGCCGTCGGG 1114
3 -GGCCCGGGGGGCGGGGCGGACTGTCCCCAGTGCGCCCCGGGCGTCGTCGCGCCGTCGGG 1113
```

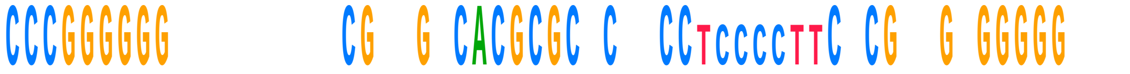

```
  CCCGGGGGGG-------CCGTCGTCACGCGCTCTCCCTCCCCTTCTCGGGGTGGGGGGGA
1 CCCGGGGGGAGGTTCTCTCGGGGCCACGCGCGCGTCC-------CCCGAAGAGGGGGACG 1260
2 CCCGGGGGGG-------CCGTCGTCACGCGCTCTCCCTCCCCTTCTCGGGGTGGGGGGGA 1167
3 CCCGGGGGGG-------CCGTCGTCACGCGCTCTCCCTCCCCTTCTCGGGGTGGGGGGGA 1166
```

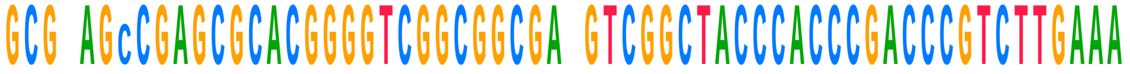

```
  GCGAAGCCGAGCGCACGGGGTCGGCGGCGATGTCGGCTACCCACCCGACCCGTCTTGAAA
1 GCGGAG-CGAGCGCACGGGGTCGGCGGCGACGTCGGCTACCCACCCGACCCGTCTTGAAA 1319
2 GCGAAGCCGAGCGCACGGGGTCGGCGGCGATGTCGGCTACCCACCCGACCCGTCTTGAAA 1227
3 GCGAAGCCGAGCGCACGGGGTCGGCGGCGATGTCGGCTACCCACCCGACCCGTCTTGAAA 1226
```

```
    CACGGACCAAGGAGTCTAAC CGTGCGCGAGTC GGGGCTCG  CGAAAGCCGCCGTGGC
    CACGGACCAAGGAGTCTAACGCGTGCGCGAGTCAGGGGCTCGTCCGAAAGCCGCCGTGGC
1   CACGGACCAAGGAGTCTAACACGTGCGCGAGTCGGGGGCTCGCACGAAAGCCGCCGTGGC   1379
2   CACGGACCAAGGAGTCTAACGCGTGCGCGAGTCAGGGGCTCGTCCGAAAGCCGCCGTGGC   1287
3   CACGGACCAAGGAGTCTAACGCGTGCGCGAGTCAGGGGCTCGTCCGAAAGCCGCCGTGGC   1286

    GCAATGAAGGTGAAGG C  CG  C CG  GGccCCGAGGTGGGATCCGAGGCCTCTCC
    GCAATGAAGGTGAAGGGCCCCGTTCCCGGGGGCCCCGAGGTGGGATCCGAGGCCTCTCC
1   GCAATGAAGGTGAAGGCCGGCGCGCTCGCCGG--CCGAGGTGGGATCCGAGGCCTCTCC   1437
2   GCAATGAAGGTGAAGGGCCCCGTTCCCGGGGGCCCCGAGGTGGGATCCGAGGCCTCTCC   1347
3   GCAATGAAGGTGAAGGGCCCCGTTCCCGGGGGCCCCGAGGTGGGATCCGAGGCCTCTCC   1346

    AGTCCGCCGAGGGCGCACCACCGGCCCGTCTCGCCCGCCGCGCCGGGGAGGTGGAGCACG
    AGTCCGCCGAGGGCGCACCACCGGCCCGTCTCGCCCGCCGCGCCGGGGAGGTGGAGCACG
1   AGTCCGCCGAGGGCGCACCACCGGCCCGTCTCGCCCGCCGCGCCGGGGAGGTGGAGCACG   1497
2   AGTCCGCCGAGGGCGCACCACCGGCCCGTCTCGCCCGCCGCGCCGGGGAGGTGGAGCACG   1407
3   AGTCCGCCGAGGGCGCACCACCGGCCCGTCTCGCCCGCCGCGCCGGGGAGGTGGAGCACG   1406

    AGCG ACG GTTAGGACCCGAAAGATGGTGAACTATGCCTGGGCAGGGCGAAGCCAGAGG
    AGCGTACGCGTTAGGACCCGAAAGATGGTGAACTATGCCTGGGCAGGGCGAAGCCAGAGG
1   AGCGCACGTGTTAGGACCCGAAAGATGGTGAACTATGCCTGGGCAGGGCGAAGCCAGAGG   1557
2   AGCGTACGCGTTAGGACCCGAAAGATGGTGAACTATGCCTGGGCAGGGCGAAGCCAGAGG   1467
3   AGCGTACGCGTTAGGACCCGAAAGATGGTGAACTATGCCTGGGCAGGGCGAAGCCAGAGG   1466

    AAACTCTGGTGGAGGTCCGTAGCGGTCCTGACGTGCAAATCGGTCGTCCGACCTGGGTAT
    AAACTCTGGTGGAGGTCCGTAGCGGTCCTGACGTGCAAATCGGTCGTCCGACCTGGGTAT
1   AAACTCTGGTGGAGGTCCGTAGCGGTCCTGACGTGCAAATCGGTCGTCCGACCTGGGTAT   1617
2   AAACTCTGGTGGAGGTCCGTAGCGGTCCTGACGTGCAAATCGGTCGTCCGACCTGGGTAT   1527
3   AAACTCTGGTGGAGGTCCGTAGCGGTCCTGACGTGCAAATCGGTCGTCCGACCTGGGTAT   1526

    AGGGGCGAAAGACTAATCGAACCATCTAGTAGCTGGTTCCCTCCGAAGTTTCCCTCAGGA
    AGGGGCGAAAGACTAATCGAACCATCTAGTAGCTGGTTCCCTCCGAAGTTTCCCTCAGGA
1   AGGGGCGAAAGACTAATCGAACCATCTAGTAGCTGGTTCCCTCCGAAGTTTCCCTCAGGA   1677
2   AGGGGCGAAAGACTAATCGAACCATCTAGTAGCTGGTTCCCTCCGAAGTTTCCCTCAGGA   1587
3   AGGGGCGAAAGACTAATCGAACCATCTAGTAGCTGGTTCCCTCCGAAGTTTCCCTCAGGA   1586
```

```
TAGCTGGCGCTCTCGCAACGCGGTCGCTCGACAACC-CGCAGTTTTATCCGGTAAAGCGA

1   TAGCTGGCGCTCTCGCAGACCCGACGCACCCCCGCCACGCAGTTTTATCCGGTAAAGCGA   1737
2   TAGCTGGCGCTCTCGCAACGCGGTCGCTCGACAACC-CGCAGTTTTATCCGGTAAAGCGA   1646
3   TAGCTGGCGCTCTCGCAACGCGGTCGCTCGACAACC-CGCAGTTTTATCCGGTAAAGCGA   1645
```

```
ATGATTAGAGGTCTTGGGGCCGAAACGATCTCAACCTATTCTCAAACTTTAAATGGGTAA

1   ATGATTAGAGGTCTTGGGGCCGAAACGATCTCAACCTATTCTCAAACTTTAAATGGGTAA   1797
2   ATGATTAGAGGTCTTGGGGCCGAAACGATCTCAACCTATTCTCAAACTTTAAATGGGTAA   1706
3   ATGATTAGAGGTCTTGGGGCCGAAACGATCTCAACCTATTCTCAAACTTTAAATGGGTAA   1705
```

```
GAAGCCCGGCTCGCTGGCGTGGAGCCGGGCGTGGAATGCGAGTGCCTAGTGGGCCACTTT

1   GAAGCCCGGCTCGCTGGCGTGGAGCCGGGCGTGGAATGCGAGTGCCTAGTGGGCCACTTT   1857
2   GAAGCCCGGCTCGCTGGCGTGGAGCCGGGCGTGGAATGCGAGTGCCTAGTGGGCCACTTT   1766
3   GAAGCCCGGCTCGCTGGCGTGGAGCCGGGCGTGGAATGCGAGTGCCTAGTGGGCCACTTT   1765
```

```
TGGTAAGCAGAACTGGCGCTGCGGGATGAACCGAACGCCGGGTTAAGGCGCCCGATGCCG

1   TGGTAAGCAGAACTGGCGCTGCGGGATGAACCGAACGCCGGGTTAAGGCGCCCGATGCCG   1917
2   TGGTAAGCAGAACTGGCGCTGCGGGATGAACCGAACGCCGGGTTAAGGCGCCCGATGCCG   1826
3   TGGTAAGCAGAACTGGCGCTGCGGGATGAACCGAACGCCGGGTTAAGGCGCCCGATGCCG   1825
```

```
ACGCTCATCAGACCCCAG-AAAAGGTGTTGGTTGATATAGACAGCAGGACGGTGGCCATG

1   ACGCTCATCAGACCCCAG-AAAAGGTGTTGGTTGATATAGACAGCAGGACGGTGGCCATG   1976
2   ACGCTCATCAGACCCCAG-AAAAGGTGTTGGTTGATATAGACAGCAGGACGGTGGCCATG   1885
3   ACGCTCATCAGACCCCAGAAAAAGGTGTTGGTTGATATAGACAGCAGGACGGTGGCCATG   1885
```

```
GAAGTCGGAATCCGCTAAGGAGTGTGTAACAACTCACCTGCCGAATCAACTAGCCCTGAA

1   GAAGTCGGAATCCGCTAAGGAGTGTGTAACAACTCACCTGCCGAATCAACTAGCCCTGAA   2036
2   GAAGTCGGAATCCGCTAAGGAGTGTGTAACAACTCACCTGCCGAATCAACTAGCCCTGAA   1945
3   GAAGTCGGAATCCGCT-AGGAGTGTGTAACAACTCACCTGCCGAATCAACTAGCCCTGAA   1944
```

```
  AATGGATGGCGCTGGAGCGTCGGGCCCATACCCGGCCGTCGCCGGCAGTCGGAACGGGAC
1 AATGGATGGCGCTGGAGCGTCGGGCCCATACCCGGCCGTCGCCGGCAGTCGAGAGTGGAC 2096
2 AATGGATGGCGCTGGAGCGTCGGGCCCATACCCGGCCGTCGCCGGCAGTCGGAACGGGAC 2005
3 AATGGATGGCGCTGGAGCGTCGGGCCCATACCCGGCCGTCGCCGGCAGTCGGAACGGGAC 2004

  GGGAGCGGC--------CGCGGGCGCGCGACCCCGGGG-----GCCGGGCGGCGTCGGCT
1 GGGAGCGGCGGGGGCGGCGCGCGCGCGCGCGTGTGGTGTGCGTCGGAGGCGGGCGGCG 2156
2 GGGAGCGGC--------CGCGGGCGCGCGACCCCGGGG-----GCCGGGCGGCGTCGGCT 2052
3 GGGAGCGGC--------CGCGGGCGCGCGACCCCGGGG-----GCCGGGCGGCGTCGGCT 2051

  TCGGCCGG------------------------------------------CCGCCGCCCG
1 GCGGCGGGGGTGTGTGGGGTCCTCCCCCGCCCCCCCCCCACGCCTCCTCCCCTCCTCCCG 2216
2 TCGGCCGG------------------------------------------CCGCCGCCCG 2070
3 TCGGCCGG------------------------------------------CCGCCGCCCG 2069

  TCCACCCCCGGGGCTCCCCCCGCGGCGTCGGGCCCCGCGGACGCTACGCCGCGACGAGTA
1 CCCACGCCCCGCTCCCCGCCCCCG-----GAGCCCCGCGGACGCTACGCCGCGACGAGTA 2271
2 TCCACCCCCGGGGCTCCCCCCGCGGCGTCGGGCCCCGCGGACGCTACGCCGCGACGAGTA 2130
3 TCCACCCCCGGGGCTCCCCCCGCGGCGTCGGGCCCCGCGGAGCCTACGCCGCGACGAGTA 2129

  GGAGGGCCGCTGCGGTGAGCCTTGAAGCCTAGGGCGCGGGCCCGGGTGGAGCCGCCGCAG
1 GGAGGGCCGCTGCGGTGAGCCTTGAAGCCTAGGGCGCGGGCCCGGGTGGAGCCGCCGCAG 2331
2 GGAGGGCCGCTGCGGTGAGCCTTGAAGCCTAGGGCGCGGGCCCGGGTGGAGCCGCCGCAG 2190
3 GGAGGGCCGCTGCGGTGAGCCTTGAAGCCTAGGGCGCGGGCCCGGGTGGAGCCGCCGCAG 2189

  GTGCAGATCTTGGTGGTAGTAGCAAATATTCAAACGAGAACTTTGAAGGCCGAAGTGGAG
1 GTGCAGATCTTGGTGGTAGTAGCAAATATTCAAACGAGAACTTTGAAGGCCGAAGTGGAG 2391
2 GTGCAGATCTTGGTGGTAGTAGCAAATATTCAAACGAGAACTTTGAAGGCCGAAGTGGAG 2250
3 GTGCAGATCTTGGTGGTAGTAGCAAATATTCAAACGAGAACTTTGAAGGCCGAAGTGGAG 2249
```

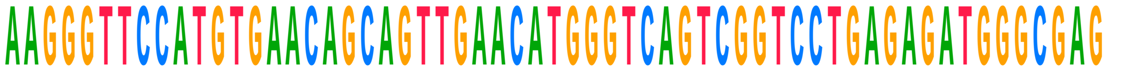

| | | |
|---|---|---|
| | AAGGGTTCCATGTGAACAGCAGTTGAACATGGGTCAGTCGGTCCTGAGAGATGGGCGAGT | |
| 1 | AAGGGTTCCATGTGAACAGCAGTTGAACATGGGTCAGTCGGTCCTGAGAGATGGGCGAGC | 2451 |
| 2 | AAGGGTTCCATGTGAACAGCAGTTGAACATGGGTCAGTCGGTCCTGAGAGATGGGCGAGT | 2310 |
| 3 | AAGGGTTCCATGTGAACAGCAGTTGAACATGGGTCAGTCGGTCCTGAGAGATGGGCGAGT | 2309 |

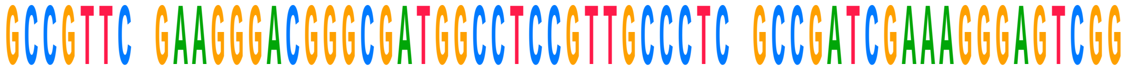

| | | |
|---|---|---|
| | GCCGTTCCGAAGGGACGGGCGATGGCCTCCGTTGCCCTCAGCCGATCGAAAGGGAGTCGG | |
| 1 | GCCGTTCTGAAGGGACGGGCGATGGCCTCCGTTGCCCTCGGCCGATCGAAAGGGAGTCGG | 2511 |
| 2 | GCCGTTCCGAAGGGACGGGCGATGGCCTCCGTTGCCCTCAGCCGATCGAAAGGGAGTCGG | 2370 |
| 3 | GCCGTTCCGAAGGGACGGGCGATGGCCTCCGTTGCCCTCAGCCGATCGAAAGGGAGTCGG | 2369 |

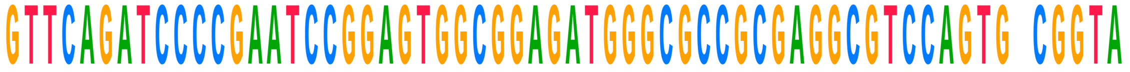

| | | |
|---|---|---|
| | GTTCAGATCCCCGAATCCGGAGTGGCGGAGATGGGCGCCGCGAGGCGTCCAGTG-CGGTA | |
| 1 | GTTCAGATCCCCGAATCCGGAGTGGCGGAGATGGGCGCCGCGAGGCGTCCAGTG-CGGTA | 2570 |
| 2 | GTTCAGATCCCCGAATCCGGAGTGGCGGAGATGGGCGCCGCGAGGCGTCCAGTG-CGGTA | 2429 |
| 3 | GTTCAGATCCCCGAATCCGGAGTGGCGGAGATGGGCGCCGCGAGGCGTCCAGTGCCGGTA | 2429 |

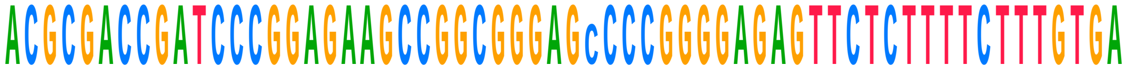

| | | |
|---|---|---|
| | ACGCGACCGATCCCGGAGAAGCCGGCGGGAGCCCCGGGGAGAGTTCTCTTTTCTTTGTGA | |
| 1 | ACGCGACCGATCCCGGAGAAGCCGGCGGGAGCCCCGGGGAGAGTTCTCTTTTCTTTGTGA | 2630 |
| 2 | ACGCGACCGATCCCGGAGAAGCCGGCGGGAGCCCCGGGGAGAGTTCTCTTTTCTTTGTGA | 2489 |
| 3 | ACGCGACCGATCCCGGAGAAGCCGGCGGGAG-CCCGGGGAGAGTTCTCTTTTCTTTGTGA | 2488 |

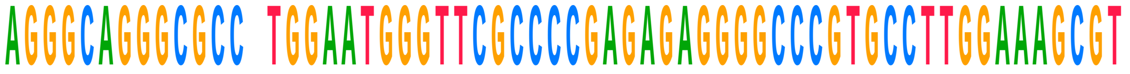

| | | |
|---|---|---|
| | AGGGCAGGGCGCCCTGGAATGGGTTCGCCCCGAGAGAGGGGCCCGTGCCTTGGAAAGCGT | |
| 1 | AGGGCAGGGCGCCCTGGAATGGGTTCGCCCCGAGAGAGGGGCCCGTGCCTTGGAAAGCGT | 2690 |
| 2 | AGGGCAGGGCGCCCTGGAATGGGTTCGCCCCGAGAGAGGGGCCCGTGCCTTGGAAAGCGT | 2549 |
| 3 | AGGGCAGGGCGCCTTGGAATGGGTTCGCCCCGAGAGAGGGGCCCGTGCCTTGGAAAGCGT | 2548 |

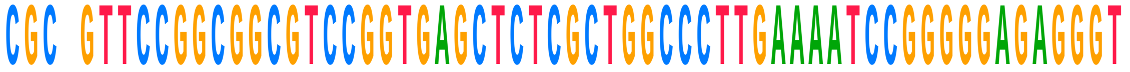

| | | |
|---|---|---|
| | CGCGGTTCCGGCGGCGTCCGGTGAGCTCTCGCTGGCCCTTGAAAATCCGGGGGAGAGGGT | |
| 1 | CGCGGTTCCGGCGGCGTCCGGTGAGCTCTCGCTGGCCCTTGAAAATCCGGGGGAGAGGGT | 2750 |
| 2 | CGCGGTTCCGGCGGCGTCCGGTGAGCTCTCGCTGGCCCTTGAAAATCCGGGGGAGAGGGT | 2609 |
| 3 | CGCAGTTCCGGCGGCGTCCGGTGAGCTCTCGCTGGCCCTTGAAAATCCGGGGGAGAGGGT | 2608 |

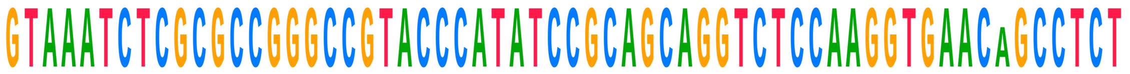

GTAAATCTCGCGCCGGGCCGTACCCATATCCGCAGCAGGTCTCCAAGGTGAACAGCCTCT

1 GTAAATCTCGCGCCGGGCCGTACCCATATCCGCAGCAGGTCTCCAAGGTGAACAGCCTCT 2810
2 GTAAATCTCGCGCCGGGCCGTACCCATATCCGCAGCAGGTCTCCAAGGTGAACAGCCTCT 2669
3 GTAAATCTCGCGCCGGGCCGTACCCATATCCGCAGCAGGTCTCCAAGGTGAAC-GCCTCT 2667

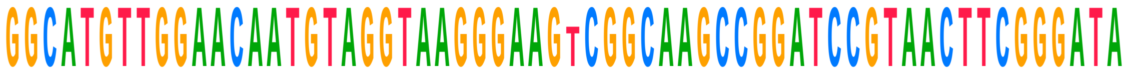

GGCATGTTGGAACAATGTAGGTAAGGGAAGTCGGCAAGCCGGATCCGTAACTTCGGGATA

1 GGCATGTTGGAACAATGTAGGTAAGGGAAGTCGGCAAGCCGGATCCGTAACTTCGGGATA 2870
2 GGCATGTTGGAACAATGTAGGTAAGGGAAGTCGGCAAGCCGGATCCGTAACTTCGGGATA 2729
3 GGCATGTTGGAACAATGTAGGTAAGGGAAG-CGGCAAGCCGGATCCGTAACTTCGGGATA 2726

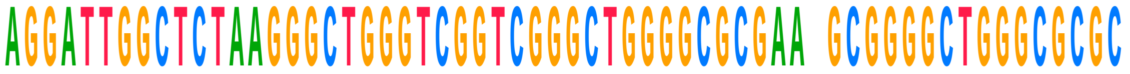

AGGATTGGCTCTAAGGGCTGGGTCGGTCGGGCTGGGGCGCGAA-GCGGGGCTGGGCGCGC

1 AGGATTGGCTCTAAGGGCTGGGTCGGTCGGGCTGGGGCGCGAA-GCGGGGCTGGGCGCGC 2929
2 AGGATTGGCTCTAAGGGCTGGGTCGGTCGGGCTGGGGCGCGAA-GCGGGGCTGGGCGCGC 2788
3 AGGATTGGCTCTAAGGGCTGGGTCGGTCGGGCTGGGGCGCGAAGGCGGGGCTGGGCGCGC 2786

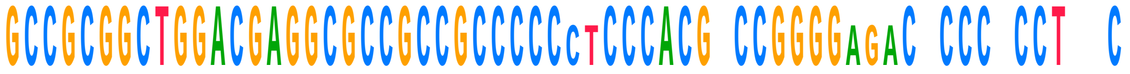

GCCGCGGCTGGACGAGGCGCCGCCGCCCCCCTCCCACGTCCGGGGAGACCCCCTCCTTTC

1 GCCGCGGCTGGACGAGGCGCCGCCGCCCCC--CCCACGCCCGGGG---CACCCCCCTCGC 2984
2 GCCGCGGCTGGACGAGGCGCCGCCGCCCCCCTCCCACGTCCGGGGAGACCCCCTCCTTTC 2848
3 GCCGCGGCTGGACGAGGCGCCGCCGCCCCCCTCCCACGTCCGGGGAGACCCCCTCCTTTC 2846

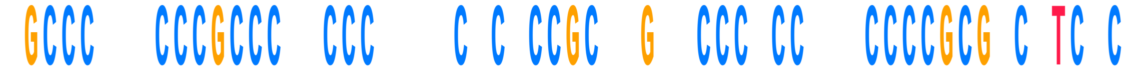

CGCCCGGGCCCGCCCTCCCCT---CTCCCCGCGGGGCCCCGCCGTCCCCCGCGTCGTCGC

1 GGCCCTCCCCCGCCCCACCCCGCGCGCGCCGCTCGCTCCCTCCCCACCCCGCGCCCTCTC 3044
2 CGCCCGGGCCCGCCCTCCCCT---CTCCCCGCGGGGCCCCGCCGTCCCCCGCGTCGTCGC 2905
3 CGCCCGGGCCCGCCCTCCCCT---CTCCCCGCGGGGCCCCGCCGTCCCCCGCGTCGTCGC 2903

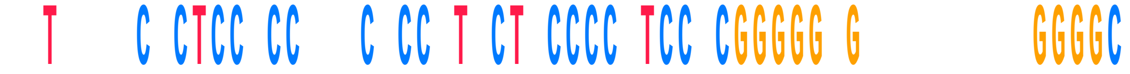

CGTGGTCCCCTCCTCC---CTCCCTTCTTCCCCGTCCGCGGGGGGGA-------CGGGGC

1 TCTCTCTCTCTCCCCCGCTCCCCGTCCTCCCCCCTCCCCGGGGGAGCGCCGCGTGGGGGC 3104
2 CGTGGTCCCCTCCTCC---CTCCCTTCTTCCCCGTCCGCGGGGGGGA-------CGGGGC 2955
3 CGTGGTCCCCTCCTCC---CTCCCTTCTTCCCCGTCCGCGGGGGGGA-------CGGGGC 2953

```
GGGTGCGGGGGGGCGCGCGCGCGCGG-CCCAGGGG-CGGCGGGTCCAACCCCGCGCGG

1  GGCGGCGGGGGGAGAAGGGTCGGGGCGG---CAGGGGCCGGCGG--CGGCCGCCGCG-GG  3158
2  GGGTGCGGGGGGGCGCGCGCGCGCGGGCCCAGGGG-CGGCGGGTCCAACCCCGCGCGG  3014
3  GGGTGCGGGGGGGCGCGCGCGCGCGG-CCCAGGGG-CGGCGGGTCCAACCCCGCGCGG  3011

GCCGGAGCGGGGGGAACCCGCGGGCCCCCGGTGGGGGGGGGCCCGGACACCCGGGGGGGN

1  GCCCCGGCGGCGGGGG--CACGGTCCCCCG--CGAGGGGGGCCCGGGCACCCGGGGGG--  3212
2  GCCGGAGCGGGGGGAACCCGCGGGCCCCCGGTGGGGGGGGGCCCGGACACCCGGGGGGGG  3074
3  GCCGGAGCGGGGGGAACCCGCGGGCCCCCGGTGGGGGGGGGCCCGGACACCCGGGGGGGA  3071

CCGGCGGCGGCGGCGACTCTGGACGCGAGCCGGGCCCTTCCCGTGGATCGCCCCAGCTGC

1  CCGGCGGCGGCGGCGACTCTGGACGCGAGCCGGGCCCTTCCCGTGGATCGCCCCAGCTGC  3272
2  CCGGCGGCGGCGGCGACTCTGGACGCGAGCCGGGCCCTTCCCGTGGATCGCCCCAGCTGC  3134
3  CCGGCGGCGGCGGCGACTCTGGACGCGAGCCGGGCCCTTCCCGTGGATCGCCCCAGCTGC  3131

GGCGGGCGTCGCGGCCGCTCCCGGGGAGCCCGGCGGGTCGCCGGCGCGGGGTTTTCCTCC

1  GGCGGGCGTCGCGGCCGCCCCCGGGGAGCCCGGCGGG-CGCCGGCGCGNCCCCCCCCCCA  3331
2  GGCGGGCGTCGCGGCCGCTCCCGGGGAGCCCGGCGGGTCGCCGGCGCGGGGTTTTCCTCC  3194
3  GGCGGGCGTCGCGGCCGCTCCCGGGGAGCCCGGCGGGTCGCCGGCGCGGGGTTTTCCTCC  3191

GGCCTCGTCCTCCCCCTTCCCCCTCCGC-GGGGTCGGG------------------GGTT

1  CCCCACGTCTCGTCGCGCGCGCGTCCGCTGGGGGCGGGGAGCGGTCGGGCGGCGGCGGTC  3391
2  GGCCTCGTCCTCCCCCTTCCCCCTCCGC-GGGGTCGGG------------------GGTT  3235
3  GGCCTCGTCCTCCCCCTTCCCCCTCCGC-GGGGTCGGG------------------GGTT  3232

CCCGGGGTTCGGGG----------------------------------------------

1  GGCGGGCGGCGGGGCGGGGCGGTTCGTCCCCCCGCCCTACCCCCCCGGCCCCGTCCGCCC  3451
2  CCCGGGGTTCGGGG----------------------------------------------  3249
3  CCCGGGGTTCGGGG----------------------------------------------  3246
```

```
            T CTCCTC    CGCG CGGCGG
------------TTCTCCTC----CGCGTCGGCGG-------------------------
1  CCCGTTCCCCCCTCCTCCTCGGCGCGCGGCGGCGGCGGCGGGCGGCGGAGGGGCCGCGGG  3511
2  ------------TTCTCCTC----CGCGTCGGCGG-------------------------  3268
3  ------------TTCTCCTC----CGCGTCGGCGG-------------------------  3265

       CCCCCGCCGGGT CGCCCCC GGGCCGCGGTTTcCCGCGCGGCGCCTCGCCTC
-----TTCCCCCGCCGGGTGCGCCCCCCGGGCCGCGGTTTCCCGCGCGGCGCCTCGCCTC
1  CCGGTCCCCCCCGCCGGGTCCGCCCCCGGGGCCGCGGTT--CCGCGCGGCGCCTCGCCTC  3569
2  -----TTCCCCCGCCGGGTGCGCCCCCCGGGCCGCGGTTTCCCGCGCGGCGCCTCGCCTC  3323
3  -----TTCCCCCGCCGGGTGCGCCCCCCGGGCCGCGGTTTCCCGCGCGGCGCCTCGCCTC  3320

GGCCGGCGCCTAGCAGCCGACTTAGAACTGGTGCGGACCAGGGGAATCCGACTGTTTAAT
GGCCGGCGCCTAGCAGCCGACTTAGAACTGGTGCGGACCAGGGGAATCCGACTGTTTAAT
1  GGCCGGCGCCTAGCAGCCGACTTAGAACTGGTGCGGACCAGGGGAATCCGACTGTTTAAT  3629
2  GGCCGGCGCCTAGCAGCCGACTTAGAACTGGTGCGGACCAGGGGAATCCGACTGTTTAAT  3383
3  GGCCGGCGCCTAGCAGCCGACTTAGAACTGGTGCGGACCAGGGGAATCCGACTGTTTAAT  3380

TAAAACAAAGCATCGCGAAGGCCCGCGGCGGGTGTTGACGCGATGTGATTTCTGCCCAGT
TAAAACAAAGCATCGCGAAGGCCCGCGGCGGGTGTTGACGCGATGTGATTTCTGCCCAGT
1  TAAAACAAAGCATCGCGAAGGCCCGCGGCGGGTGTTGACGCGATGTGATTTCTGCCCAGT  3689
2  TAAAACAAAGCATCGCGAAGGCCCGCGGCGGGTGTTGACGCGATGTGATTTCTGCCCAGT  3443
3  TAAAACAAAGCATCGCGAAGGCCCGCGGCGGGTGTTGACGCGATGTGATTTCTGCCCAGT  3440

GCTCTGAATGTCAAAGTGAAGAAATTCAATGAAGCGCGGGTAAACGGCGGGAGTAACTAT
GCTCTGAATGTCAAAGTGAAGAAATTCAATGAAGCGCGGGTAAACGGCGGGAGTAACTAT
1  GCTCTGAATGTCAAAGTGAAGAAATTCAATGAAGCGCGGGTAAACGGCGGGAGTAACTAT  3749
2  GCTCTGAATGTCAAAGTGAAGAAATTCAATGAAGCGCGGGTAAACGGCGGGAGTAACTAT  3503
3  GCTCTGAATGTCAAAGTGAAGAAATTCAATGAAGCGCGGGTAAACGGCGGGAGTAACTAT  3500

GACTCTCTTAAGGTAGCCAAATGCCTCGTCATCTAATTAGTGACGCGCATGAATGGATGA
GACTCTCTTAAGGTAGCCAAATGCCTCGTCATCTAATTAGTGACGCGCATGAATGGATGA
1  GACTCTCTTAAGGTAGCCAAATGCCTCGTCATCTAATTAGTGACGCGCATGAATGGATGA  3809
2  GACTCTCTTAAGGTAGCCAAATGCCTCGTCATCTAATTAGTGACGCGCATGAATGGATGA  3563
3  GACTCTCTTAAGGTAGCCAAATGCCTCGTCATCTAATTAGTGACGCGCATGAATGGATGA  3560
```

ACGAGATTCCCACTGTCCCTACCTACTATCCAGCGAAACCACAGCCAAGGGAACGGGCTT

1 ACGAGATTCCCACTGTCCCTACCTACTATCCAGCGAAACCACAGCCAAGGGAACGGGCTT 3869
2 ACGAGATTCCCACTGTCCCTACCTACTATCCAGCGAAACCACAGCCAAGGGAACGGGCTT 3623
3 ACGAGATTCCCACTGTCCCTACCTACTATCCAGCGAAACCACAGCCAAGGGAACGGGCTT 3620

GGCGGAATCAGCGGGGAAAGAAGACCCTGTTGAGCTTGACTCTAGTCTGGCACGGTGAAG

1 GGCGGAATCAGCGGGGAAAGAAGACCCTGTTGAGCTTGACTCTAGTCTGGCACGGTGAAG 3929
2 GGCGGAATCAGCGGGGAAAGAAGACCCTGTTGAGCTTGACTCTAGTCTGGCACGGTGAAG 3683
3 GGCGGAATCAGCGGGGAAAGAAGACCCTGTTGAGCTTGACTCTAGTCTGGCACGGTGAAG 3680

AGACATGAGAGGTGTAGAATAAGTGGGAGGCCCCCGGCGCCCCCCCG--TTCCCCGCGAG

1 AGACATGAGAGGTGTAGAATAAGTGGGAGGCCCCCGGCGCCCCCCCGGTGTCCCCGCGAG 3989
2 AGACATGAGAGGTGTAGAATAAGTGGGAGGCCCCCGGCGCCCCCCCG--TTCCCCGCGAG 3741
3 AGACATGAGAGGTGTAGAATAAGTGGGAGGCCCCCGGCGCCCCCCCG--TTCCCCGCGAG 3738

GGG-TCGGGGCGGGGTCCGCCGGCCTCGCGGGCCGCCGGTGAAATACCACTACTCTCATC

1 GGGCCCGGGGCGGGGTCCGCCGGCCCTGCGGGCCGCCGGTGAAATACCACTACTCTGATC 4049
2 GGG-TCGGGGCGGGGTCCGCCGGCCTCGCGGGCCGCCGGTGAAATACCACTACTCTCATC 3800
3 GGG-TCGGGGCGGGGTCCGCCGGCCTCGCGGGCCGCCGGTGAAATACCACTACTCTCATC 3797

GTTTTTTCACTGACCCGGTGAGGCGGGGGGGCGAGCCCCGAGGGGCTCTCGCTTCTGGCG

1 GTTTTTTCACTGACCCGGTGAGGCGGGGGGGCGAGCCCCGAGGGGCTCTCGCTTCTGGCG 4109
2 GTTTTTTCACTGACCCGGTGAGGCGGGGGGGCGAGCCCCGAGGGGCTCTCGCTTCTGGCG 3860
3 GTTTTTTCACTGACCCGGTGAGGCGGGGGGGCGAGCCCCGAGGGGCTCTCGCTTCTGGCG 3857

CCAAGCGCCCGTCCGCGCGCGCGGGCGGGCGCGACCCGCTCCGGGGACAGTGCCAGGTGG

1 CCAAGCGCCCGGCCGCGCGC-CGGCCGGGCGCGACCCGCTCCGGGGACAGTGCCAGGTGG 4168
2 CCAAGCGCCCGTCCGCGCGCGCGGGCGGGCGCGACCCGCTCCGGGGACAGTGCCAGGTGG 3920
3 CCAAGCGCCCGTCCGCGCGCGCGGGCGGGCGCGACCCGCTCCGGGGACAGTGCCAGGTGG 3917

| | | |
|---|---|---|
| | GGAGTTTGACTGGGGCGGTACACCTGTCAAACGGTAACGCAGGTGTCCTAAGGCGAGCTC | |
| 1 | GGAGTTTGACTGGGGCGGTACACCTGTCAAACGGTAACGCAGGTGTCCTAAGGCGAGCTC | 4228 |
| 2 | GGAGTTTGACTGGGGCGGTACACCTGTCAAACGGTAACGCAGGTGTCCTAAGGCGAGCTC | 3980 |
| 3 | GGAGTTTGACTGGGGCGGTACACCTGTCAAACGGTAACGCAGGTGTCCTAAGGCGAGCTC | 3977 |

| | | |
|---|---|---|
| | AGGGAGGACAGAAACCTCCCGTGGAGCAGAAGGGCAAAAGCTCGCTTGATCTTGATTTTC | |
| 1 | AGGGAGGACAGAAACCTCCCGTGGAGCAGAAGGGCAAAAGCTCGCTTGATCTTGATTTTC | 4288 |
| 2 | AGGGAGGACAGAAACCTCCCGTGGAGCAGAAGGGCAAAAGCTCGCTTGATCTTGATTTTC | 4040 |
| 3 | AGGGAGGACAGAAACCTCCCGTGGAGCAGAAGGGCAAAAGCTCGCTTGATCTTGATTTTC | 4037 |

| | | |
|---|---|---|
| | AGTACGAATACAGACCGTGAAAGCGGGGCCTCACGATCCTTCTGACCTTTTGGGTTTTAA | |
| 1 | AGTACGAATACAGACCGTGAAAGCGGGGCCTCACGATCCTTCTGACCTTTTGGGTTTTAA | 4348 |
| 2 | AGTACGAATACAGACCGTGAAAGCGGGGCCTCACGATCCTTCTGACCTTTTGGGTTTTAA | 4100 |
| 3 | AGTACGAATACAGACCGTGAAAGCGGGGCCTCACGATCCTTCTGACCTTTTGGGTTTTAA | 4097 |

| | | |
|---|---|---|
| | GCAGGAGGTGTCAGAAAAGTTACCACAGGGATAACTGGCTTGTGGCGGCCAAGCGTTCAT | |
| 1 | GCAGGAGGTGTCAGAAAAGTTACCACAGGGATAACTGGCTTGTGGCGGCCAAGCGTTCAT | 4408 |
| 2 | GCAGGAGGTGTCAGAAAAGTTACCACAGGGATAACTGGCTTGTGGCGGCCAAGCGTTCAT | 4160 |
| 3 | GCAGGAGGTGTCAGAAAAGTTACCACAGGGATAACTGGCTTGTGGCGGCCAAGCGTTCAT | 4157 |

| | | |
|---|---|---|
| | AGCGACGTCGCTTTTTGATCCTTCGATGTCGGCTCTTCCTATCATTGTGAAGCAGAATTC | |
| 1 | AGCGACGTCGCTTTTTGATCCTTCGATGTCGGCTCTTCCTATCATTGTGAAGCAGAATTC | 4468 |
| 2 | AGCGACGTCGCTTTTTGATCCTTCGATGTCGGCTCTTCCTATCATTGTGAAGCAGAATTC | 4220 |
| 3 | AGCGACGTCGCTTTTTGATCCTTCGATGTCGGCTCTTCCTATCATTGTGAAGCAGAATTC | 4217 |

| | | |
|---|---|---|
| | ACCAAGCGTTGGATTGTTCACCCACTAATAGGGAACGTGAGCTGGGTTTAGACCGTCGTG | |
| 1 | ACCAAGCGTTGGATTGTTCACCCACTAATAGGGAACGTGAGCTGGGTTTAGACCGTCGTG | 4528 |
| 2 | ACCAAGCGTTGGATTGTTCACCCACTAATAGGGAACGTGAGCTGGGTTTAGACCGTCGTG | 4280 |
| 3 | ACCAAGCGTTGGATTGTTCACCCACTAATAGGGAACGTGAGCTGGGTTTAGACCGTCGTG | 4277 |

```
  AGACAGGTTAGTTTTACCCTACTGATGATGTGTTGTTGCCATGGTAATCCTGCTCAGTAC
1 AGACAGGTTAGTTTTACCCTACTGATGATGTGTTGTTGCCATGGTAATCCTGCTCAGTAC 4588
2 AGACAGGTTAGTTTTACCCTACTGATGATGTGTTGTTGCCATGGTAATCCTGCTCAGTAC 4340
3 AGACAGGTTAGTTTTACCCTACTGATGATGTGTTGTTGCCATGGTAATCCTGCTCAGTAC 4337

  GAGAGGAACCGCAGGTTCAGACATTTGGTGTATGTGCTTGGCTGAGGAGCCAATGGGGCG
1 GAGAGGAACCGCAGGTTCAGACATTTGGTGTATGTGCTTGGCTGAGGAGCCAATGGGGCG 4648
2 GAGAGGAACCGCAGGTTCAGACATTTGGTGTATGTGCTTGGCTGAGGAGCCAATGGGGCG 4400
3 GAGAGGAACCGCAGGTTCAGACATTTGGTGTATGTGCTTGGCTGAGGAGCCAATGGGGCG 4397

  AAGCTACCATCTGTGGGATTATGACTGAACGCCTCTAAGTCAGAATCCCGCCCAGGCGGA
1 AAGCTACCATCTGTGGGATTATGACTGAACGCCTCTAAGTCAGAATCCCGCCCAGGCGGA 4708
2 AAGCTACCATCTGTGGGATTATGACTGAACGCCTCTAAGTCAGAATCCCGCCCAGGCGGA 4460
3 AAGCTACCATCTGTGGGATTATGACTGAACGCCTCTAAGTCAGAATCCCGCCCAGGCGGA 4457

  ACGATACGGCAGCGCCGAAGGAGCCTCGGTTGGCCCCGGATAGCCGGCTCCCCGTCCGTC
1 ACGATACGGCAGCGCCG-CGGAGCCTCGGTTGGCCTCGGATAGCCGGTCCCCCGCCTGTC 4767
2 ACGATACGGCAGCGCCGAAGGAGCCTCGGTTGGCCCCGGATAGCCGGCTCCCCGTCCGTC 4520
3 ACGATACGGCAGCGCCCAAGGAGCCTCGGTTGGCCCCGGATAGCCGGCTCCCCGTCCGTC 4517

  CCCGTCCGGCGG---GTCCCCGCCTCGTCGC-CCCCCCGGGTGCG---GGGCGGGTCCCC
1 CCCG-CCGGCGGGCCGCCCCCCCTCCACGCGCCCCGCGCGCGCGGGAGGGCGCGT-GCC 4825
2 CCCGTCCGGCGG---GTCCCCGCCTCGTCGC-CCCCCCGGGTGCG---GGGCGGGTCCCC 4573
3 CCCGTCCGGCGG---GTCCCCGCCTCGTCGC-CCCCCCGGGTGCG---GGGCGGGTCCCC 4570

  CCGCCGGGCGTCGGGACCGGGGTCCGGTGCGGAGAGCCATTCGTCCCGGGAAACGGGGTG
1 CCGCCGCGCGCCGGGACCGGGGTCCGGTGCGGAGTGCCCTTCGTCCTGGGAAACGGGGCG 4885
2 CCGCCGGGCGTCGGGACCGGGGTCCGGTGCGGAGAGCCATTCGTCCCGGGAAACGGGGTG 4633
3 CCGCCGGGCGTCGGGACCGGGGTCCGGTGCGGAGAGCCATTCGTCCCGGGAAACGGGGTG 4630
```

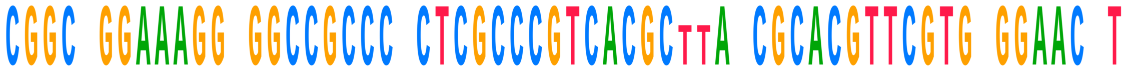

| | | |
|---|---|---|
| | CGGCCGGAAAGGGGGCCGCCCTCTCGCCCGTCACGCTTAACGCACGTTCGTGTGGAACCT | |
| 1 | CGGCTGGAAAGGCGGCCGCCCCCTCGCCCGTCACGC--ACCGCACGTTCGTGGGGAACCT | 4943 |
| 2 | CGGCCGGAAAGGGGGCCGCCCTCTCGCCCGTCACGCTTAACGCACGTTCGTGTGGAACCT | 4693 |
| 3 | CGGCCGGAAAGGGGGCCGCCCTCTCGCCCGTCACGCTTAACGCACGTTCGTGTGGAACTT | 4690 |

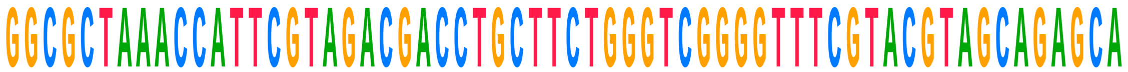

| | | |
|---|---|---|
| | GGCGCTAAACCATTCGTAGACGACCTGCTTCTGGGTCGGGGTTTCGTACGTAGCAGAGCA | |
| 1 | GGCGCTAAACCATTCGTAGACGACCTGCTTCTGGGTCGGGGTTTCGTACGTAGCAGAGCA | 5003 |
| 2 | GGCGCTAAACCATTCGTAGACGACCTGCTTCTGGGTCGGGGTTTCGTACGTAGCAGAGCA | 4753 |
| 3 | GGCGCTAAACCATTCGTAGACGACCTGCTTCTGGGTCGGGGTTTCGTACGTAGCAGAGCA | 4750 |

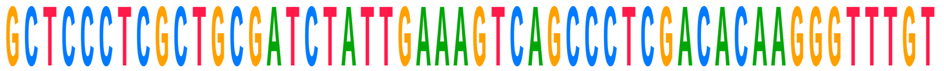

| | | |
|---|---|---|
| | GCTCCCTCGCTGCGATCTATTGAAAGTCAGCCCTCGACACAAGGGTTTGT- | |
| 1 | GCTCCCTCGCTGCGATCTATTGAAAGTCAGCCCTCGACACAAGGGTTTGTC | 5054 |
| 2 | GCTCCCTCGCTGCGATCTATTGAAAGTCAGCCCTCGACACAAGGGTTTGT- | 4803 |
| 3 | GCTCCCTCGCTGCGATCTATTGAAAGTCAGCCCTCGACACAAGGGTTTGT- | 4800 |
