## Supplementary Table S2 for "Neuronal RNA granules are ribosome complexes stalled at the pre-translocation state"

| **Gene ID or Protein ID** | **Identity** | **Gene ID or**  **Protein ID** | **Identity** |
| --- | --- | --- | --- |
| LOC100360573 | 98% to eS12 | M0RBS7 | 98% to ELAV-like 4 |
| RGD1559531 | 91% to eL21 | A0A0G2K8A9 | 90% to **LARP4** |
| RGD1562399 | 95% to uS5 | D3ZXI2 | Putative nucleic acid binding protein |
| D4A1Q0 | 98% to eL36 |  |  |
| A0A0G2JY70 | 91% to uS10 |  |  |
| Rpl30l1 | 92% to eL30 |  |  |
| D3ZFZ8 | 93% to eS24 |  |  |
| D3ZVY6 | 91% to eL31 |  |  |

Supplementary Table S2.
