## Supplementary Table S4 for "Neuronal RNA granules are ribosome complexes stalled at the pre-translocation state"

| **Rat CH-tissue** | | | **SH-SY5Y** | | |
| --- | --- | --- | --- | --- | --- |
| **Protein** | **Ribosomes per protein** | **Standard deviation** | **Protein** | **Ribosomes per protein** | **Standard deviation** |
| Caprin1 | 9.54 | 4.2 | ELAVL4 | 7.8 | 1.4 |
| Elavl4 | 25.69 | 19.6 | SERBP1 | 8.1 | 1.6 |
| G3bp2 | 8.36 | 1.8 | CAPRIN1 | 18.6 | 8.3 |
| Elavl3 | 12.96 | 2.7 | G3BP1 | 47.2 | 0.3 |
| G3bp1 | 24.26 | 13.6 | ELAVL3 | 53.6 | 6.0 |
| Serbp1 | 14.72 | 0.2 | G3BP2 | 74.8 | 11.7 |
| Stau2 | 17.60 | 3.0 | STAU2 | 61.0 | 11.8 |
| Fmr1 | 37.84 | 20.4 | FXR1 | 99.2 | 26.3 |
| Elavl2 | 13.88 | 6.2 | STAU1 | 91.2 | 55.0 |
| Stau1 | 59.85 | 27.0 | FMR1 | 117.2 | 34.3 |
| Fxr1 | 47.37 | 10.9 | USP10 | 212.7 | 46.3 |
| Fxr2 | 55.21 | 2.5 | ELAVL2 | 274.7 | 54.3 |
| Usp10 | 123.64 | 37.0 | FXR2 | 540.5 | 275.6 |

Supplementary Table S4. Relative abundance of RBPs in nRNAg purified from rat cortex-hippocampus tissue or human neuroblastoma SH-SY5Y. The relative abundance values were calculated based on the iBAQ intensity values listed in Supplementary Table S1. Ribosomal proteins average iBAQ intensity value was taken as reference and was assigned 1 equimole. The relative abundance of RBPs were calculated with respect to the ribosome. Calculations were done based on 2 biological replicates and standard deviations was calculated.
