## Supplementary Table S7 for "Neuronal RNA granules are ribosome complexes stalled at the pre-translocation state"

| **CH replicate 1** | **CH replicate 1** | **SH-SY5Y replicate 1** | **SH-SY5Y replicate 2** |
| --- | --- | --- | --- |
| L39 | S10 | L36al | L10 |
| L36 | S27 | S26 | L3l |
| L21 | S15 | S15 | L26l |
| L6 | S25 | L26l1 | L22l1 |
| S24 | S4y2 | S27l |  |
| L3l |  | S3 |  |
|  |  | L3l |  |
|  |  | L22l1 |  |

Supplementary Table S7.
