## Supplementary Table S8 for "Neuronal RNA granules are ribosome complexes stalled at the pre-translocation state"

| **r-protein**  **name** | **Mutated residue** |
| --- | --- |
| L3 (uL3) | E140D/D141T/S151N/V162I/V331I |
| L4 (uL4) | R143C/I144V/G160S/L169Q/I285M/D352K/A354L/A355E/Q362A |
| L5 (uL18) | K19R/T126N/ S230N/K259R |
| L6 (eL6) | M77L/P95T/K139R/A142S/I149V/V180A/I201V/N205K/K221P/I264V/I281M/A276S/V274Q |
| L7 (uL30) | I30V/Q39L/M41T/I130V/A173S/I175V/Y182F |
| L9 (uL6) | P130T/A190P |
| L13 (eL13) | V9I/R20Q/A23D/V90M/V157I/V168A |
| L14 (eL14) | V15I/Q44R/H69R/Q75K/Q78E/E96D/N125T/K132R |
| L17 (uL22) | Q54K |
| L18 (eL18) | V92I/Q93L/R115K/T106S/D129E/C134R |
| L18a (eL20) | S135G |
| L21 (eL21) | V75I |
| L26 (uL24) | H100R |
| L27a (uL15) | L38M/K59R/T95N/A97V/S139G |
| L28 (eL28) | I64M or M125V |
| L29 (eL29) | M65V/K79Q/E80A/I85M/V89S/D94S/Y98F/R107K/A108I/A110S/R111Y/I112M/L116R |
| L35 (uL29) | N96T/N101K |
| L35a (eL33) | S7C |
| L36a (eL42) | R38K |
| L37a (eL43) | K87E |
| SA (uS2) | A4G |
| S2 (uS5) | M61I |
| S3 (uS3) | S195T |
| S17 (eS17) | K45N and P134A |
| S19 (eS19) | M88R |
| S21 (eS21 | G25A/K41R/V42S/A54G |
| S27a (eS31) | A133G |

Supplementary Table S8. Ribosomal protein primary sequence differences between human and Rattus norvegicus that were modeled into structure. Listed are only the r-proteins and residues that exist in the structure. Old and new nomenclature of r-protein names are listed (Ban *et al*, 2014).

Ban N, Beckmann R, Cate JH, Dinman JD, Dragon F, Ellis SR, Lafontaine DL, Lindahl L, Liljas A, Lipton JM *et al* (2014) A new system for naming ribosomal proteins. *Curr Opin Struct Biol* 24: 165-169
