## Supplementary Table S9 for "Neuronal RNA granules are ribosome complexes stalled at the pre-translocation state"

**Reverse transcription was applied under the following conditions**

| Step | Temperature (◦C) | Time (min) |
| --- | --- | --- |
| Primer annealing | 25◦C | 7 min |
| Reverse Transcription | 40◦C | 15 min |
| Enzyme Inactivation | 85 ◦C | 5 min |

**Polymerase chain reaction (PCR) amplification conditions**

| Step | Temperature (◦C) | Time |
| --- | --- | --- |
| Initiation step (1x Cycle) | 98◦C | 40 sec |
| Denaturation (35x Cycles) | 98◦C | 10 sec |
| Primer Annealing (35x Cycles) | 50◦C | 15 sec |
| Extension (35x Cycles) | 72◦C | 50 sec |
| Extension (1x Cycle) | 72◦C | 5 min |

Supplementary Table S9. Reaction conditions for reverse transcription and subsequent PCR of rRNA purified from rat CH nRNAg.
